# Inferring hominin history with recurrent gene flow from single unphased genomes and a two-locus statistic

**DOI:** 10.64898/2026.04.11.717825

**Authors:** Nicholas W. Collier, Simon Gravel, Aaron P. Ragsdale

**Affiliations:** Department of Integrative Biology, University of Wisconsin–Madison, Madison, WI, USA; Department of Human Genetics, McGill University, Montreal, QC, Canada

**Keywords:** human evolution, demographic inference, population history, ancient DNA, archaic introgression, two-locus statistics

## Abstract

The emerging picture of hominin evolution is one of complexity, population structure, and gene flow, which recent genomic inference approaches have begun to resolve. Among these are methods based on two-locus statistics, which summarize information contained in genealogical correlations between linked loci. Although the inclusion of ancient samples could provide increased power to distinguish between competing models, these methods typically rely on large samples from present-day populations, and it remains challenging to apply them to ancient DNA (aDNA), which is sparsely sampled, unphased, and time-stratified. Here we develop an inference framework based on a set of multi-population two-locus statistics that are applicable to aDNA because they can be estimated from single unphased diploid genomes. We connect these statistics to an existing system of two-locus summaries and use them to model divergence and gene flow among populations represented by seven ancient hominin individuals and one contemporary human. We infer a demographic model with two episodes of gene flow from early anatomically modern humans (AMH) to Neanderthals and an introgression from an unsampled hominin lineage to Denisovan ancestors, broadly consistent with previous work. We also learn parameters of ancient Eurasian AMH population structure, reinforcing previous findings that early European farmers traced a large fraction of their ancestry to a lineage which split early from other non-African AMH and received little or no introgression from Neanderthals. Using both simulation and empirical data, we show that accurately estimating parameters associated with multiple gene flow episodes requires their joint inference due to their correlated effects on diversity.

## Introduction

Investigations of ancient DNA (aDNA) have revealed formerly unknown events in hominin evolution. It is now widely accepted many living humans carry a small fraction of genetic material inherited from the Neanderthal and Denisovan lineages (reviewed by Nielsen et al., 2017; Gokcumen, 2020; Bergström et al., 2021). Other studies have found evidence of gene flow from early anatomically modern humans (AMH) into Neanderthal populations (Posth et al., 2017; Rogers et al., 2020; Li et al., 2024). It has been hypothesized that unsampled and uncharacterized hominin lineages contributed ancestry to Neanderthals and Denisovans (Prüfer et al., 2014; Rogers et al., 2020) and to AMH (Durvasula and Sankararaman, 2020; Cousins et al., 2025; Rogers et al., 2026). Other work has used aDNA to investigate the worldwide dispersal of AMH and the ancestry of recent and contemporary human populations (e.g., Lazaridis et al., 2014). The identifiability and existence of further episodes of gene flow in hominin evolution remain open questions; likewise, their significance in explaining neutral and adaptive genetic variation in contemporary human populations.

The inference of demographic history using aDNA faces several inherent challenges (Orlando et al., 2021). Samples are scarce due to the paucity of suitable subfossil material; many samples cannot be sequenced to high depth, limiting the applicability of inference techniques which demand high-coverage data; aDNA has undergone substantial fragmentation and chemical damage, precluding haplotype phasing with long-read sequencing; and the lack of suitable reference panels and small sample sizes renders computational phasing challenging. Some model-based demographic inference techniques do not require phasing or large sample sizes and have been applied to aDNA, including the *F* -statistics (Patterson et al., 2012; Peter, 2016), which measure shared genetic drift, and summaries of derived allele frequencies (Kamm et al., 2020; Rogers et al., 2020). Other techniques model the genealogical information contained in multi-locus haplotypes: by estimating the distribution of coalescence times along a diploid genome, the pairwise sequentially Markovian coalescent (PSMC) model (Li and Durbin, 2011) infers the history of pairwise coalescence rates, which contains information about both population size history and population structure (Mazet et al., 2016). Still other methods exploit identical-by-descent (IBD) tract sharing, which reflects recent shared ancestry among samples and can be proxied by long-range linkage disequilibrium (LD) in unphased data (Fournier et al., 2023).

As tractable summaries of the statistical associations between pairs of linked loci, two-locus statistics are in a sense intermediate between one-locus summaries and PSMC-based approaches which model coalescence times of multi-locus haplotypes, but have yet to be applied broadly to aDNA. Two-locus statistics reflect properties of pairs of sampled gene genealogies (Hudson, 1985) and capture information about demographic history through its effects on the covariance between pairwise coalescence times (McVean, 2002). Demographic inference using two-locus statistics often makes use of the system of Hill and Robertson (1968), which models the evolution of 𝔼[*D*^2^] (the variance of the covariance measure of LD, *D*) and related summaries. These statistics are sensitive to historical population size changes (Rogers, 2014), and Ragsdale and Gravel (2019) extended them to develop a multi-population framework that allows for the joint inference of population splits, size changes, and gene flow.

The application of two-locus statistics to aDNA inference has attracted theoretical attention from Biddanda et al. (2022), who showed how the expected correlation of pairwise divergences and the between-population analog of 𝔼[*D*^2^] are affected by time-stratified sampling. However, measuring within-population two-locus statistics from aDNA can be challenging. We are often limited to a single diploid (or even pseudo-haploid) sample from each population of interest, but the standard estimation of two-locus statistics requires larger sample sizes – for example, estimating *D*^2^ requires four haplotypes, or four diploid samples if sequences are unphased (Ragsdale and Gravel, 2019, 2020). This prevents the use of high-order statistics like 𝔼[*D*^2^] with single diploid samples, and motivates the development of methods that utilize statistics which can capture allele sharing and genealogical covariance information from such samples.

In this work, we develop a demographic inference approach based on *H*_2_, a family of multi-population two-locus statistics that can be estimated from one unphased diploid per population. 𝔼 [*H*_2_] statistics are linear combinations of two-locus summaries from the well-studied Hill-Robertson (HR) system (Hill and Robertson, 1968) and its multi-population extension (Ragsdale and Gravel, 2019). The decay of 𝔼[*H*_2_] across recombination distances summarizes joint distributions of pairwise coalescence times and collects information about their marginal expectations and covariances, which can be used to learn demographic history. This property allows 𝔼[*H*_2_] to discriminate between models with and without ancestral population structure which may be difficult to distinguish using one-locus diversity patterns. Motivated by this potential boon to model identifiability, we fit a richly parameterized model that describes the ancestry of populations represented by eight individuals (a contemporary human, three ancient Eurasian AMH, three Neanderthals, and a Denisovan). Using a simulation-based method to perform formal model choice, we test the statistical support for gene flow events previously inferred with other methodologies. We jointly model these features and estimate key parameters, including population divergence times, rates of gene flow, and introgression proportions.

## Results and Discussion

### Estimating *H*_*2*_ from single unphased diploid genomes

Heterozygosity *H*_*x*_ is the probability of observing a difference in allelic state between the two genome copies of a diploid at locus *x*. Assuming random mating, *H*_*x*_ is equivalently the probability of observing a difference between two randomly sampled haploid genome copies; if *x* is biallelic with allele frequencies *p* and 1 − *p, H*_*x*_ = 2*p*(1 − *p*). Heterozygosity can be estimated from a single diploid genome using *Ĥ*_*x*_ = **1**_het(*x*)_. As we sample increasingly many realizations of the evolutionary process (multiple loci), the average of this estimate is expected to converge to its expectation 𝔼[*H*]. Analogously, we define *H*_2_ as the probability of observing differences between two sampled genome copies at two loci. Consider two biallelic loci labeled left (*L*) and right (*R*), where alleles *a, A* and *b, B* segregate. Four two-locus haplotypes are possible (*AB, Ab, aB, ab*), with frequencies (*f*_*AB*_, *f*_*Ab*_, *f*_*aB*_, *f*_*ab*_). Assuming random mating and the availability of phasing information,

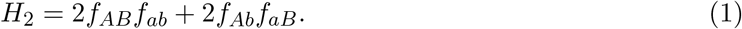

If we consider the two genome copies of a diploid, then *H*_2_ = *H*_*L*_*H*_*R*_, regardless of whether there is phasing information. The one-diploid estimator is *Ĥ*_2_ = *Ĥ*_*L*_*Ĥ*_*R*_.

To extend to two diploids indexed by *i* and *j*, and assuming phasing information, we define a *phased* between-diploid statistic by sampling one haplotype from each diploid.

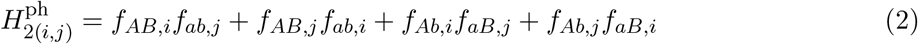

In this work, where each diploid sample represents a population of interest, it is equivalent to let *i, j* index populations and to call 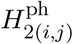 a between-population statistic. Although we define the between-population statistic using haplotype probabilities, throughout this work we use an *unphased* statistic 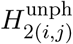, which can be estimated from genotype data (in what follows, we write *H*_2(*i,j*)_ to represent the unphased statistic and label phasing only when there is a need to explicitly discuss both statistics). To estimate the unphased statistic, we average over all possible phasings, assuming an equal probability of each (see Supplementary Section 2.3). The lack of phasing information causes 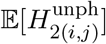 to diverge from 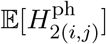 at small recombination fractions. Each *H*_2_ statistic (within-diploid, phased between-diploid, unphased between-diploid) is a linear combination of summaries from the HR system, so we can compare either phased or unphased data to model predictions (see Appendix A).

### 𝔼[*H*_*2*_] captures genealogical covariance

𝔼[*H*_2_] summarizes the joint distribution of pairwise coalescence times. Let *T*_*L*_ and *T*_*R*_ be pairwise coalescence times at left and right loci. In Appendix B, we derive the following relation between 𝔼[*H*_2_] and the covariance of pairwise coalescence times, or genealogical covariance:

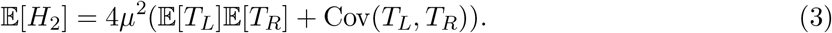

Equation 3 relies on assumptions about the mutation process, but holds for any population history. Under demographic equilibrium, measuring time in coalescent units *τ*_*L*_ = *T*_*L*_*/*2*N*_*e*_, we have

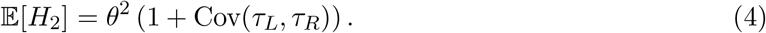

The genealogical covariance at equilibrium is shown in Figure 1A. At equilibrium, Var(*τ*_*L*_) = 1, so we can replace Cov(*τ*_*L*_, *τ*_*R*_) with corr(*τ*_*L*_, *τ*_*R*_) in Equation 4. We see that 𝔼[*H*_2_] decays as a function of recombination distance *ρ* = 4*N*_*e*_*r*, going from 2*θ*^2^ at *ρ* = 0, where corr(*τ*_*L*_, *τ*_*R*_) = 1 because loci always share the same genealogical history, to *θ*^2^ at long distances where corr(*τ*_*L*_, *τ*_*R*_) is negligible (Figure 1B). Demographic events perturb the joint distribution of coalescence times, altering the patterns of Cov(*T*_*L*_, *T*_*R*_), and in turn 𝔼[*H*_2_], decay (Figure 1C-F; see Appendix C for further discussion). 𝔼[*H*_2_] is informative about events in the distant past because it evolves slowly; under the infinite-sites model, the appearance of polymorphism at two formerly invariant loci requires one mutation to happen at each locus, an event with a long waiting time.

**Figure 1.**
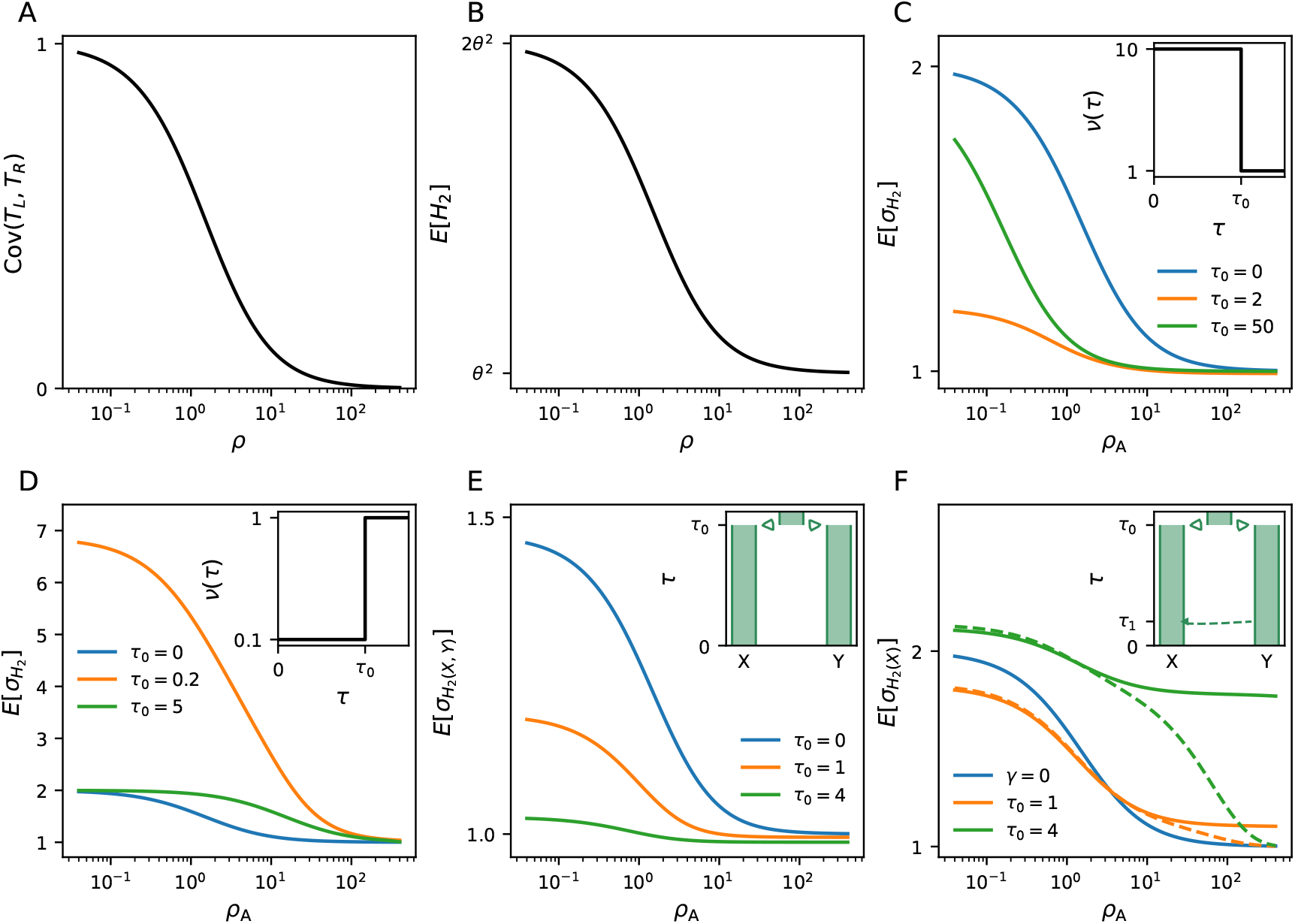
*H*_*2*_ under non-equilibrium demography. (A) Equilibrium genealogical covariance plotted against the population recombination rate *ρ* = 4*N*_*e*_*r*. (B) Equilibrium 𝔼[*H*_2_] in terms of the population mutation rate *θ* = 4*N*_*e*_*µ*. (C) The diversity-normalized statistic 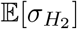 (see Appendix C) is depressed following population size expansion. The time since expansion *τ*_0_ is given in genetic units (2*N*_A_); the relative size history *ν*(*τ*) is displayed in the inset panel. (D) 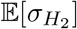 is elevated following a population contraction (inset panel). (E) The between-population statistic 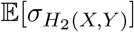 decreases following divergence with isolation for *τ*_0_ genetic units (inset panel). (F) 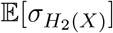 is distorted following introgression (inset panel; the introgression proportion is *γ* = 0.1). Solid lines show the expectation immediately following introgression (*τ*_1_ = 0), and broken curves show the expectations at *τ*_1_ = 0.025.

### Ancestral population structure is theoretically identifiable

Several recent population genetic studies have inferred models of ancestral structure marked by introgression from an isolated deme (Durvasula and Sankararaman, 2020; Fan et al., 2023; Cousins et al., 2025) or persistent contact among differentiated demes (Ragsdale et al., 2023) in early human evolution. Motivated by this work, we show that 𝔼[*H*_2_] contains information that can distinguish between models with structured and panmictic ancestral populations, even when they have the same marginal coalescence time distribution. As an example, consider the non-stationary ancestral structure model shown in Figure 2A. To find a panmictic model with a matching marginal distribution, we compute the instantaneous coalescence rate (ICR) profile *λ*(*t*) of the structured model. We invert this rate to obtain a panmictic population size function *N* (*t*) = 1*/*2*λ*(*t*) (Figure 2B) with an equal ICR profile (Figure 2D) (Mazet et al., 2016). The 𝔼[*H*_2_] curves generated by panmictic and structured models of this form are distinguishable (Figure 2C) when *M* = 4*N*_*e*_*m* is small, because ancestral structure tends to isolate ancestral gametes in different demes, inducing more genealogical covariance than expected for a panmictic model (see Section C). Following the end of the structured epoch, excess covariance decays rapidly when *ρ* is large, but persists for longer at small recombination fractions (Figure 1F). For both structured and panmictic models, the curve is depressed below 2𝔼[*H*]^2^ at low recombination fractions, reflecting an overall growth-like history (see Figure 1C). The distinguishability of panmictic and structured model expectations depends on the particular structured history considered and its parameters, and increases with longer divergence times, lower gene flow, and larger minor ancestry proportion (Figures S5, S6).

**Figure 2.**
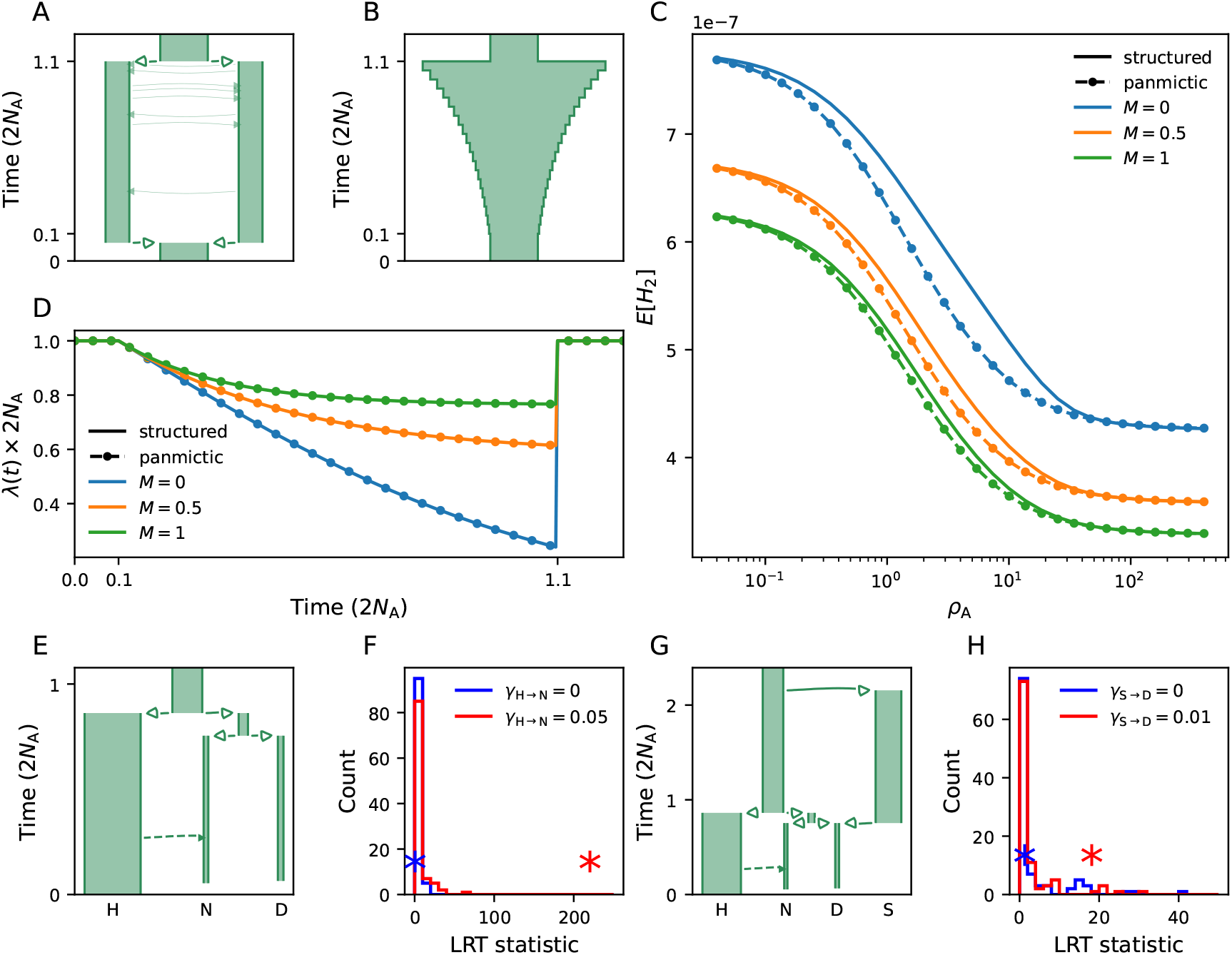
Indentifiability of structure and gene flow. (A) A model of ancestral population structure, where *N*_*A*_ is the ancestral size, demes have effective sizes *N*_*e*_ = *N*_*A*_*/*2, and the betweendeme migration rate is *M* = 4*N*_*e*_*m*. Demes from the structured epoch contribute equally (*γ* = 0.5) to the sampled population. (B) A panmictic model with *λ*(*t*) profile matching the ancestral structure model. The time discretization plotted is coarser than the one used to compute *λ*(*t*) and 𝔼[*H*_2_]. (C) When *M* is small, 𝔼[*H*_2_] curves differ for structured and panmictic models with identical *λ*(*t*) profiles. (D) Scaled ICR profiles 2*N*_A_*λ*(*t*) for models plotted in panel C, which are equivalent between the structure (A) and panmictic (B) models. (E) Topology of the H→N analog model. (F) Simulated BLRT for *γ*_H→N_ ≥ 0 on data generated with models where *γ*_H→N_ = 0 (*p*_obs_ = 0.44) and *γ*_H→N_ = 0.05 (*p*_obs_ *<* 0.01). Histograms display empirical null distributions and asterisks show observed LRT scores. The replicates shown had median *p*_obs_ among 10 independent realizations. Results for all replicates are given in Table S3. (G) Topology of the S→D analog model. (H) Simulated BLRT for *γ*_S→D_ ≥ 0 on data where *γ*_S→D_ = 0 (*p*_obs_ = 0.32) and *γ*_S→D_ = 0.01 (*p*_obs_ = 0.06).

### A method of formal model choice

To rigorously determine which historical features are supported by data, we need a criterion for model choice. The likelihood-ratio test (LRT) is a standard method for comparing the goodness of fit of a constrained model to that of a nested, less-constrained model. Because the LRT statistic for composite likelihoods (see Inference and model choice) has an unknown null distribution, we cannot use the generic LRT. Instead, we utilize the bootstrap LRT (BLRT), which uses simulation to construct an empirical null distribution and calculate observed *p*-values (*p*_obs_) (see Supplementary Section 3.5). We carried out a simulated analysis of BLRT power on two families of demographic models that are analogous to proposed histories for AMH, Neanderthals and Denisovans (H, N, and D, respectively). One model family was designed around a test for introgression from H to N at 250 kya (thousand years ago) (Figure 2E). The other family incorporated an additional introgression event from the unsampled population S to D at 700 kya (Figure 2G). We expected 𝔼[*H*_2_] to be sensitive to these ancient events due to its slow evolution. For each family of models, we performed simulations with several different introgression proportions *γ*, including *γ* = 0, then applied the BLRT to test the hypothesis that *γ* ≥ 0. (see Supplementary Section 4.5). We display BLRT distributions and LRT statistics for the trials with median *p*_obs_ at different *γ* in Figure 2F, H, and summarize all replicates in Table S2. Realized *p*_obs_-values suggest that we have moderate power to detect introgression events when the proportions are close to their proposed magnitudes (∼5% for H→N, ∼1% for S→D), and that power increases with proportion. We observed no false positive tests (rejections of the null model when true *γ* = 0).

### Inferring models of hominin introgression

#### General approach

Here, we test and refine models of divergence and gene flow between Neanderthals, Denisovans, AMH, and unsampled lineages. We refer to the ancestral populations of sampled AMH, Neanderthals, and Denisovans as simply *AMH, Neanderthals*, and *Denisovans*, without regard for the morphological or taxonomic implications of these terms (for a perspective on informal nomenclature in hominin evolution, see Hawks, 2025). Rather than initially including all representative samples (Table 2), we began by testing hypotheses on subsets of samples, as simpler models are more tractable to fit to data. We focused on three hypothesized gene flow events: (1) early AMH introgression to Neanderthals, (2) “ghost” or so-called “Superarchaic” introgression to Denisovans, and (3) population structure among ancient Eurasian AMH. Throughout, we fit models to two *H*_2_ datasets, each estimated using a human recombination map inferred with distinct methods and data, which we labeled Bhérer (Bhérer et al., 2017) and Zhou (Zhou et al., 2020) (see Obtaining data). Unless otherwise noted, we used a fixed mutation rate of 1.3 *×* 10^−8^ per bp per generation and a generation time of 29 years (see Supplementary Section 3.7). We report likelihoods and *p*_obs_-values for the Bhérer dataset in the main text and from both datasets in figure captions and Table S2. We display maximum-likelihood estimates (MLE) of parameters and 95% bootstrap confidence intervals (CIs) from the Bhérer dataset throughout the main text and in all supplementary tables except Table S7, where we compare focal models inferred with the two datasets.

#### Early AMH contributed ancestry to Neanderthals

To test support for gene flow from early AMH to Neanderthals, we fit a model to data from a contemporary African (Yoruba), an early European hunter-gatherer (Loschbour), the Vindija, Altai, and Chagyrskaya Neanderthals, and the Denisova-3 Denisovan (Denisova). Our preliminary work supported a Neanderthal-Denisovan clade, with a roughly 50 ky (thousand year) existence (from 779 to 726 kya) for the Neanderthal-Denisovan common ancestor (ND), and a recent (123 kya) divergence between sampled Neanderthal lineages. The first of these findings stands in contrast to a recent phylogenetic analysis of hominin skull morphology, which inferred a Denisovan-AMH clade (Feng et al., 2025). We did not model migration between sampled Neanderthal lineages, because their relatively brief independent existences (between 4 and 15.5 ky) might make migration rates difficult to identify. We did however model symmetric migration between the ancestral Neanderthal (AN) and Altai demes and Denisova – gene flow between these populations occurred intermittently in central Asia, as evidenced by inferred introgressed tracts in sequenced genomes from each lineage (Peter, 2020; Peyregne et al., 2025). We inferred a Neanderthal-Denisovan migration rate of 2.58 *×* 10^−6^ (0.407 – 5.43 *×* 10^−6^) per gamete per generation. Although this rate is small (*N*_*e*_*m* ≪ 1), migration continues for a long enough time (from 726 kya to 115 kya) to appreciably impact expectations. Tracing the ancestry of a single allele copy from Denisova back in time, the probability that the copy migrated from the Neanderthal deme (ignoring the chance that an allele copy migrated more than once) is 5.29% (calculated with 1 − (1 − *m*)^*t*^, where *t* is the time in generations).

After modeling various combinations of introgression events, we found the strongest support for a model with one AMH introgression to AN 250 kya and one to the western Neanderthal (WN) lineage, which is ancestral to Vindija and Chagyrskaya, 110 kya (Figure 3A). We fixed introgression time parameters at or near the values inferred by Li et al. (2024) to prevent runaway behavior; when we estimated them as free parameters, we observed a strong positive correlation between introgression proportion and time across bootstrap samples, which suggests confounding (the effect is seen in simulated data, Figure S23 and see Supplementary Section 4.4). A model with only AMH→AN is overwhelmingly supported against a model with no AMH contribution (Δ*LL* = 69, BLRT *p*_obs_ *<* 0.002; see Figure 3B). Compared to this one-pulse model, a two-pulse model with the later event AMH→WN also finds strong support (Δ*LL* = 78, BLRT *p*_obs_ *<* 0.002; see Figure 3C). The qualitative improvement in the fit of the model is mostly seen in higher 𝔼[*H*_2_(Neanderthal)] at intermediate recombination distances and in subtle changes to between-population curves (Figure 3D– I). There is noticeable improvement in the fit for the Denisova curve (Figure 3G), likely because the null model is more misspecified, biasing parameter estimates.

**Figure 3.**
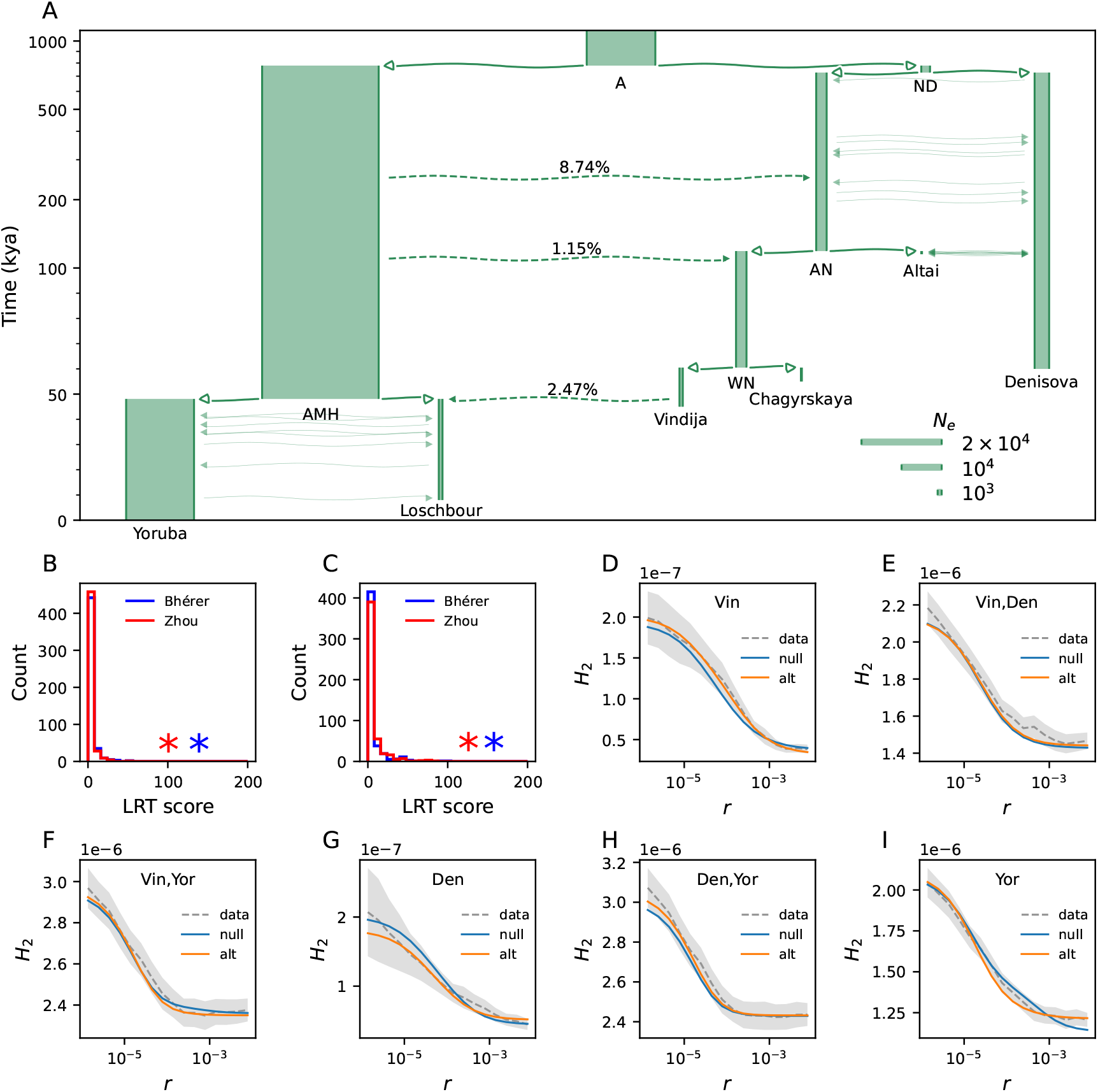
Gene flow between human, Neanderthal and Denisovan ancestral lineages. (A) The maximum-likelihood (ML) AMH–Neanderthal introgression model. The time scale is linear below 100 kya and logarithmic above; broken one-headed arrows denote instantaneous gene flow events; solid double-headed arrows denote continuous gene flow. The density of continuous gene flow arrows is the same for each pairwise migration and is unrelated to the inferred migration rate (see Table S4 for inferred parameters). (B) BLRT for AMH→AN pulse (both datasets *p*_obs_ *<* 0.002). (C) BLRT for AMH→WN, nested in the alternate model from the test in panel B (both datasets *p*_obs_ *<* 0.002). (D-I) Comparisons between data and expectations under models without AMH-to-Neanderthal introgression (null) and with AMH→AN, AMH→WN pulses (alt) for a subset of *H*_2_ statistics (see Figure S7 for all fitted curves). Shaded areas are 95% bootstrap CI.

We included introgression from a relative of WN to early Eurasian AMH in each of the described models. We fixed the time of this event to 48 kya, based on the inferences of Iasi et al. (2024). As our inferences changed little when we approximated the event as a direct instantaneous contribution from Vindija to Loschbour, we adopted this simplification. In the null model without AMH-to-Neanderthal gene flow, we inferred an unreasonably large Vindija-to-Loschbour introgression proportion of 8.69% (6.24 – 11.3%). Jointly modeling both AMH-to-Neanderthal pulses resulted in a smaller introgression proportion (2.47%, 1.29 – 3.78%), which stands in better agreement with earlier work (e.g., 1.5 – 2.1%, Prüfer et al., 2014).

We inferred population size decreases in each sampled Neanderthal deme (Figure 3A), consistent with IICR curves inferred with the PSMC, which decline precipitously preceding the lifetimes of sampled Neanderthal and Denisovan individuals (Prüfer et al., 2017). This has been interpreted as evidence of population size collapse, under the assumption that populations are panmictic (Prüfer et al., 2014). However, this finding is also compatible with other models: Rogers (2024) showed that the inferred IICR curves are plausibly explained by geographic population structure. With spatial structure, recent ancestors are expected to live in closer proximity, and to therefore have a higher probability of sharing parents, than ancient ancestors. Strong structure therefore causes recent coalescence rates to be larger than ancient ones, a pattern which is interpreted as a small recent effective size in a panmictic model (see also Mazet et al. 2016).

#### Denisovans received gene flow from an unsampled lineage

We next examined support for gene flow to Denisovans from a ghost lineage that has been called “Superarchaic” by several authors and speculatively identified with an Asian *Homo erectus* population (Prüfer et al., 2014; Rogers et al., 2020). We fixed the effective size of the Superarchaic lineage (S) to 20,000 and its split time from the ancestral population to 2,000 kya, which are similar to the values inferred by Rogers et al. (2020). The divergence time was also motivated by the estimated occupancy of *H. erectus* populations in Eurasia (from 1,800 kya in Indonesia and 1,700 kya in the Republic of Georgia, Antón, 2003) and is close to the MLE when the split time is fitted as a free parameter (1,940 kya). We justified fixing the population size of S with the observation that changing the effective size of a ghost lineage which makes a small ancestry contribution to a sampled lineage has a negligible effect on 𝔼[*H*_2_] (Figure S3). This is because the probability that multiple gametes with ancestral allele copies are inherited from the ghost lineage is small (*γ*^2^ for two ancestral gametes and introgression proportion *γ*), so the coalescence rate in that lineage has little impact on model expectations and its effective size would be poorly constrained as a free parameter. We incorporated both AMH-to-Neanderthal introgression episodes inferred above into our primary test for Superarchaic introgression (Figure 4A).

**Figure 4.**
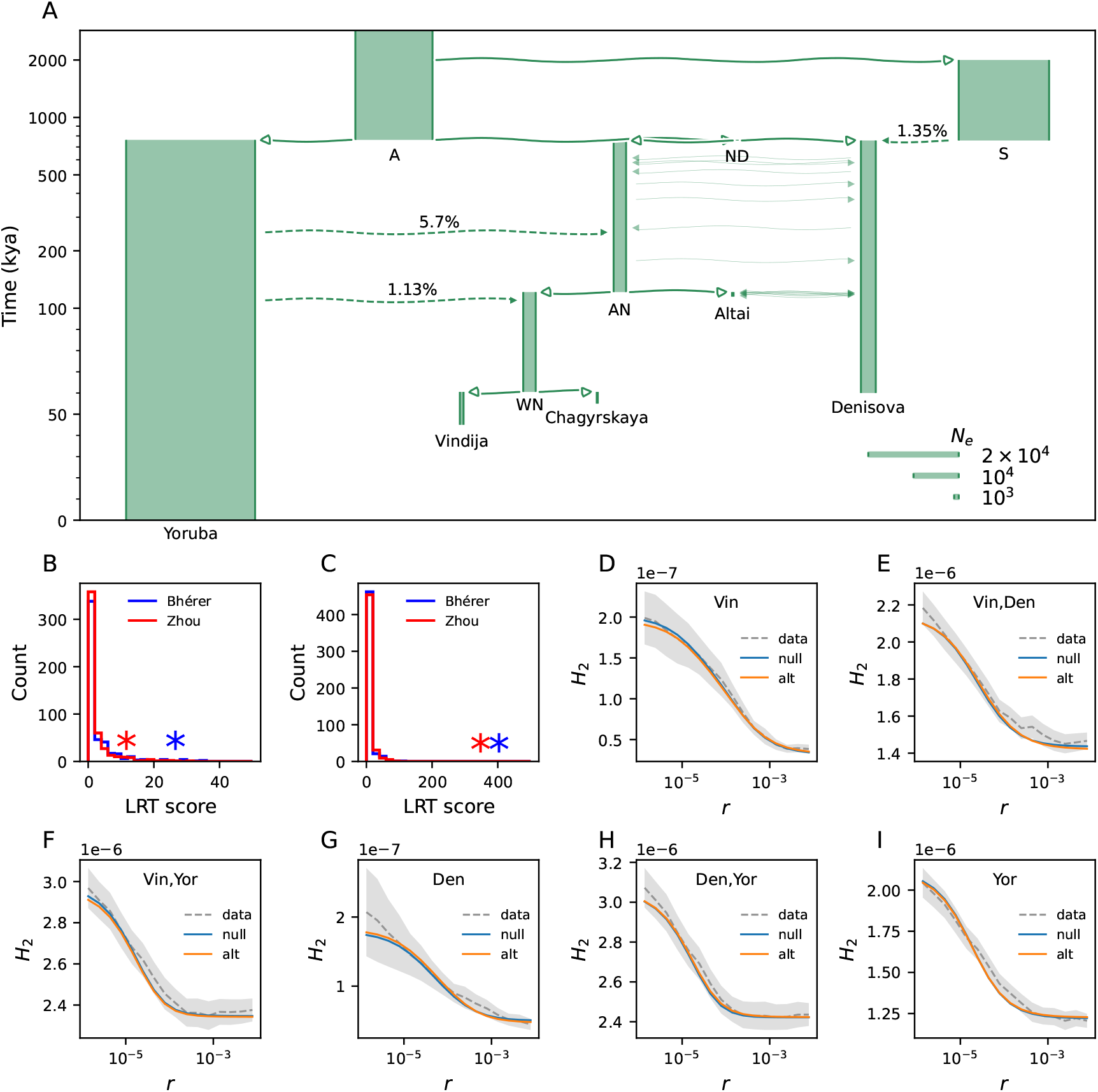
Neanderthal and Denisovan history with Superarchaic introgression. (A) The Superarchaic-to-Denisovan introgression model (see Figure 3 caption for model-plotting conventions and Table S5 for inferred parameters). (B) BLRT for S→Denisova pulse (Bhérer *p*_obs_ = 0.018, Zhou *p*_obs_ = 0.052). (C) BLRT for S→Denisova, *not* nested in the AMH-to-Neanderthal introgression model (both datasets *p*_obs_ *<* 0.002). (D–I) Comparisons between data and expectations under models without (null) and with (alt) S→Denisova introgression (nested in the AMH-to-Neanderthal model) for a subset of *H*_2_ statistics (see Figure S8 for all fitted curves). Shaded areas are 95% bootstrap CI.

With this parameterization, we observed marginal significance in a test for Superarchaic-to-Denisovan introgression (Δ*LL* = 13, BLRT *p*_obs_ = 0.018; see Figure 4B). The qualitative improvement of the fit to observed *H*_2_ is subtle, visible mostly as a depression of 𝔼[*H*_2_(Denisova, Neanderthal)] curves at small *r* (Figure 4D–I). We infer a Superarchaic ancestry contribution of 1.35% (0.27 – 2.3%), which lies at the lower end of previously estimated values (e.g., 0.5 – 8%, 1.67 – 2.65% from Prüfer et al., 2014; Rogers et al., 2020, respectively). Incorporating Superarchaic-to-Denisovan gene flow reduces the estimated AMH→AN introgression proportion from 8.82% (7.28 – 11.4%) to 5.70% (3.69 – 7.60%), suggesting that the two events are confounded to some extent. They appear to explain similar features in the data by elevating 𝔼[*H*_2_(Yoruba, Denisovan)] relative to 𝔼[*H*_2_(Yoruba, Neanderthal)]. In a test for the presence of Superarchaic introgression where AMH- to-Neanderthal introgressions are not jointly modeled, the null model was overwhelmingly rejected (Δ*LL* = 201, BLRT *p*_obs_ *<* 0.002; see Figure 4C) and we inferred a larger Superarchaic contribution (3.19%). These statistical tests support the hypothesis that Superarchaic-to-Denisovan and reciprocal AMH–Neanderthal gene flow both shaped the genetic relationship between hominin lineages (Prüfer et al., 2014; Rogers et al., 2020).

#### A Neolithic European derived ancestry from a basal Eurasian lineage

We then turned to modeling AMH lineages, represented by an early Eurasian (Ust’Ishim), a European hunter-gatherer (Loschbour), and a European farmer (Stuttgart), in greater detail. Most contemporary human populations outside Africa are thought to inherit a small fraction (∼2%) of Neanderthal ancestry (Prüfer et al., 2014). Recent work has supported the hypothesis that living non-Africans trace their Neanderthal ancestry to a single shared episode of gene flow which occurred 43.5 – 50.5 kya, though early Eurasian AMH (including the Ust’Ishim man) may have derived unique Neanderthal ancestry from other sources (Iasi et al., 2024). We found that explicitly modeling separate Neanderthal contributions made little improvement to the fit of our models, so we assumed a single pulse of Neanderthal-to-OOA (out-of-Africa AMH) introgression as an approximation. Based on sampling times, preliminary results, and the admixture graphs inferred by Lazaridis et al. (2014), we modeled the Ust’Ishim lineage (estimated age 45 ky) as splitting first from OOA, with Loschbour and Stuttgart lineages diverging later and remaining isolated. This model underestimated *H*_2_(Stuttgart, Denisova) and *H*_2_(Stuttgart, Neanderthal) (Figure 5E), which is analogous to underestimating the divergence between these samples. This large residual was resolved when we attributed a fraction of Stuttgart ancestry to a basal Eurasian (BE) deme which diverged from the OOA deme before Neanderthal-to-OOA introgression, reducing the amount of Neanderthal ancestry expected for the Stuttgart lineage (Figure 5A).

**Figure 5.**
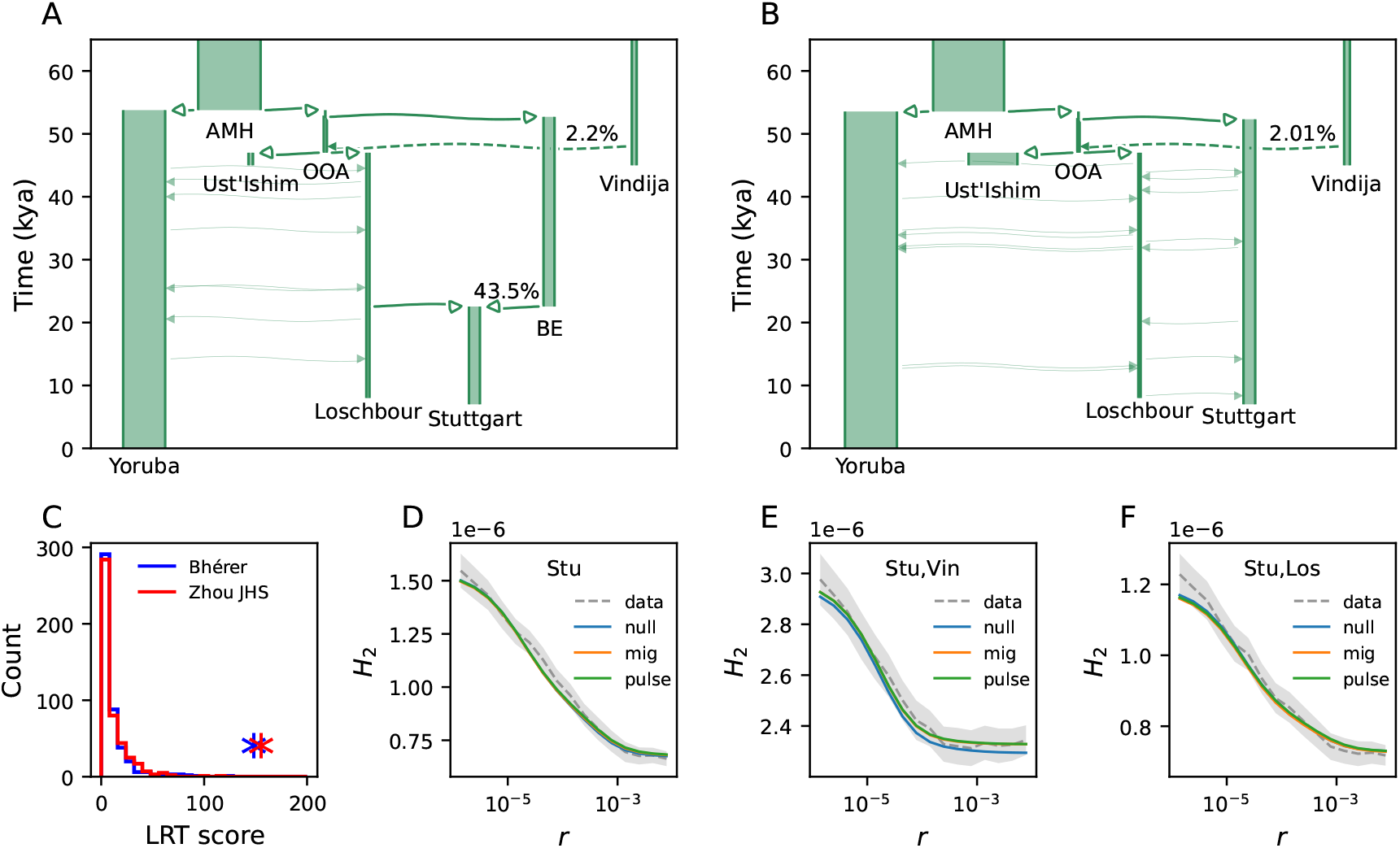
Population structure in early European history. (A) The Loschbour–Stuttgart admixture model, where the Stuttgart lineage is formed by admixture from the Loschbour and basal Eurasian (BE) demes, from 60 kya. (see Figure 3 caption for model-plotting conventions and Table S6 for inferred parameters). (B) The Loschbour–Stuttgart migration model, from 60 kya. (C) BLRT for Loschbour-BE admixture (both datasets *p*_obs_ *<* 0.002). (D-F) Comparison between data and expectations for Loschbour–Stuttgart isolation (null), migration (mig), and admixture (pulse) models for a subset of *H*_2_ statistics (see Figure S9 for all fitted curves). Shaded areas are 95% bootstrap CI.

This model agrees with the results of Lazaridis et al. (2014), who estimated that the Stuttgart individual derived ∼44% of her ancestry from the BE lineage, which received little to no Neanderthal introgression (Lazaridis et al., 2016). We inferred a BE ancestry proportion of 56.5% (30.7 – 70.3%) for Stuttgart and rejected the null hypothesis that the Stuttgart lineage derived *entirely* from BE, without admixture from more recent ancestors of Loschbour (Δ*LL* = 73, BLRT *p*_obs_ *<* 0.002; see Figure 5C). In addition to this admixture model, we fit a model where Stuttgart and Loschbour demes exchange migrants continuously (Figure 5B) and compared it to the same isolation null model (Δ*LL* = 64; no BLRT conducted). These models produced similar expected curves (Figure 5D–F) and parameter estimates (Table S6) – for instance, we inferred Yoruba–OOA separation times of 53.8 (51.1 – 57.8) and 53.6 (49.8 – 56.7) kya under admixture and migration models respectively (cf. 50 – 60 kya as reviewed by Bergström et al., 2021). In what follows, we use the admixture model due to its meaningful correspondence with earlier work (e.g., Lazaridis et al., 2014).

#### A joint model of Eurasian hominin gene flow

We fit a focal model which described eight samples and jointly incorporated well-supported features (Figure 6). Although this model has 28 free parameters, we were able to obtain convergence through successive rounds of optimization (see Supplementary Section 3.3). We found that the Neanderthal–Denisovan clade diverged from AMH 798 kya (748 – 827 kya), which is somewhat older than the 737 kya estimate from Rogers et al. (2020), but consistent with Peyregne et al. (2025)’s estimate of 694 – 825 kya. Work by Gómez-Robles (2019) on the rate of hominin dental evolution also supports a relatively ancient divergence (*>* 800 kya). We inferred a Neanderthal–Denisovan divergence time of 688 kya (639 – 734 kya), which is considerably older than the 504 – 585 kya interval estimated by Peyregne et al. (2025); this may be partially because we modeled migration between Denisovan and Neanderthal populations. Our model also recapitulated the recent divergence of sampled Neanderthal lineages, with recent reductions in effective population size, inferred by Prüfer et al. (2017) and Mafessoni et al. (2020) (but see Rogers, 2024).

**Figure 6.**
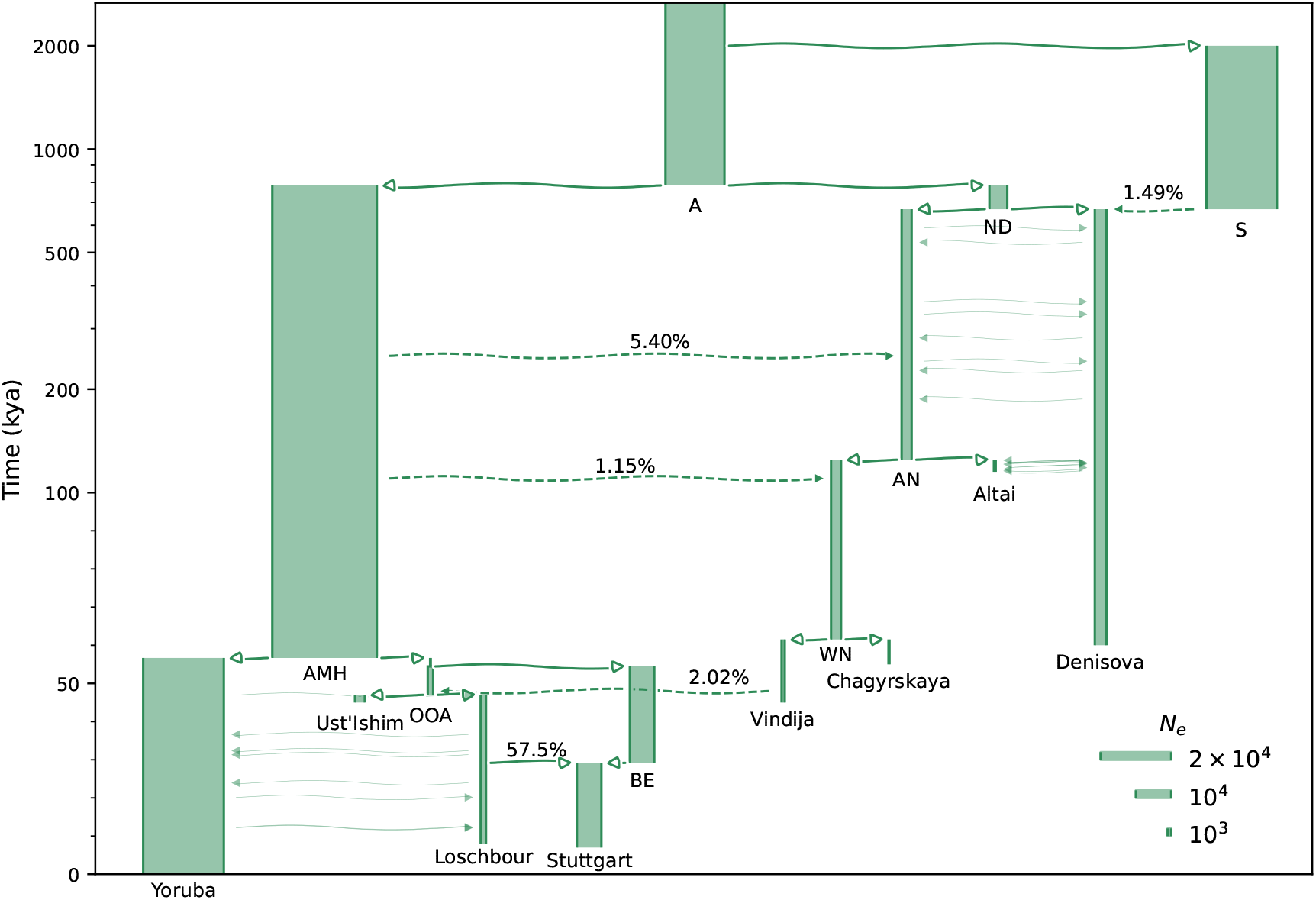
Early hominin history in Eurasia with recurrent gene flow. The full model combines and re-infers all statistically supported parameters from the subset models above. See Figure 3 caption for model-plotting conventions and Table 1 for a summary of key parameters.

Although the model provided a reasonable fit to data (Figures S10, S11), we observed a consistent residual in the domain *r* = 10^−5^ to *r* = 10^−4^, which was reproduced in datasets from both recombination maps (Figure S12). While this pattern could be artefactual, perhaps relating to some property of the recombination maps used to estimate *H*_2_, it may be partially explained by unmodeled ancestral population structure in the AMH lineage. This would be consistent with the properties of 𝔼[*H*_2_] shown in Figure 2C, D, which suggest that fitting a model with a panmictic ancestral population to data generated by a structured history may lead to inferences which underestimate 𝔼[*H*_2_] at small recombination fractions. Because the inferred model has long epochs with piecewise-constant population sizes, we don’t expect residuals to correspond exactly with those seen in this theoretical demonstration. This interpretation is complicated by the presence of the residual in observations which do not include AMH (e.g., *H*_2_(Denisovan, Neanderthal)). As this was not the focus of this work, our sample of AMH lineages was likely too sparse and affected by recent bottlenecks to meaningfully resolve any features of human ancestral population structure.

**Table 1:** Inferred parameters from full model. Maximum-likelihood estimates and confidence intervals of key focal model parameters (see Table S7 for all estimates).

| Parameter | Description | MLE | 95% CI |
| --- | --- | --- | --- |
| $T_{\text{ND-AMH}}$ (ky) | AMH–ND split time | 798 | 748 – 827 |
| $T_{\text{AN-Den}}$ (ky) | AN–Denisova split time | 688 | 639 – 734 |
| $T_{\text{WN-Alt}}$ (ky) | WN–Altai split time | 123 | 117 – 137 |
| $T_{\text{Cha-Vin}}$ (ky) | Chagyrskaya–Vindija split time | 60.5 | 57.3 – 67.7 |
| $T_{\text{Yor-OOA}}$ (ky) | Yoruba–OOA split time | 56.9 | 53.6 – 60 |
| $T_{\text{OOA-BE}}$ (ky) | OOA–BE split time | 54.7 | 49.7 – 57.6 |
| $T_{\text{Los-Stu}}$ (ky) | Loschbour→Stuttgart admixture time | 29.4 | 13.2 – 35.9 |
| $N_A$ | Ancestral $N_e$ | 16500 | 15900 – 17300 |
| $m_{\text{N-Den}}$ ($\times 10^{-5}$) | Denisova–AN, Denisova–Altai migration rate | 0.393 | 0.0866 – 0.685 |
| $m_{\text{Yor-Los}}$ ($\times 10^{-5}$) | Yoruba–Loschbour migration rate | 6.24 | 2.22 – 8.98 |
| $\gamma_{\text{S} \rightarrow \text{Den}}$ | S→Denisova pulse proportion | 0.0149 | 0.00748 – 0.0265 |
| $\gamma_{\text{AMH} \rightarrow \text{AN}}$ | AMH→AN pulse proportion | 0.054 | 0.0282 – 0.0702 |
| $\gamma_{\text{AMH} \rightarrow \text{WN}}$ | AMH→WN pulse proportion | 0.0115 | 0.00651 – 0.0166 |
| $\gamma_{\text{Vin} \rightarrow \text{OOA}}$ | Vindija→OOA pulse proportion | 0.0202 | 0.012 – 0.0274 |
| $\gamma_{\text{Los} \rightarrow \text{Stu}}$ | Loschbour→Stuttgart admixture proportion | 0.575 | 0.333 – 0.752 |

**Table 2:** Ancient samples considered in this work.References and methodologies for sample ages are given in Supplementary Section 3.6 and Table S1.

| Label | Est. age (ky) | Location | Reference |
| --- | --- | --- | --- |
| Altai Neanderthal | 115 | Denisova Cave, Russia | Prufer et al. (2014) |
| Chagyrskaya Neanderthal | 55 | Chagyrskaya Cave, Russia | Mafessoni et al. (2020) |
| Vindija Neanderthal | 45 | Vindija Cave, Croatia | Prufer et al. (2017) |
| Denisova-3 Denisovan | 60 | Denisova Cave, Russia | Meyer (2012) |
| Ust’Ishim AMH | 45 | Omsk Oblast, Russia | Fu et al. (2014) |
| Loschbour AMH | 8 | Mullerthal, Luxembourg | Lazaridis et al. (2014) |
| Stuttgart AMH | 7 | Stuttgart, Germany | Lazaridis et al. (2014) |

We predicted one-locus diversity statistics (*H, F*_2_) for the focal model and compared them to genome-wide empirical averages as an informal validation. The model accurately predicted broad observed patterns in these statistics (Figure S22). We also computed expected IICR curves (Figure S21) and compared them to previous PSMC estimates (e.g., Prüfer et al., 2014). Although this model features only piecewise-constant effective population sizes which tend to span long epochs, it was able to predict some features of empirical PSMC curves. In predicted and empirical curves, Neanderthal samples have marginally higher IICR than AMH samples before 800 kya, and Denisovan IICR is higher still. More recently, Neanderthal and Denisovan IICR decline then drop precipitously shortly before sampling. In our model, the patterns of ancient IICR inflation in Denisovans and Neanderthals are driven by introgression and mirror the drop in ICR seen under models of structure (cf. Figure 2D).

For almost all parameters, there is substantial overlap in 95% CI obtained from the Bhérer and Zhou datasets (Table S7). The most notable differences are a smaller size and briefer existence for ND when the model is fitted to the Zhou dataset. To evaluate the robustness of estimates to mutation rate specification, we also fit the focal model to the Bhérer dataset using relatively low and high fixed mutation rates (1.1 *×* 10^−8^ and 1.5 *×* 10^−8^ per bp per generation) (Table S8). In general, using a lower mutation rate inflated effective size and time parameters, while a higher rate diminished them, as expected. The behavior of introgression proportions is difficult to interpret, but demonstrates that estimates of these parameters are sensitive to mutation rate specification. To evaluate the robustness of estimated parameters to the presence of Superarchaic-to-Denisovan introgression, we also fit a version of the focal model which lacked this feature (Table S9). Most estimates remain near the center of the focal model CI, with the exception of the introgression proportion *γ*_AMH→AN_, which increased from 5.4% (2.82 – 7.02%) to 8.6% (6.39–11.3%).

### Assumptions and caveats

Like other genetic studies of demographic history, our work is susceptible to biases caused by model misspecification and unrealistic biological assumptions. We tried to avoid the first of these by performing a preliminary exploration of plausible models, then applying a formal statistical test to discriminate between relatively well-fitting alternatives. Nonetheless, we made many approximations to simplify our models. We treated populations as discrete entities, with random mating, piecewise-constant sizes, and instantaneous divergence. Some of these assumptions allow us to model the evolution of HR statistics, while others are useful for formally testing tractable demographic models. Of course, the true evolutionary history includes unmodeled populations, continuously fluctuating population sizes, population structure induced by the spatial distribution of individuals, and variable migration rates. Although neglecting these complexities leads to a degree of model misspecification, it allows us to fit interpretable models with identifiable parameters. Inferred parameters are conditional on the proposed models, which allow us to test statistical support for features of evolutionary importance but could be sensitive to the relaxation of assumptions or the incorporation of additional complexity (Chikhi et al., 2026).

We also made a number of simplifying biological assumptions. We assumed that the genome-wide average germline mutation rate was constant across all lineages throughout the modeled period, which may be roughly acceptable, as the mean germline mutation rate appears unlikely to have changed dramatically throughout the timespan relevant to human diversity (Scally, 2016). We attempted to account for uncertainty in the value of the mutation rate by fitting the focal model with several different fixed rates – alternatively, we could have fit the mutation rate as a free parameter, which may tend to widen CIs for all parameters. We also assumed a constant generation time for all lineages throughout the studied period. This assumption is not merely a scaling of time parameters, because we specified sampling times and fixed event times in physical units (see Supplementary Section 3.7 for a discussion of mutation rates and generation times). We further assumed that the mutation and recombination maps used to estimate *H*_2_ have remained static throughout time, though these will have evolved somewhat over the time period considered here (Harris, 2015; Spence and Song, 2019).

## Conclusion

In this work, we developed a new method for inferring demographic models from single unphased diploid genomes based on *H*_2_ statistics and demonstrated its applicability to high-coverage aDNA. Our approach is flexible and efficient, allowing for the inference of demographic models with arbitrary complexity within the limits of parameter identifiability and computational tractability – the time to compute extended HR expectations scales exponentially with the number of modeled populations. Some techniques for estimating population histories from aDNA infer objects like admixture graphs (*F* -statistics) or IICR curves (PSMC) rather than parameters associated with prescribed demographic models. Others, including *momi* (Kamm et al., 2020) and *Legofit* (Rogers, 2019), explicitly infer demographic models using one-locus diversity patterns. Theory shows that two-locus statistics (and related structures like the PSMC transition matrix – see Cousins et al., 2025) summarize information about the joint distribution of coalescence times which gives them power to identify ancestral reticulations that may be invisible to summaries of one-locus pairwise diversity. Using these advances, we inferred a demographic model that broadly explained observed *H*_2_ patterns and integrated major supported features in hominin evolution, including recurrent interbreeding between Neanderthals and AMH, introgression from a distantly-related, unsampled lineage to Denisovans, and population structure in western Eurasian AMH.

We implemented the BLRT in our framework to formally evaluate support for hypothesized model features. Using simulation, we showed that when models are well-specified, this test has power to detect ancient introgression events when minor ancestry proportions are not too small. In this study, we took some steps towards embedding hypothesized demographic features in a hierarchy of nested models – an approach which could be taken much further in future work, especially if less computationally expensive statistical tests can be devised (e.g., methods summarized by Coffman et al., 2016). After verifying statistical support for the features of interest, we learned parameters jointly in a focal model. By learning parameters jointly, we aimed to avoid the type of confounding that we highlighted in the subset models and demonstrated using simulation. The inference of parameterized demographic models comes with challenges – we only see the topologies that we explicitly select, and we currently lack a systematic way to effectively explore model space. Although simulation shows that we are able to accurately recover parameters when models are well-specified, we caution that unmodeled features may confound demographic inferences and interpretations, not only in this work but broadly in the field.

The *H*_2_ family of statistics may be useful in many further settings. It could be deployed alongside other extended HR statistics, allowing inference that combines high-quality ancient genomes with large samples from contemporary populations. Informed by current work on the structure of the ancestral human metapopulation, this could expand our understanding of human history by permitting lineages represented only by aDNA to be incorporated directly into these models. Although we have confined ourselves to aDNA samples which were sequenced to high depth, techniques to estimate *H*_2_ accurately from low-coverage whole-genome sequences are the subject of ongoing work and will broaden the scope within which the methods developed here can be applied. Because many questions asked with aDNA inference concern the history of the upper Late Pleistocene and Holocene (e.g., the peopling of the Americas – see Willerslev and Meltzer, 2021, for a review), this work should involve a careful characterization of the sensitivity of 𝔼[*H*_2_] to demographic change across short timespans. Here, we have focused on events that occurred in the distant past – the efficacy of the methods developed here for studying more recent events is an open question.

## Methods

### Accounting for local recombination and mutation rate variation

To estimate *H*_2_ from sequence data, we aggregated information across a vast number of pairs of sites. Here we discuss how we account for local variation in genomic maps when averaging across the genome; we leave in-depth discussions of two-site *H*_2_ estimators to Supplementary Sections 2.2 – 2.3. To capture local recombination rate variation, we binned site pairs by the genetic distance between them using an estimated recombination map. We expressed recombination bin edges using recombination fractions *r* to facilitate a comparison with model expectations. Genetic map distances can be transformed to recombination fractions using Haldane’s mapping function (Haldane, 1919), but in practice we transformed bin edges given in units of *r* into Morgans using the inverse of the mapping function in order to reduce the computational cost of binning site pairs. Recombination maps assign map positions to the sites at the edges of irregular genomic windows; we used linear interpolation to estimate the map coordinates of sites within windows.

We estimated *H*_2_ by computing a numerator that sums over observed two-locus diversity and a denominator that sums over the total number of observed nucleotide pairs. The denominator was determined by a genetic mask, which describes which sites are considered to be accessible, and a recombination map. We forbade the counting of site pairs that span centromeres due to uncertainty about the intervening recombination distances. Let *x, y* index sites that pass the mask, let *S*_*k*_ be the set of indices (*x, y*) for site pairs assigned to recombination bin *k*, and let *n*_*k*_ be the cardinality of *S*_*k*_ (see Supplementary Section 2.4 more discussion of site pair counting). Then *H*_2_ on some contiguous region of the genome can be estimated by dividing the sum of site pair *Ĥ*_2_(*x, y*) by the number of accessible site pairs *n*_*k*_:

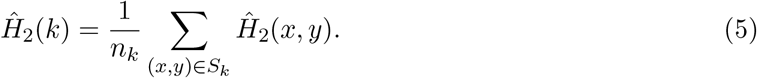

In practice, it is more efficient to restrict the sum over *Ĥ*_2_(*x, y*) to include only polymorphic sites, as invariant sites cannot contribute to *Ĥ*_2_.

Many variables, including nucleotide context, methylation, and replication timing, influence the germline mutation rate at a particular site (Ségurel et al., 2014). Hence, there is substantial local mutation rate variation in the human genome, which has been modeled at the site resolution (Carlson et al., 2018; Seplyarskiy et al., 2023; Karczewski et al., 2020). Recall that 𝔼[*H*_2_] is proportional to *µ*^2^; if we allow mutation rates to differ at left and right loci, we can write 𝔼[*H*_2_] ∝ *µ*_*L*_*µ*_*R*_. Using simulation (see Supplementary Section 4.3), we found that heterogeneous mutation maps can distort the *H*_2_ decay curve (Figure S20). This distortion is unrelated to the genealogy of samples, and instead reflects an unequal average *µ*_*L*_*µ*_*R*_ across bins that emerges due to the autocorrelation of local mutation rates. We found similar features in empirical curves and propose that they may be caused in part by local mutation rate variation (Figure S19).

To mitigate confounding caused by this phenomenon, we weighted site pairs by relative mutation rates. Let *u*(*x*) be the estimated mutation rate at site *x* and *ū* be the genome-wide average; the adjusted estimator for bin *k* is

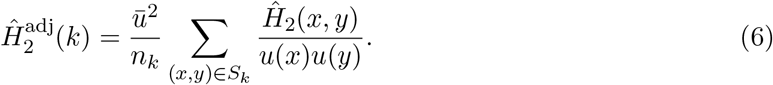

This adjustment, which we apply to all empirical statistics in this work, is intended to scale bin *Ĥ*_2_(*k*) so that the entire curve corresponds to a single average estimated mutation rate, *ū*. It diminishes contributions to observed *H*_2_ from site pairs with relatively high estimated mutation rates and inflates contributions from pairs with low rates. The adjustment performed well under simulation, restoring the observed decay curve to correspondence with its expectation (Figure S20). To use this adjustment, we required fine-scale estimates of the human germline mutation rate – information about the empirical map we used is provided in Obtaining data.

### Obtaining data

We downloaded high-coverage sequence data for three Neanderthals, a Denisovan, and three ancient AMH from the Max Planck Institute for Evolutionary Anthropology and obtained the genome sequence of a contemporary Yoruba individual from the Simons Genome Diversity Project (SGDP) (Mallick et al., 2016) (Table 2, and see Supplementary Section 1.2). To perform inference with aDNA, we needed sample ages specifying the approximate time at which each ancient individual lived. We relied on physical (radiometric) estimates of sample age rather than molecular ones to avoid a potential source of circularity (see Supplementary Section 3.6 and Table S1). We restricted our analyses to sites that pass the 1000 Genomes combined accessibility mask and a set of filters specifying the callability of aDNA samples (see Supplementary Section 1.3). To mitigate the influence of linked and direct selection on our inferences, we excluded sites within 10^−4^ Morgans of CCDS exons (Pruitt et al., 2009). To compute focal site-exon map distances, we used the recombination map inferred by Bhérer et al. (2017).

To evaluate the robustness of our estimates and inferences to recombination map specification, we fit models to *H*_2_ datasets estimated with two recombination maps inferred from non-overlapping datasets with different methodologies (see Supplementary Section 1.4). Bhérer et al. (2017) used the distribution of crossovers in a large pedigree dataset drawn primarily from European populations, while Zhou et al. (2020) utilized inferred IBD tracts in a sample set composed of unrelated African-Americans. A direct comparison of statistics computed with these maps is given in Figure S15. To avoid a possible source of circularity, we did not use recombination maps inferred from LD patterns in our analyses. We plot *H*_2_ curves estimated with two LD-based recombination maps in Figure S17. To compute mutation rate weightings, we downloaded the site-resolution mutation map inferred by Seplyarskiy et al. (2023) (see Supplementary Section 1.5). Their model *Roulette* incorporated extended nucleotide context and other genomic covariates of the mutation rate to explain variation in putative young mutations (rare variants from a large sequence dataset).

### Inference and model choice

We binned site pairs by genetic distance into *n* = 16 log-spaced bins extending from *r* = 10^−6^ to *r* = 10^−2^. In the human genome, *r* = 10^−6^ corresponds to an average physical distance of 100 base pairs, with considerable local heterogeneity. The lower bound was motivated in part by the strong violation of our mutation assumptions entailed by multi-nucleotide mutation events, which tend to occur at distances of tens or hundreds of base pairs (Harris and Nielsen, 2014). We treated the within- and between-population statistics **x**_*k*_ in bin *k* as multivariate normally distributed and estimated the covariance matrix **Σ**_*k*_ with the block bootstrap (see Supplementary Section 1.6). Let *M* denote a model topology and associated parameterization and let *θ* be a vector of parameters. We obtained model expectations **y**_*k*_(*M, θ*) using the extended HR engine implemented in `moments.LD` (Ragsdale and Gravel, 2019) (see Supplementary Section 3.2). The likelihood function for bin *k* is

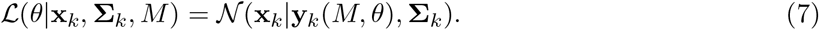

The objective likelihood function is composite, being the product of the *n* binwise likelihoods. In practice, we drop the coefficients of normal distributions because they are constants.

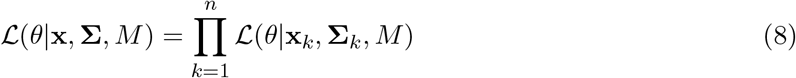

We used algorithms implemented in `scipy.optimize` to explore parameter space and minimize the negative logarithm of the objective likelihood function (Virtanen et al., 2020; Powell, 1964; Nelder and Mead, 1965) (see Supplementary Section 3.3). We can quantify the degree of uncertainty in MLE parameters either by refitting models to bootstrap samples and considering the distribution of bootstrap MLE, or by computing covariance matrices for *θ* in the form of the Godambe Information Matrix (GIM) (as described by Coffman et al., 2016). These methods generally give similar results. Throughout this work, we used empirical 95% CIs obtained with the bootstrap, because we considered them more robust and more readily interpretable (see Supplementary Section 3.4). To discriminate between null and nested alternate models, we used the bootstrap likelihood-ratio test (BLRT) (see Supplementary Section 3.5).

## Code and software availability

We downloaded the empirical data used in this project from publicly available sources specified in Supplementary Section 1. A Python package containing estimation and inference functions (`dpluspy`) is available at https://github.com/nwcol/dpluspy. A GitHub repository with scripts to reproduce datasets, create figures, and run simulated and empirical analyses may be found at https://github.com/nwcol/ancient_introgression_paper. We created model plots with `demesdraw`, available at https://github.com/grahamgower/demesdraw.

## Supporting information

Supplementary objects may be found the **supplementary material** (PDF).

## Acknowledgments

We thank Gustavo Barroso for thoughtful comments on the manuscript. We also thank the UW–Madison Center for High Throughout Computing (CHTC) for their kind assistance and the use of their computational resources (Center for High Throughput Computing, 2006). This work was supported by NIH Award R35-GM154962 to APR.

## Appendix

### A The relationship between 𝔼[*H*_2_] and the Hill-Robertson System

The canonical HR system (Hill and Robertson, 1968) consists of the four-haplotype moments 𝔼[*D*^2^], 𝔼[*Dz*], and 𝔼[*π*_2_]. These moments participate in a closed set of equations which is sufficient to model the evolution of 𝔼[*D*^2^], which is the variance of the basic covariance LD measure *D*. The other two statistics are defined as

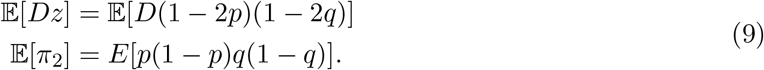

We call these “four-haplotype” statistics because we require four phased genome copies to estimate them from data. 𝔼[*π*_2_] is the probability that, taking an ordered sample of four haplotypes, we observe difference-by-state between the first two haplotypes at the left locus and between the second two haplotypes at the right locus. It is analogous to 𝔼[*H*_2_], except that 𝔼[*π*_2_] considers diversity between a distinct pair of haplotypes at each locus. 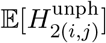 is in some sense intermediate between 𝔼[*π*_2_] and 𝔼[*H*_2_], because there is uncertainty about whether the allele copies sampled from a diploid originated on the same haplotype or on different ones. It can be shown that

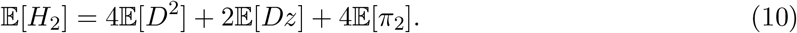

That 𝔼[*H*_2_] is a linear combination of four-haplotype HR statistics is intuitively related to the fact that recombination during the ancestral process can separate allele copies from two sampled haplotypes onto as many as four ancestral gametes. As *r* → ∞, 𝔼[*H*_2_] converges to 4𝔼[*π*_2_] and the other moments approach zero; at loci with uncorrelated genealogies, the probability of observing double polymorphism should be equal regardless of whether we observe loci in the same pair of haplotypes (*H*_2_) or in two different pairs of haplotypes (*π*_2_). The factor of four comes from the fact that 𝔼[*π*_2_] is a *sorted* statistic, and measures the probability of observing a specific sequence of haplotypes (e.g., *a*·, *A*·, ·*b*, ·*B*, where · marks arbitrary alleles) in an ordered sample.

Ragsdale and Gravel (2019) extended the HR system to higher moments (*D*^4^, *D*^6^, etc.) and implemented population divergence and gene flow into the system. This required the addition of multi-population statistics, such as 𝔼[*D*_*i*_*D*_*j*_], where *i, j, k, l* index populations. Let *z*_(*i,j*)_ = (1−2*p*_*i*_)(1−2*p*_*j*_) and *π*_2(*i,j,k,l*)_ = *p*_*i*_(1 − *p*_*j*_)*q*_*k*_(1 − *q*_*l*_). Cross-population 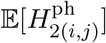 is a linear combination of the multi-population parallels to the canonical HR statistics:

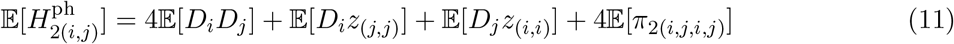

The unphased between-population statistic equals 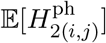 following a generation of free recombination (*r* = 1*/*2), but without drift, mutation or migration. Recombination decays terms of 𝔼[*D*] by the recursion 𝔼[*D*]_*t*+1_ = (1 − *r*)𝔼[*D*]_*t*_, so we can find 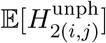 by halving these terms in Equation 11.

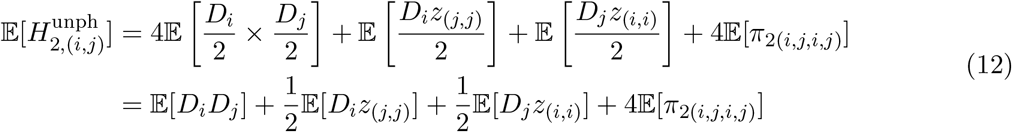

Assuming random mating, interposing a round of free recombination is equivalent to randomly exchanging allele copies between haplotypes in a diploid sample. Averaging over the possible outcomes of free recombination models the lack of phasing information in genotype data.

### B How 𝔼[*H*_2_] captures genealogical covariance

#### The within-diploid statistic

𝔼[*H*_2_] decays as the recombination fraction between loci increases, reflecting the decay of genealogical covariance – pairs of genealogies at loci separated by large recombination distances are more likely to be dissociated by ancestral recombination than those at tightly-linked loci. We assume an infinite-sites mutation model, so that we can ignore multiple- or back-mutations. We also assume that mutation occurs with equal rate *µ* at each locus, although this assumption is easily relaxed. Suppose we sample a genome from a diploid population. Conditioning on coalescence times at left and right loci (*T*_*L*_ and *T*_*R*_), the probability of observing a polymorphism at locus *L* is 𝔼[*H*_*L*_ | *T*_*L*_] = 2*µT*_*L*_. As 𝔼[*H*_2_ | *T*_*L*_, *T*_*R*_] = 𝔼[*H*_*L*_*H*_*R*_ | *T*_*L*_, *T*_*R*_], and because (conditional on the coalescence times) the neutral mutation process is independent of the sample genealogy and independent at each locus (Hudson et al., 1990),

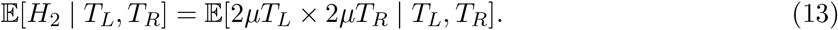

Integrating over the joint distribution of *T*_*L*_ and *T*_*R*_ to obtain an unconditional expectation gives 𝔼[*H*_2_] = 4*µ*^2^𝔼[*T*_*L*_*T*_*R*_]. After substituting with an identity of the covariance, we obtain a relation between 𝔼[*H*_2_] and Cov(*T*_*L*_, *T*_*R*_).

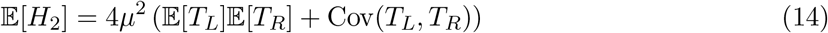

This expression for 𝔼[*H*_2_] is agnostic to the specific history of the population, but depends on our assumption that mutations follow the infinite-sites model.

Now consider a sample from an equilibrium population with diploid size *N*_*e*_. We measure time in coalescent units *τ*_*L*_ = *T*_*L*_*/*2*N*_*e*_, so that 𝔼[*T*_*L*_] = 𝔼[*T*_*R*_] = 2*N*_*e*_, and

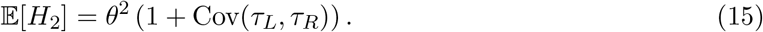

Because Var(*τ*_*L*_) = 1 at demographic equilibrium, Cov(*τ*_*L*_, *τ*_*R*_) = corr(*τ*_*L*_, *τ*_*R*_). Kaplan and Hudson (1985) found an analytical expression for equilibrium Cov(*τ*_*L*_, *τ*_*R*_) in terms of *ρ* = 4*N*_*e*_*r*, where *r* is the recombination fraction between the two loci (Figure 1A).

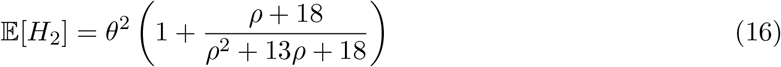

Because 𝔼[*H*_*L*_] = 𝔼[*H*_*R*_] = *θ*, 𝔼[*H*_2_] approaches 𝔼[*H*]^2^ = *θ*^2^ as *ρ* → ∞, and at *ρ* = 0 we have 𝔼[*H*_2_] = 2𝔼[*H*]^2^. Weakly linked loci (*ρ* ≫ 1) have virtually uncorrelated genealogies, because the allele copies on each haplotype tend to be quickly dissociated by recombination and to coalesce separately. Conversely, tightly linked pairs of loci (*ρ* ≪ 1) are only rarely dissociated by ancestral recombination in the history of a sample, and tend to coalesce at the same time. They approximate the behavior of non-recombining loci, which always share the same genealogy, with Cov(*τ*_*L*_, *τ*_*R*_) = Var(*τ*_*L*_) = 1.

Genealogical covariance is related to the probability that coalescence occurs simultaneously at the left and right loci, between gametes that both carry ancestral material at each locus; at equilibrium, Cov(*τ*_*L*_, *τ*_*R*_) = P(*τ*_*L*_ = *τ*_*R*_) (Simonsen and Churchill, 1997). Loci linked by *ρ* ∼ 1 have an intermediate probability of double coalescence, because the rates of coalescence (1, when time is scaled by 2*N*_*e*_) and recombination (*ρ/*2) are similar in magnitude and (roughly speaking) ancestral allele copies tend to be maintained together on the same gametes for a moderate fraction of time before the first coalescence.

Because *ρ* is a function of *N*_*e*_, the effective population size influences not only the scale of the 𝔼[*H*_2_] curve (through *θ*^2^ = (4*N*_*e*_*µ*)^2^), but also the position of the center of the decay curve when it is plotted against physical recombination units *r* (Figure S4).

#### The between-diploid statistic

Throughout this work, we dealt with genotype rather than haplotype data, because phasing aDNA is challenging. We therefore used an *unphased* between-diploid or between-population statistic 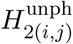. This statistic captures several components of genealogical covariance, which we elucidate here. Suppose that we sample individual diploids from two populations labeled *i, j*. In each diploid, we observe a two-locus genotype that reflects an unknown *latent* haplotype arrangement. Let latent haplotypes from each population/diploid be labeled *i*_0_, *i*_1_ and *j*_0_, *j*_1_. Let *A*_0_*A*_1_*B*_0_*B*_1_ represent the allele copies that make up a two-locus genotype, where subscripts denote identity rather than state. To understand how 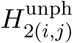 relates to underlying haplotypes, we can imagine assembling allele copies into four synthetic haplotypes *A*_0_*B*_0_, *A*_0_*B*_1_, *A*_1_*B*_0_, *A*_1_*B*_1_. If we assume that coupling and repulsion phases are equally probable when there is polymorphism (see Supplementary Section 2.3), then the identities of allele copies on a synthetic haplotype will match one of the latent haplotypes with probability 1*/*2 (e.g., *A*_0_*B*_0_ and *A*_1_*B*_1_). We assign each synthetic haplotype an equal probability of representing a latent haplotype, then average across 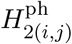 for each between-diploid pair of synthetic haplotypes.

Seen another way, we can imagine taking the latent haplotypes from each diploid and throwing an assignment to the sampled synthetic haplotype onto them randomly at each site. Let 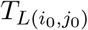 stand for the coalescence time between left allele copies on latent haplotypes *i*_0_, *j*_0_. With probability 1*/*4, synthetic haplotypes drawn from each diploid sample both match latent haplotypes, and the coalescence times between the sampled allele copies are (without loss of generality– haplotypes are exchangeable) 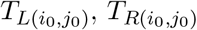. Recall that 𝔼[*H*_2_] is proportional to 𝔼[*T*_*L*_*T*_*R*_]. Enumerating other ways of throwing assignments onto the latent haplotypes yields

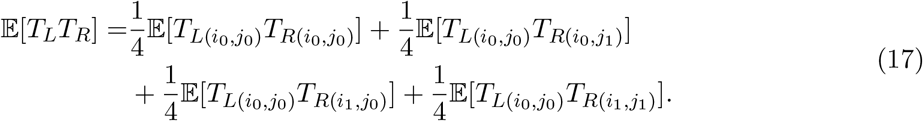

Pairwise coalescence times of allele copies which come from entirely distinct latent haplotypes (e.g. 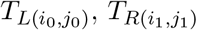) are correlated because there is some probability that one or more pairs of them may coalesce onto the same gamete during the ancestral process. Marginal coalescence times are all equivalent, so letting *T*_*L*_, *T*_*R*_ stand for coalescence times without respect to the particular identities of allele copies,

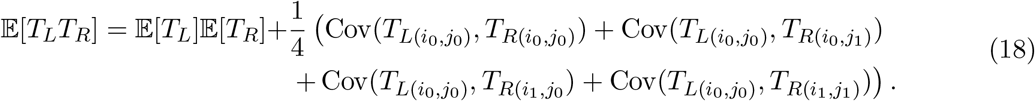

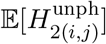 reflects genealogical covariances associated with several sample configurations, unlike 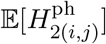, which captures the same covariance term as the within-diploid statistic, 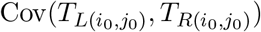). Notably, 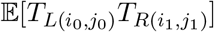 is proportional to the HR statistic 𝔼[*π*_2(*x,y,x,y*)_].

### C Genealogical covariance holds information about population history

#### Normalizing by one-locus diversity

We can better understand the sensitivity of 𝔼[*H*_2_] to demographic events by reasoning about how population history alters the ancestral process and the joint distribution of pairwise coalescence times. To emphasize effects on genealogical covariance, and de-emphasize the changes in one-locus diversity induced by demographic processes, we introduce a statistic 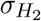 which is normalized by expected one-locus diversity. The expression of 𝔼[*H*_2_] in Equation 14 includes the product of single-locus expectations 4*µ*^2^*𝔼[T*_*L*_]*𝔼[T*_*R*_] = *𝔼[H*_*L*_]*𝔼[H*_*R*_]. We normalize 𝔼[*H*_2_] by this product to obtain

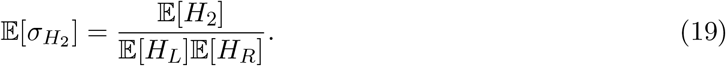

The normalized statistic is related to the genealogical covariance by Equation 14 as

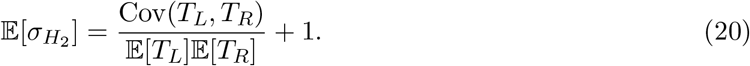

At equilibrium, 𝔼[*T*_*L*_] = Var(*T*_*L*_) = 1, so we can write 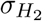] = corr(*T*_*L*_, *T*_*R*_) + 1. When mutation rates at left and right loci are unequal, 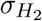 negates confounding by mutation variation, because mutation rates cancel from it. Because 𝔼[*H*_2_] approaches 𝔼[*H*]^2^ as *ρ* goes to infinity, the normalized statistic converges to 1 at long recombination distances. We also define an unphased between-population analog 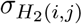 by replacing *H*_2_ with *H*_2(*i,j*) unphased_ and *H*_*L*_, *H*_*R*_ with *H*_*L*(*i,j*)_, *H*_*R*(*i,j*)_. Note that 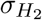 is normalized by a different denominator than other weighted two-locus statistics, such as 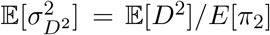 (Ohta and Kimura, 1969), which were also introduced to make measures of LD independent of one-locus diversity, and which have also been applied in demographic inference (Ragsdale et al., 2023). Normalizing by 𝔼[*π*_2_] may be a useful step in future inference approaches.

#### Population size change

Non-equilibrium histories induce transient changes in the joint distribution of coalescence times. For example, population growth or contraction alter the rate of coalescence, so that with passing time, a changing proportion of coalescence events happen in the epoch following departure from equilibrium and the ancestral epoch. Changes in the average shape of gene trees lead to different average amounts of time in which recombination can act to reduce genealogical correlation. For instance, population growth produces a 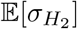 curve which is depressed relative to the equilibrium expectation at tightly linked sites (Figure 1C), because for recombining loci, an increase in average pairwise coalescence time (following expansion) gives recombination more time to act at a given physical distance *r* than before. At short distances, 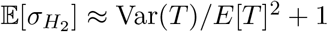, and the behavior parallels the well-known observations about variance in *T*_MRCA_ under changing population sizes (Wakely, 2008): variance in coalescence time falls following a population expansion and rises after a contraction. Thus, population expansion leads to a relatively flat 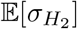 curve, and contraction produces a steep curve which can be dramatically elevated above the equilibrium shape (Figure 1D). Due to the slow rate at which mutation increases 𝔼[*H*_2_] (doubly heterozygous sites must have experienced two independent mutations), observations in populations which have recently undergone bottlenecks followed by expansions are expected to be dominated by the effects of the initial contraction, and expansion takes a long time to dramatically alter the shape of 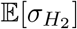.

#### Multi-population dynamics

Multi-population demographic processes (such as population isolation and introgression) also affect genealogical covariance, which is reflected in 𝔼[*H*_2(*i,j*) unphased_] and 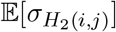 as discussed in Appendix B. Suppose that two populations of equal size *i, j* have diverged at time *τ*_0_ before the present, and that we sample one haplotype from each population. Because the populations are isolated, coalescence cannot occur at either locus until both haplotypes find themselves in the ancestral epoch. For non-recombining loci, this means that the genealogical covariance remains constant with increasing *τ*_0_, while 𝔼[*T*] increases in direct proportion to *τ*_0_. At loci with *r >* 0, recombination during the isolation epoch *t < τ*_0_ decreases genealogical covariance by dissociating ancestral allele copies and increasing the probability that coalescence at the right and left loci will occur at different times in the ancestral epoch. For both these reasons, the 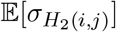 curve becomes increasingly flat as divergence time increases (Figure 1E), reflecting the shrinking magnitude of Cov(*T*_*L*_, *T*_*R*_) relative to the expected coalescence time in the denominator (Equation 20).

We can construct a model of introgression by letting one (donor) population contribute an ancestry proportion *γ* to the other (recipient) population at time *τ*_1_ *< τ*_0_. As we increase the isolation time *τ*_0_ − *τ*_1_, the 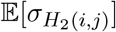 curve for the recipient population broadly increases as positive covariance is induced across *r*, while the relative excess covariance of tightly-linked sites over weakly-linked sites drops (Figure 1F). We can understand this by considering two haplotypes sampled from the recipient population immediately following introgression (in the *F*_1_ generation, when there has not yet been recombination between gametes with different ancestries). Haplotypes are inherited from different parental populations with probability 2*γ*(1 − *γ*). Such samples will have divergence-like histories, because they are unable to coalesce outside the ancestral epoch. This accounts for the drop in the relative excess genealogical covariance at tightly-linked sites, much like in the divergence model. Positive covariance is induced across *r* because isolated haplotypes have correlated ancestries due to population structure. At large *r*, this pattern parallels the elevation of long-distance LD observed in admixed populations (Loh et al., 2013) and is rapidly eroded by recombination in the generations following introgression (Figure 1F).

