## Supplemental Material for "Inferring hominin history with recurrent gene flow from single unphased genomes and a two-locus statistic"

April 11, 2026

**6 Supplementary Tables** **17**

**7 Supplementary Figures** **25**

### 1 Obtaining and preparing data

#### 1.1 Overview

Scripts for reproducing the datasets and analyses used in this study are available in the project repository at `https://github.com/nwcol/ancient_introgression_paper`, where download links for data from external sources are also given. A major project dependency, `dpluspy`, can be found in the repository at `https://github.com/nwcol/dpluspy`. This package contains extensions to the `moments-LD` software tailored to estimate $H_2$ from sequence data, perform the block bootstrap, and fit demographic models. It takes its name from $D^+$, an earlier name for the $H_2$ statistic (the equation for $\mathbb{E}[H_2]$ in haplotype frequencies resembles $D = f_{AB}f_{ab} - f_{Ab}f_{aB}$ with a sign flip).

#### 1.2 Obtaining sequence data

Throughout, we used data in human genome build GRCh37. We downloaded genomic VCF (GVCF) files for ancient samples (Meyer 2012; Prüfer et al. 2014; Prüfer et al. 2017; Mafessoni et al. 2020; Lazaridis et al. 2014), and obtained VCF files for modern sequences from the Simons Genome Diversity Project (SGDP) (Mallick et al., 2016). Although we used data obtained from a single contemporary person from the Yoruba population in Nigeria, labeled Yoruba-1 (`LP6005442-DNA_B02`) in the analyses presented here, we provide an extended dataset with more modern individuals in the project repository. We processed these VCF files using `bcftools` (Danecek et al., 2021). First, we extracted sites with non-homozygous reference genotypes from ancient sample GVCFs using `bcftools view` (in this step we lose information about aDNA coverage, which is required to compute the denominator of the statistic; however, this information can be obtained from sample-specific BED files discussed in the next section). After extracting variant sites from GVCFs, we simplified the resulting VCF files using `bcftools view` and merged them together with sample-specific SGDP VCFs using `bcftools merge`. Because our estimators can accommodate multiallelic sites, we did not filter out multiallelic SNPs. We performed no quality filtering at this stage.

#### 1.3 Filtering sequence data

We downloaded *filterbed* BED files recommended for analyses that use ancient samples. These filters integrate filters on sample-specific site coverage and mappability, and exclude regions marked by structural variation. We took the set of sites passed by every sample-specific filterbed mask as well as the 1000 Genomes combined callability mask as a basis for our genomic mask. From this mask, we excluded sites which fell beyond the edges of the recombination maps that we used in analyses (see Section 1.4). We also masked sites which were not covered by the Roulette mutation map (see Section 1.5). Last, we restricted our analysis to sites which are more than $10^{-4}$ $M$

(0.01 $cM$, or on average $10^4$ bp) from the nearest exon, determined using the Bhérer sex-averaged recombination map (see next section). We chose this threshold because it offered a reasonable tradeoff between maximizing the distance between filter-passing and putatively highly constrained (exonic/promoter) sites, and maximizing the number of sites available for estimating $H_2$. We obtained exon annotations from the CCDS (consensus conding sequence) database (Pruitt et al., 2009), then filtered them to remove untranslated regions (UTRs) and redundant exon IDs and merged any overlaps. As shown in Figure S18, estimated $H_2$ is highly sensitive to the buffer imposed about exonic sites. Larger buffers increase the magnitude of observed $H_2$ across the entire curve relative to $H_2$ computed from filter-passing interexonic sites, suggesting that average selective constraint has a strong effect on the statistic. After these filtering steps, we were left with approximately 663 Mb (million base pairs) of callable, interexonic nucleotides, from which we computed $H_2$ and single-locus statistics as described below.

#### 1.4 Recombination maps

Because local recombination rates vary considerably throughout the human genome, we used estimated sex-averaged recombination maps to compute the distances between sites in order to bin $H_2$ statistics. We used two recombination maps independently inferred by Bhérer et al. (2017) (*Bhérer*) and Zhou et al. (2020) (*Zhou*) in the analyses presented in this work. The Bhérer map was estimated from observed crossovers in several large published pedigree studies drawn predominantly from European populations, while the Zhou map set was estimated using inferred identical-by-descent segments in a sample of unrelated African Americans from the Jackson Heart Study (JHS). Mean $H_2$ computed with each map falls within the confidence interval corresponding to the other map for nearly every bin of every statistic (Figure S15). To avoid a possible source of circularity, we did not use recombination maps estimated from LD patterns in our main analyses. As a supplementary analysis to investigate the robustness of $H_2$ to map specification, we computed the statistic using two such maps (Figure S17). One of these, estimated from 1000 Genomes phase 1 computationally-phased Omni microarray data using the `LDhat` model (Auton et al., 2014), we call *omni*. The other, estimated from 1000 Genomes phase 3 haplotype data using the `pyrho` model by Spence and Song (2019); Kamm et al. (2016), we denote as *pyrho*. With both omni and pyrho datasets, we used the maps estimated using variation from the Yoruba population. $H_2$ estimated with the omni map correspond closely to those from focal non-LD-based maps, while statistics estimated with the pyrho map display a conseriably steeper decay curve which diverges from other maps as $r$ approaches zero.

#### 1.5 Mutation maps

We rescaled site-pair contributions to $H_2$ by the estimated local mutation rate to mitigate a spurious distortion of the observed curve (Section 2.4). This technique requires site-resolution estimates of the mutation rate in humans with broad coverage. We used the *Roulette* mutation map estimated by Seplyarskiy et al. (2023) from rare ($< 0.1\%$ frequency), putatively young SNVs in the gnomAD dataset. The `Roulette` mutation model was fit for mutation types grouped by genomic compartment (gene bodies, promoters, intergenic sites), pentamer context, methylation rate at CpG sites, and gene expression in exons (the last two were measured in testis tissue and discretized). Covariates of the mutation rate modeled by Seplyarskiy et al. (2023) include extended nucleotide context at four nucleotides up- and downstream of pentamers and replication fork direction. In addition to these variables, these workers included the local mutability (as observed in the frequency of rare SNVs) of trimers encapsulated within pentamers, which they argue captures effects associated with local genomic properties, including histone modification and replication timing. The maps are represented in VCF format, with three mutation rates away from the reference sequence at each covered site. For expediency, we transformed the estimated maps into vectors holding the aggregate mutation rate at each site. Note that the genome-wide average of the map used in this adjustment does not need to have any connection to the mutation rate parameter used in inference. The adjustment uses information about the ratio of site rates to the average rate, which cancels their magnitude.

##### 1.6 Bootstrapping

We performed a block bootstrap across genomic regions to estimate the joint distribution of observed $H_2$ statistics for likelihood calculations. We also used the bootstrap to obtain empirical confidence intervals by refitting models to bootstrap replicates (see Section 3.4). We partitioned chromosome arms into non-overlapping windows containing approximately equal numbers of filter-passing sites (1.3 Mb), which yielded 510 windows throughout the genome. Within each window, we computed the numerator and denominator of the $H_2$ statistic, binned by recombination fractions. Where pairs of sites straddled window boundaries, we included them in the window of the left (lower-indexed) site. We forbade the counting of site-pairs that straddled centromeres, due to uncertainty associated with the recombination fractions of such pairs. This rule was unneeded on telocentric chromosomes (e.g., chromosome 22), because the short arms of these chromosomes are highly repetitive and are entirely masked out. To form each bootstrap replicate, we sampled 510 windows from these data with replacement and computed total $H_2$ and $H$ statistics. We assembled 510 such replicates for each dataset and used them to estimate variance-covariance matrices for the $H_2$ statistics in each bin.

#### 2 Estimating statistics from data

##### 2.1 Overview

We categorize $H_2$ statistics and their estimators by whether they consider haplotypes that are both sampled from one diploid (within-diploid) or each from a separate diploid (between-diploid). In this work, the sample size per population is always one diploid, so within-diploid and between-diploid statistics are respectively within-population and between-population measures. We also categorize between-diploid statistics by whether they take haplotype (phased) or genotype (unphased) sequence data. These statistics differ in expectation, and we use the unphased one in our analyses because we work with unphased aDNA sequences. For a single diploid, the phased/unphased distinction is moot, because (assuming perfect confidence in genotype calls) there is no uncertainty about whether a pair of genotypes are both polymorphic or not– and observed $H_2$ is equal for the coupling and repulsion phases. The distinction is important for the between-diploid statistic because we only sample one haplotype from each diploid, and must make assumptions in order to average across the uncertainty in underlying haplotype states.

##### 2.2 Phased estimators

We can write the phased estimator by enumerating the possible ways to sample haplotype pairs in phasing and coupling configurations from a sample, without replacement. The result resembles the haplotype expression of $\mathbb{E}[H_2] = 2\mathbb{E}[f_{AB}f_{ab}] + 2\mathbb{E}[f_{Ab}f_{aB}]$ with a finite sample correction. For a sample of with haplotype counts $(n_{AB}, n_{Ab}, n_{aB}, n_{ab})$ and arbitrary size $n = \sum n_{\cdot\cdot}$,

$$\hat{H}_2^{\text{ph}} = \frac{2n_{AB}n_{ab} + 2n_{Ab}n_{aB}}{n(n-1)}. \tag{S1}$$

In the pertinent case with a single diploid, this simplifies to $\hat{H}_2^{\text{ph}} = n_{AB}n_{ab} + n_{Ab}n_{aB}$. Whether $i$ and $j$ index diploids, or populations with $\geq 1$ sampled haplotypes each, the between-diploid or between-population estimator is

$$\hat{H}_{2(i,j)}^{\text{ph}} = \frac{n_{AB,i}n_{ab,j} + n_{AB,j}n_{ab,i} + n_{Ab,i}n_{aB,j} + n_{Ab,j}n_{aB,i}}{n_i n_j}. \tag{S2}$$

Although we have ignored the possibility that the sample has more than two alleles segregating at a site (though this is impossible in the one-diploid case), it is straightforward to extend these estimators to cases where one or both loci are multi-allelic.

#### 2.3 Unphased estimators

In this work, all within-population measures are estimated from a single diploid and we are unconcerned with estimators for larger sample sizes. We can write the one-diploid estimator using indicators for heterozygosity $\hat{H}_L$ and $\hat{H}_R$, where $\hat{H}_L$ equals 1 when we observe heterozygosity at the left locus and 0 otherwise.

$$\hat{H}_2^{\text{unph}} = \hat{H}_L \hat{H}_R \tag{S3}$$

Although we lack information about whether double heterozygote genotypes ($AaBb$) represent phasing or repulsion haplotype configurations, each configuration contributes equally to $H_2$. A comparison of Equations S1 and S3 shows that given matching haplotype/genotype observations, they will always make the same $H_2$ estimate.

We can begin to think about the between-diploid estimator by proposing to estimate haplotype frequencies or sampling probabilities $\hat{f}_{\cdot\cdot(\cdot)}$ from unphased genotype data.

$$\hat{H}_{2(i,j)}^{\text{unph}} = \hat{f}_{AB,i}\hat{f}_{ab,j} + \hat{f}_{AB,j}\hat{f}_{ab,i} + \hat{f}_{Ab,i}\hat{f}_{aB,j} + \hat{f}_{Ab,j}\hat{f}_{aB,i} \tag{S4}$$

For all two-locus genotypes except $AaBb$, the underlying haplotypes are unambiguous (e.g., $AABb \sim AB$, $Ab$). To deal with double heterozygote genotypes, we make the assumption that phasing and repulsion configurations $AB$, $ab$ and $Ab$, $aB$ are equally probable, or that $D = 0$. This allows us to compute haplotype probabilities directly from observed allele frequencies. Let $\hat{p}_i$, $\hat{q}_i$ be the observed frequencies of alleles $A$, $B$ in population $i$. Then, by assumption, $\hat{f}_{AB,i} = \hat{p}_i\hat{q}_i$, $\hat{f}_{Ab,i} = \hat{p}_i(1-\hat{q}_i)$, and so forth. Substituting haplotype sampling probabilities into Equation S4 and simplifying, we get

$$\hat{H}_{2(i,j)}^{\text{unph}} = \hat{H}_{L(i,j)}\hat{H}_{R(i,j)}. \tag{S5}$$

So $\hat{H}_{2(i,j)}^{\text{unph}}$ can be estimated from pairwise difference probabilities $\hat{H}_{L(i,j)} = \hat{p}_i(1-\hat{p}_j) + \hat{p}_j(1-\hat{p}_i)$, paralleling Equation S3. This estimator is specifically *between-diploid*. With more than one diploid per population, it ignores the individuality of diploids by pooling their allele frequencies and wastes information about the haplotypes which are possible in each diploid (for example, haplotype $Ab$ could not be found in a diploid with genotype $AABB$). In fact, as sample sizes approach infinity and the probability of sampling left/right allele copies that originated on the same haplotype goes to zero, this estimator will become proportional to the HR statistic $\pi_{2(i,j,i.j)}$.

#### 2.4 Approach for estimating genome-wide $H_2$

Our approach for estimating genome-wide $H_2$ from diploids relies on separately tallying the total number of accessible site-pairs (the denominator of the statistic) and the sum of $H_2$ across variant sites (its numerator). Let $k$ index a recombination distance bin defined by the half-open interval $[d_k, d_{k+1})$. In practice, we define recombination bin edges in units of $r$, the recombination fraction, but when binning sites we translate the bin edges into Morgans ($M$) using the inverse Haldane mapping function so that we can treat distances additively;

$$d = \frac{1}{2}\log(1-2r). \tag{S6}$$

Also let $\delta(x, y)$ measure the absolute value of the map distance (in $M$) between loci indexed by $x$, $y$. Define a binning function $b_k(x, y) = \mathbf{1}_{d_k \leq \delta(x,y) < d_{k+1}}$, which returns 1 when the pair $x$, $y$ belongs to recombination bin $k$ and 0 otherwise. Let $\hat{H}_2(x, y)$ measure $H_2$ at sites $x$, $y$; for one diploid, $\hat{H}_2 = \hat{H}(x)\hat{H}(y)$, where $\hat{H}(x)$ is an indicator for observed heterozygosity at site $x$. The aggregate estimated $H_2$ for the $k$th recombination bin measured on some contiguous segment of a chromosome is

$$\hat{H}_2(k) = \frac{\sum_x \sum_{y>x} b_k(x, y)\hat{H}_2(x, y)}{\sum_x \sum_{y>x} b_k(x, y)}. \tag{S7}$$

With a mutation map defining site mutation rates $u(x)$ and a genome-wide average rate $\overline{u}$, the adjustment for local mutation rate variation is

$$\hat{H}_2^{\text{adj}}(k) = \frac{\overline{u}^2}{\sum_x \sum_{y>x} b_k(x, y)} \times \sum_x \sum_{y>x} \frac{b_k(x, y)\hat{H}_2(x, y)}{u(x)u(y)}. \tag{S8}$$

In practice we restrict the sum over $H_2(x, y)$ to only polymorphic sites.

When estimating $H_2$ from a sequence of length $L$, each double sum must pass over $\binom{L}{2}$ terms, so tallying site-pairs with a naive loop in Python is prohibitively expensive for human chromosomes. Instead, we used the `searchsorted()` function from NumPy to find the number of site-pairs in each bin. Passing `searchsorted(a, v)` on a sorted array `a` and array `v` returns the indices of `a` where elements of `v` should be inserted to maintain the sorted order. If `rmap` is a vector of genetic map coordinates, then taking the sum of the vector computed with `np.searchsorted(rmap, rmap + bins[k + 1]) - np.searchsorted(rmap, rmap + bins[k])` gives the count of site-pairs in bin $k$. A slightly different approach was required for the smallest bin if its lower bound is 0, to avoid counting self-self pairs; we replaced the first `np.searchsorted` term with a vector that counts from 1 to $L - 1$. To compute the numerator of within-diploid $H_2$, we restricted `rmap` to include only the map coordinates of sites that are heterozygous in the sample.

To estimate the unphased between-diploid statistic for diploids $i$, $j$, we first computed a vector of pairwise difference probabilities $\hat{H}_{x(i,j)}$ for each variant site. Then we applied Equation S8, only substituting $\hat{H}_2 = \hat{H}_{x(i,j)}\hat{H}_{y(i,j)}$. In practice it would likely be feasible to loop naively over these terms to compute the numerator of $H_{2(i,j)}^{\text{unph}}$, but we instead used a vectorized technique similar to the one described above for tallying site-pairs. In accordance with our bootstrap scheme, we applied these calculations to find the binned denominators (which are shared across diploid samples) and the numerators (which are unique to each diploid and between-diploid pair) within each bootstrap window. We saved or cached raw sums to make future bootstrapping more straightforward (running these computations on a laptop computer took $\sim$30 minutes).

#### 2.5 Estimating one-locus diversity statistics

We estimated $\hat{H}$ and $\hat{H}_{i,j}$, the probability of single-locus pairwise difference between populations $i$ and $j$, to allow comparisons between $\hat{H}_2$ and observed measures of one-locus diversity and to allow a comparison of those measures to predictions from best-fit models. We applied the same filters that we used to estimate $\hat{H}_2$. For one diploid, $\hat{H}$ is just the count of heterozygous sites that pass filters divided by the total number of filter-passing sites. To compute $\hat{H}_{i,j}$ at a site, we count the number of different-by-state allele pairs that can be drawn from populations $i$ and $j$. Letting $X_k$, $Y_l$ be the $k$th, $l$th alleles sampled from populations $i$, $j$, with $X_k \in \{0, 1, ...\}$, with haploid sample sizes $n_i$, $n_j$, then $\hat{H}_{i,j} = \mathbb{P}(X \neq Y)$, or

$$\hat{H}_{i,j} = \frac{1}{n_i n_j} \sum_{k=1}^{n_i} \sum_{l=1}^{n_j} \mathbf{1}_{X_k \neq Y_l} \tag{S9}$$

There is no need to re-weight sites by local mutation rates when estimating one-locus statistics, because as averages or first moments they are not distorted by mutation rate variation.

### 3 Fitting demographic models

#### 3.1 General approach

We treated the $H_2$ statistics in each recombination-distance bin as multivariate Gaussian-distributed, and assumed independence between bins. To model population histories, we made a number of assumptions, which we will make explicit here. We initialized all models from a panmictic ancestral deme, which (in all models discussed in this work) remained at demographic equilibrium until the first population split. We fit the size of the ancestral population as a free parameter $N_\mathrm{A}$. All modeled demes are panmictic with piecewise-constant effective sizes. To model gene flow, we always assumed either symmetric, continuous migration or instantaneous, one-way introgression. We also assumed that population separation was instantaneous and that demes only bifurcate.

To construct complex models, we added complexity gradually. Guided by the literature, we initially fit two- and three-population models to find appropriate topologies (split orders) and rough estimates of effective sizes and divergence times. Before conducting formal tests for, e.g., episodes of gene flow, we fit models including these features to check whether we tended to infer nonzero values for the associated parameters, and whether their inclusion improved model fits. We generally avoided fixing parameters, except where external evidence for a particular value was compelling and parameter constraint appeared weak. Due to the behavior of some introgression time parameters (collision with constraints imposed by deme epochs, or implausible runaway behavior), we fixed several of them at previously-inferred values (the AMH-Neanderthal introgressions) or pinned them to split times (Superarchaic-Denisovan introgression occurs immediately following Denisovan/Neanderthal divergence). In other cases, we fixed parameters which were tightly constrained above and below by other events (for instance, if we assume that the Ust'Ishim lineage shared its Neanderthal ancestry with modern people, then the split time of Ust'Ishim from Loschbour must occur before 45 kya, the approximate lifetime of the Ust'Ishim man, but after the introgression from Neanderthals to Eurasian AMH at 48 kya).

#### 3.2 Model specification

We represented models using the `demes` Python package and the corresponding Demes specification (Gower et al., 2022), a standard format for describing population genetic models. All models discussed in this work are supplied in the project repository in this format. To compute expected statistics from Demes models, we used the `moments-Demes` interface of the `moments` software. This subpackage parses Demes models into the internal data structure used to represent demographic models by `moments` and calculates expected two-locus statistics using `moments-LD`. From the resulting arrays of HR statistics, we computed $H_2$ with the `H2()` method.

To specify model parameterizations, we used the simple YAML format supported by `moments-LD`, which allowed us to flexibly map parameters to features in demographic models (effective population sizes, divergence times, migration rates, and so forth). This format is described in the `moments`documentation available at https://momentsld.github.io/moments/extensions/demes.html. We imposed bounds to prevent parameters from diverging to biologically implausible or model-violating (e.g., negative) values during optimization. Some typical bounds include $10^{-8}$–$10^{-2}$ for migration rates, $10^{-4}$–1 for introgression proportions, and 100–$10^5$ or $10^6$ for effective sizes (with smaller lower bounds in Neanderthal and Denisovan populations). Where necessary, we imposed relative constraints on parameter values; for instance, if populations $AB$ and $C$ split from an ancestral population before $AB$ splits into $A$ and $B$, the divergence time parameters must satisfy $T_{\text{AB–C}} > T_{\text{A–B}}$. When parameters constrained in this way collided (converged to one value), we interpreted it as a sign of model misspecification and adjusted the topology to relax the constraint (e.g., in the prior example, we might hypothesize that either $A$ or $B$ actually shares more recent ancestry with $C$).

#### 3.3 Optimization strategies

When fitting complex models, we used a combination of parallelization and serial re-optimization to find MLE parameters. Throughout, we used numerical optimization functions implemented in the `scipy` Python package (Virtanen et al., 2020). We usually fit multiple copies of each model in parallel on a computation cluster. In each replicate, we perturbed the initial parameters by sampling them uniformly from a fixed interval about some initial guess. We parallelized fits to explore parameter space as extensively as possible and increase the probability that we discover global optima on the likelihood surface. In the very first preliminary fits, and when introducing new parameters, we drew initial guesses from the literature, or arbitrarily selected values which appeared reasonable. Subsequently, we used MLE parameter values from previous rounds of optimization as initial conditions. For each replicate, we typically began optimization using the simplex Nelder-Mead algorithm as implemented in the `scipy.optimize.fmin()` function (Nelder and Mead, 1965). We then alternated between the `scipy.optimize.fmin_l_bfgs_b()` (Byrd et al., 1995; Zhu et al., 1997) and `scipy.optimize.fmin_powell()` (Powell, 1964) functions until the change in log-likelihood fell beneath a fixed threshold, indicating consistent convergence to a point in parameter space (this threshold was more sensitive than the convergence criteria given to `scipy` functions). In some cases we applied a small 'jitter' between rounds of optimization by perturbing parameter values by 1-5% to help the algorithms escape local optima or flat regions of the likelihood surface.

#### 3.4 Quantifying uncertainty

We exploited the block bootstrap procedure described in Section 1.6 to quantify uncertainty in MLE parameter values. To produce confidence intervals (CI) for inferred parameters, we refit the model of interest to a large number (here 510) of block bootstrap samples. We then took the

2.5% and 97.5% quantiles of bootstrap sample MLE as empirical 95% CI. The CI we present are therefore generally asymmetric about the MLE and strictly non-negative. This strategy did not preclude the collision of CI with upper or lower bounds, which we interpreted as a sign of poor parameter constraint if the bounds were chosen reasonably. For instance, collision of the lower CI with the lower bound of a pulse proportion parameter could indicate low statistical support for the modeled pulse. Collisions between CI and parameter bounds are rare in the models we present; they are marked by * in Tables S4-S9. Notably, the effective size of the Ust'Ishim population is poorly constrained, likely due to the brief existence (2 ka) of this deme and the negligible effect of $N_e$ on diversity during this time window. To measure the correspondence between expected and observed statistics, we calculated residuals with $(\text{Model}_{i,j} - \text{Data}_{i,j})/\sqrt{V_{i,j}}$, where $V_{i,j}$ is the bootstrap variance of the $i$ statistic in its $j$th bin.

##### 3.5 Model choice

The conventional likelihood-ratio test (LRT) has an unknown asymptotic null distribution under composite likelihoods, so we cannot use it to perform model choice. Adjustments have been proposed to restore the expected asymptotic null distribution under composite likelihood (for an application to demographic inference, see Coffman et al. 2016), but we were unable to obtain consistent results when we applied these techniques to simulated $H_2$ datasets. Instead, we used the bootstrap likelihood-ratio test (BLRT). To perform the BLRT, we used Monte-Carlo simulation to estimate the null distribution of the likelihood-ratio (LR) statistic. Specifically, we ran a large number of genome-scale coalescent simulations (typically 500) under the MLE null model. We then fit null and alternative models to each simulated dataset. By computing the LR statistic for each replicate, we obtained an empirical null distribution. We then used this distribution to compute empirical $p$-values ($p_{\text{obs}}$) by counting the number of simulated LRT scores greater than the empirical test statistic. Let $LL_{i,0}$ and $LL_{i,1}$ be the log-likelihoods of the null and alternative model in the $i$th simulated dataset. Then with $n$ replicates with scores $\lambda_i = 2(LL_{i,1} - LL_{0,1})$,

$$p_{\text{obs}} = \frac{1}{n} \sum_{i=1}^{n} \mathbf{1}_{\lambda_i > \lambda_{\text{obs}}}. \tag{S10}$$

Using the BLRT obviates the need to determine the degrees of freedom of tests, which can become complex when there are multiple boundary parameters. These are parameters, such as migration rates, which take a value on the boundary of parameter space (usually 0) in the null model. We note that the results of the BLRT are sensitive to model parameterization.

We performed coalescent simulations in `msprime` (Kelleher et al., 2016) using the estimated parameters of the human genome. For each replicate, we carried out ancestry simulations under the relevant null model across 22 autosomal chromosomes of appropriate size. We performed these simulations with an empirical recombination map (Bhérer or Zhou) and simulated mutation with a windowed version of the Roulette mutation map (averaged on 10kb intervals; using a site-resolution mutation map at the genome-wide scale was prohibitively computationally expensive). We emitted the resulting genotypes in VCF files, then applied to these the same procedures (masking, bootstrapping) that we used to estimate empirical statistics. We carried out a simulated demonstration of this BLRT approach in our framework using simple models (see Section 4.5).

##### 3.6 Assigning sample ages to ancient sequences

To fit models that include ancient samples, we must provide point sample age estimates or sampling times. We used ages estimated with physical (radiometric, stratigraphic, etc.) methods

| Label | Est. age (ka) | Technique | Reference |
|---|---|---|---|
| Altai Neanderthal | **110**[1] (90–130) | Multiple[2] | Douka et al. (2019) |
| | 129–136 | Branch shortening | Prüfer et al. (2014) |
| Chagyrskaya Neanderthal | **55**[1] (49–59) | Optical dating of sediments | Kolobova et al. (2020) |
| | 80 | Branch shortening | Mafessoni et al. (2020) |
| Vindija Neanderthal | **45** (42–46) | Radiocarbon | Devièse et al. (2017) |
| | 60–65 | Branch shortening | Prüfer et al. (2017) |
| Denisova Denisovan | **60**[1] (52–76) | Multiple[2] | Douka et al. (2019) |
| | 74–82 | Branch shortening | Meyer (2012) |
| Ust'Ishim AMH | **45** (43–47) | Calibrated radiocarbon | Fu et al. (2014) |
| | 49 (31–66) | Branch shortening | Fu et al. (2014) |
| Loschbour AMH | **8** | Radiocarbon | Toussaint et al. (2009) |
| Stuttgart AMH | **7** | Radiocarbon | Stäuble (2005) |

Table S1: References and technologies for ages assigned to aDNA samples. Where multiple ages estimates are given, the one used in this study is bolded. Ranges indicate 95% confidence intervals given by the respective authors; in some cases (marked [1]), our point age estimates were not given by authors but were selected to fall close to the centers of CIs or at the means of posterior age distributions. [2] Douka et al. (2019) estimated ages using a Bayesian model that incorporated radiometric, stratigraphic and genetic information.

rather than genetic ones (branch-shortening, etc.) to avoid a potential source of circularity. We provide point estimates, confidence intervals, and references for each ancient sample in Table S1. For reference, we provide branch-shortening estimates where they are available. Branch-shortening uses a deficit in accumulated derived alleles on an ancient lineage relative to a modern one to predict sample age. This technique therefore relies on the ascertainment of derived variation and point estimates of the *per annum* mutation rate.

##### 3.7 Selecting the mutation rate parameter

Throughout, we used a generation time of 29 years and a mutation rate $\mu = 1.3 \times 10^{-8}$ bp$^{-1}$ generation$^{-1}$ (or $4.48 \times 10^{-10}$ bp$^{-1}$ year$^{-1}$). We chose this generation time based on a survey of extant human populations by Fenner (2005), who recommended a generation time of 28–30 years for autosomal genetic studies. We implicitly project this assumed generation time into the remote past, an assumption which is difficult to justify but also difficult to avoid in this framework. We adopted the specified mutation rate based on several pedigree studies (Rahbari et al., 2016; Jónsson et al., 2017) which made point estimates of $1.28 \times 10^{-8}$ and $1.29 \times 10^{-8}$ bp$^{-1}$ generation$^{-1}$ with average parental ages of 29.8 and 30.1 years respectively. We caution that these rates are not representative of all pedigree studies, and appear to lie towards the upper end of the reported values (Moorjani et al., 2016). To evaluate the robustness of inferred parameters and model fits to the mutation rate parameter (which alters the shape of $\mathbb{E}[H_2]$ curves due to the change in the ratio of $\rho$ to $\theta$) and provide parameter estimates at different point mutation rates, we refit the full model with $\mu = 1.1 \times 10^{-8}$ bp$^{-1}$ generation$^{-1}$ ($3.79 \times 10^{-10}$ bp$^{-1}$ year$^{-1}$) and $1.5 \times 10^{-8}$ bp$^{-1}$ generation$^{-1}$ ($5.17 \times 10^{-10}$ bp$^{-1}$ year$^{-1}$) (Table S8, Figures S14).

### 4 Simulations

#### 4.1 Two-locus simulations in msprime

$\mathbb{E}[H_2]$ is fundamentally a tractable summary of the joint distribution of pairwise coalescence times. This distribution was analyzed for equilibrium populations by Simonsen and Churchill (1997), but it would be challenging to obtain analytically for non-equilibrium models. To develop intuition about the qualitative behavior of $\mathbb{E}[H_2]$ under various population processes, we used Monte-Carlo coalescent simulation in msprime (Kelleher et al., 2016; Baumdicker et al., 2022) to produce estimated distributions. For a given demographic model, we simulated a large number ($10^5$) of discrete sequences of length 2, separated by a per-bp recombination fraction $r$, then recorded site branch lengths $T_L$ and $T_R$ for each replicate. We show distributions for several models in Figures S1, S2.

#### 4.2 General approach for genome-scale simulations

In the analyses presented in Sections 4.3, 4.4, and 4.5, we used a common approach to generate genome-scale simulated datasets. To create each replicate dataset, we utilized coalescent simulation in msprime (Kelleher et al., 2016; Baumdicker et al., 2022) to independently simulate the 22 human autosomes under some demographic model. To mimic the empirical joint distribution of $H_2$ statistics, we used inferred recombination and mutation maps (otherwise, statistics from simulated data tend to have considerably lower bootstrap variances than empirical observations, which might result in tighter confidence intervals and more sensitive statistical tests than we would expect to see with empirical data). Where not otherwise stated, we performed simulations with the Bhérer recombination map and a 1kb-windowed version of the Roulette mutation map, then applied the same filters and mutation rate adjustments that we used to parse empirical data. We scaled the Roulette map to a genome-wide average rate of $1.3 \times 10^{-8}$ so that the number of mutations we observe in data simulated under inferred models would correspond roughly to that in empirical data (the average rate of the unscaled Roulette map is $\sim 1.11 \times 10^{-8}$ across the sites that pass our filters).

#### 4.3 Local mutation rate variation

Local mutation rate (LMR) variation can distort the $H_2$ curve, causing it to display non-monotonicity and unexpected asymptotic behavior, because rate autocorrelation causes an uneven distribution of average pairwise mutation rates at different recombination distances. We ran simulations to demonstrate the efficacy of the LMR-adjusted $H_2$ estimator. Our objective was to show that the adjustment brings observed $H_2$ curves into closer agreement with their expectations than the LMR-naive statistic when data is generated under a model with realistic patterns of LMR. For this demonstration, we used the focal model with Bhérer dataset MLE parameters.

We computed both LMR-adjusted and LMR-naive $H_2$ from simulated data. Qualitatively, the relative magnitudes and shapes of these curves (Figure S20) closely match empirical LMR-adjusted and naive curves (Figure S19). The adjustment of simulated data brings long-distance $H_2$ into near-equality with observed $H^2$, as expected, and restores monotonicity (non-monotonic decay in LMR-naive curves is slight but apparently present). Empirical data is brought into closer alignment with $H^2$ at long range by LMR adjustment, but continues to display non-monotonic decay at very long distances. The source of this pattern is unclear, but it could result from systematic error in the mutation map which reduces the efficacy of the adjustment, error in the relative mutation rates on different chromosomes (which contribute different fractions of site-pairs to different bins), or the inadequacy of a static map for describing evolution.

##### 4.4 Jointly modeling features mitigates bias in inferences

When jointly fitting model features, we observed shifts in parameter MLE towards values previously reported in the literature. Because model misspecification can bias parameter estimates, we expect that more-detailed models which capture more episodes of gene flow should produce less-biased MLE, when the modeled episodes actually took place (this is true of all model features, not only gene flow events). We constructed a simple model with three episodes of instantaneous gene flow, corresponding in time and proportion to Superarchaic-to-Denisovan, AMH-to-Neanderthal, and Neanderthal-to-OOA events (Figure S23A) to test how model misspecification confounds parameter estimates. To each simulated dataset, we fit eight models with every permutation of presence/absence for each pulse.

As expected, fitting the ground-truth model to data provides a substantially higher likelihood than misspecified models (Figure S23B) and generally produces parameter estimates with lower error (Figure S23C-O). Under this parameterization, not modeling the most recent Neanderthal-Loschbour pulse appears to have a greater effect than excluding the two older pulses. This may be because this event is considerably more recent than the other two gene flow events, or because those events leave correlated signals in patterns of genetic variation. When we do not model either the Superarchaic-to-Denisovan or AMH-to-Neanderthal introgressions, the other pulse proportion is inflated (Figure S23D, E). This corresponds to the patterns observed when fitting models to empirical data. Many other parameters (in particular the effective sizes of the Neanderthal and Denisovan demes and their common ancestor ND) are also biased when estimated with the misspecified models investigated here.

##### 4.5 A demonstration of model choice

We performed a simple demonstration of the BLRT. This analysis was limited in scope by the large computational expense involved in running the BLRT (which requires large numbers of simulated replicates to estimate null distributions) and is intended only as a basic validation of the method in this framework. We dealt with one family of models focused on AMH-to-Neanderthal introgression (Main Text Figure 2C, D) and one family focused on Superarchaic-to-Denisovan gene flow (Main Text Figure 2E, F). Each model family was composed of models with different introgression proportions ($\gamma = 0$, 1%, 5%). We simulated 10 independent focal replicates for each model and applied the BLRT as described in Section 3.5 with 100 samples per replicate. This configuration allowed us to test BLRT performance on datasets where either the null or alternate model was the ground truth.

The results suggest that we have reasonable power to identify introgression events, at least under idealized conditions and when introgression proportions are moderately large ($\gtrsim$ 1%, Table S3). There were no false positives (rejections of the null model when the ground truth model lacked introgression). For AMH-to-Neanderthal models, we rejected the null model in 3/10 and 9/10 cases with $\gamma = 0.01$ and 0.05 at an $\alpha = 0.05$ threshold. For the Superarchaic-to-Denisovan models, we rejected the null model in 4/10 and 10/10 cases at $\gamma = 0.01$ and 0.05, respectively. In this analysis, we ignored the possibility of model misspecification, which may bias hypothesis tests and inferred parameters, by assuming that all features of the model except the presence of introgression were known. We also treated the introgression time as a known quantity and fixed this parameter. Further, we did not model misspecification of the recombination and mutation maps, which could distort observed $H_2$ curves and lead to spurious inference.

### 5 Model predictions

#### 5.1 Patterns of one-locus diversity

As a crude check on our focal model, we made a qualitative comparison between several observed and expected measures of one-locus diversity. We examined heterozygosity $H$, between-population heterozygosity $H_{i,j}$, and the $F$ statistic $F_2$, which is a measure of genetic drift branch length between samples. The expectations of the $F$-statistics can be computed directly from expected $H$ statistics, e.g. where $i$ and $j$ index populations,

$$\mathbb{E}[F_2(i,j)] = \mathbb{E}[H_{i,j}] - \frac{\mathbb{E}[H_i] + \mathbb{E}[H_j]}{2} \tag{S11}$$

and they can be estimated from sequence data using observed $\hat{H}$ and $\hat{H}_{i,j}$ (Peter, 2016). Throughout, we used the same sites to compute $\hat{H}$ statistics as we did to find $\hat{H}_2$ in the preceding analyses. To compute expectations, we applied Equation S11 to terms of $\mathbb{E}[H]$ predicted by `moments-LD` (Jouganous et al., 2017; Ragsdale and Gravel, 2019). In general, the block structure of expected and observed $H$ and $F_2$ statistics is in good correspondence (Figures S22). Expected statistics tend to fall in or near empirical 95% CIs, suggesting that the focal model is a reasonable predictor of overall one-locus diversity patterns.

#### 5.2 Instantaneous coalescence rate profiles

We also computed the inverse instantaneous coalescence rate (IICR) expected under the best-fit full model. Let $T$ be the coalescence time for a pair of allele copies, and $f(t)$ the marginal distribution of $T$. Also let $S(t) = \mathbb{P}(T \geq t)$ be the survival function (the probability that coalescence has not happened yet at time $t$). Then the ICR is the hazard function of $T$, or

$$\lambda(t) = \frac{f(t)}{S(t)}. \tag{S12}$$

At equilibrium, $\lambda(t) = 1/(2N_e)$ and its inverse (the IICR) is $\lambda(t)^{-1} = 2N_e$. In populations with changing sizes, the ICCR contains information about the historic size trajectory.

We calculated IICR profiles using `msprime` by using the `coalescence_rate_trajectory()` on a fine grid of times to obtain an ICR curve for each sampled lineage, then inverting each curve by taking $\lambda(t)^{-1}$. Let $i$ index populations, and let $p(i,t)$ be the probability that two sampled allele copies have not coalesced and are in population $i$ at time $t$. The ICR computed by `msprime` is

$$\lambda(t) = \frac{1}{S(t)} \times \sum_i \frac{p(i,t)}{2N_e(i,t)}. \tag{S13}$$

It is possible to calculate cross-ICR (between allele copies sampled in different populations) using the same method, but we avoid this because it is challenging to infer empirical cross-coalescence rates for unphased data (as this procedure requires haplotype information), so there are no empirical observations to compare against.

### 6 Supplementary Tables

| Test | Dataset | $LL_0$ | $LL_1$ | $\Delta LL$ | $p_{\text{obs}}$ |
|---|---|---|---|---|---|
| (1.1) AMH→AN | Bhérer | -567 | -498 | 69 | $< 0.002$* |
| | Zhou | -538 | -488 | 50 | $< 0.002$* |
| (1.2) AMH→WN \| AMH→AN | Bhérer | -498 | -420 | 78 | $< 0.002$* |
| | Zhou | -488 | -426 | 62 | $< 0.002$* |
| (2.1) S→Denisova \| AMH→AN, AMH→WN | Bhérer | -313 | -300 | 13 | 0.018* |
| | Zhou | -313 | -307 | 5 | 0.052 |
| (2.2) S→Denisova | Bhérer | -591 | -390 | 201 | $< 0.002$* |
| | Zhou | -584 | -411 | 172 | $< 0.002$* |
| (3) BE→Stuttgart \| AMH→AN | Bhérer | -152 | -78 | 73 | $< 0.002$* |
| | Zhou | -176 | -99 | 77 | $< 0.002$* |

Table S2: A summary of BLRT featured in the main text (BLRT sample size $n = 500$), showing the hierarchy of dependence between tests (events assumed in the null model of a test are given after $|$). The $p_{\text{obs}}$ marked with * are significant at the $\alpha = 0.05$ level.

| Model | AMH→N replicate | $\Delta LL$ | $p_{\text{obs}}$ | Model | S→D replicate | $\Delta LL$ | $p_{\text{obs}}$ |
|---|---|---|---|---|---|---|---|
| $\gamma_{\text{H}\to\text{N}} = 0$ | 0 | 4 | 0.11 | $\gamma_{\text{S}\to\text{D}} = 0$ | 0 | 11 | 0.11 |
| | 1 | 1 | 0.3 | | 1 | 11 | 0.1 |
| | 2 | 0 | 0.77 | | 2 | 14 | 0.07 |
| | 3 | 0 | 0.57 | | 3 | 1 | 0.32 |
| | 4 | 0 | 0.36 | | 4 | 0 | 0.57 |
| | 5 | 0 | 0.44 | | 5 | 0 | 0.46 |
| | 6 | 3 | 0.14 | | 6 | 0 | 0.52 |
| | 7 | 0 | 0.53 | | 7 | 2 | 0.26 |
| | 8 | 0 | 0.44 | | 8 | 0 | 0.4 |
| | 9 | 0 | 0.32 | | 9 | 3 | 0.21 |
| $\gamma_{\text{H}\to\text{N}} = 0.01$ | 0 | 0 | 0.54 | $\gamma_{\text{S}\to\text{D}} = 0.01$ | 0 | 29 | 0.01* |
| | 1 | 0 | 0.76 | | 1 | 18 | 0.06 |
| | 2 | 2 | 0.21 | | 2 | 7 | 0.16 |
| | 3 | 0 | 0.8 | | 3 | 0 | 0.59 |
| | 4 | 20 | $< 0.01$* | | 4 | 150 | $< 0.01$* |
| | 5 | 16 | 0.02* | | 5 | 15 | 0.01* |
| | 6 | 13 | 0.05 | | 6 | 33 | $< 0.01$* |
| | 7 | 4 | 0.18 | | 7 | 7 | 0.18 |
| | 8 | 10 | 0.06 | | 8 | 5 | 0.05 |
| | 9 | 18 | 0.03* | | 9 | 7 | 0.08 |
| $\gamma_{\text{H}\to\text{N}} = 0.05$ | 0 | 220 | $< 0.01$* | $\gamma_{\text{S}\to\text{D}} = 0.05$ | 0 | 149 | $< 0.01$* |
| | 1 | 168 | $< 0.01$* | | 1 | 168 | $< 0.01$* |
| | 2 | 117 | $< 0.01$* | | 2 | 419 | $< 0.01$* |
| | 3 | 324 | $< 0.01$* | | 3 | 181 | $< 0.01$* |
| | 4 | 193 | $< 0.01$* | | 4 | 288 | $< 0.01$* |
| | 5 | 16 | 0.08 | | 5 | 162 | $< 0.01$* |
| | 6 | 360 | $< 0.01$* | | 6 | 239 | $< 0.01$* |
| | 7 | 248 | $< 0.01$* | | 7 | 124 | $< 0.01$* |
| | 8 | 134 | $< 0.01$* | | 8 | 204 | $< 0.01$* |
| | 9 | 131 | $< 0.01$* | | 9 | 252 | $< 0.01$* |

Table S3: A summary of simulated BLRT results (with BLRT sample size $n = 100$). $\Delta LL$ are rounded to the nearest integer, and $p_{\text{obs}}$ marked with * are significant at the $\alpha = 0.05$ level.

| | Null model $LL = -568$ | | Two-pulse model $LL = -420$ | |
|---|---|---|---|---|
| Parameter | MLE | 95% CI | MLE | 95% CI |
| $T_{\text{ND–AMH}}$ (kya) | 819 | 760 – 865 | 779 | 731 – 831 |
| $T_{\text{AN–Den}}$ (kya) | 662 | 585 – 781 | 726 | 654 – 788 |
| $T_{\text{WN–Alt}}$ (kya) | 123 | 118 – 133 | 119 | 116 – 125 |
| $T_{\text{Cha–Vin}}$ (kya) | 62.9 | 58.3 – 76.1 | 60.5 | 58 – 66.7 |
| $T_{\text{Yor–Los}}$ (kya) | 48 | 48 – 48 | 48.1 | 48 – 66.8 |
| $T_{\text{Vin}\to\text{Los}}$ | 48 | — | 48 | — |
| $N_{\text{A}}$ | 16600 | 15900 – 17500 | 17100 | 16300 – 17900 |
| $N_{\text{ND}}$ | 12200 | 3630 – 18400 | 2180 | 807 – 4850 |
| $N_{\text{Den}}$ | 2830 | 2570 – 3100 | 3620 | 3150 – 4000 |
| $N_{\text{N}}$ | 3220 | 2950 – 3430 | 2610 | 2300 – 2830 |
| $N_{\text{Alt}}$ | 303 | 127 – 736 | 189 | 61.7 – 670 |
| $N_{\text{Cha}}$ | 387 | 140 – 1110 | 234 | 110 – 563 |
| $N_{\text{Vin}}$ | 1070 | 729 – 1870 | 834 | 635 – 1250 |
| $N_{\text{AMH}}$ | 52100 | 41500 – 65200 | 29100 | 26600 – 32200 |
| $N_{\text{Yor}}$ | 3380 | 2800 – 4490 | 16900 | 10200 – 52400 |
| $N_{\text{Los}}$ | 459 | 392 – 549 | 923 | 785 – 1310 |
| $m_{\text{N–Den}}$ ($\times 10^{-5}$) | 0.745 | 0.536 – 0.956 | 0.258 | 0.0407 – 0.543 |
| $m_{\text{Yor–Los}}$ ($\times 10^{-5}$) | 40.8 | 32 – 48.6 | 9.86 | 2.62 – 16.4 |
| $\gamma_{\text{Vin}\to\text{Los}}$ | 0.0869 | 0.0624 – 0.113 | 0.0247 | 0.0129 – 0.0378 |
| $T_{\text{AMH}\to\text{AN}}$ (kya) | 250 | — | 250 | — |
| $T_{\text{AMH}\to\text{WN}}$ (kya) | 110 | — | 110 | — |
| $\gamma_{\text{AMH}\to\text{AN}}$ | | | 0.0874 | 0.0629 – 0.114 |
| $\gamma_{\text{AMH}\to\text{WN}}$ | | | 0.0115 | 0.00613 – 0.0172 |

Table S4: A summary of parameter MLE and 95% bootstrap CI for models with (Two-pulse model) and without (Null) AMH-to-Neanderthal introgressions (Bhérer dataset).

| | Null model $LL = -314$ | | S→Denisova model $LL = -300$ | |
|---|---|---|---|---|
| Parameter | MLE | 95% CI | MLE | 95% CI |
| $T_{\text{ND–AMH}}$ (kya) | 766 | 715 – 815 | 763 | 714 – 816 |
| $T_{\text{AN–Den}}$ (kya) | 726 | 624 – 760 | 757 | 592 – 761 |
| $T_{\text{AMH}\to\text{AN}}$ (ka) (fixed) | 250 | — | 250 | — |
| $T_{\text{WN–Alt}}$ (kya) | 119 | 117 – 161 | 121 | 118 – 139 |
| $T_{\text{AMH}\to\text{WN}}$ (ka) (fixed) | 110 | — | 110 | — |
| $T_{\text{Cha–Vin}}$ (kya) | 61 | 58.6 – 104 | 60.5 | 58.6 – 69.2 |
| $N_{\text{A}}$ | 17300 | 16600 – 18200 | 17100 | 16300 – 17900 |
| $N_{\text{ND}}$ | 1690 | 429 – 5980 | 311 | 469 – 8000 |
| $N_{\text{N}}$ | 2590 | 1810 – 2830 | 2760 | 2490 – 3060 |
| $N_{\text{Alt}}$ | 205 | 74 – 2810 | 332 | 116 – 1780 |
| $N_{\text{Cha}}$ | 254 | 139 – 1920 | 231 | 129 – 718 |
| $N_{\text{Vin}}$ | 851 | 681 – 2960 | 819 | 689 – 1340 |
| $N_{\text{Den}}$ | 3570 | 3030 – 4110 | 3290 | 2820 – 4010 |
| $N_{\text{Yor}}$ | 27800 | 26300 – 29400 | 28500 | 26800 – 30200 |
| $m_{\text{N–Den}}$ ($\times 10^{-5}$) | 0.27 | 0.001* – 0.567 | 0.465 | 0.001* – 0.726 |
| $\gamma_{\text{AMH}\to\text{N}}$ | 0.0882 | 0.0602 – 0.114 | 0.057 | 0.0369 – 0.076 |
| $\gamma_{\text{AMH}\to\text{WN}}$ | 0.0115 | 0.00728 – 0.0158 | 0.0113 | 0.00709 – 0.0157 |
| $T_{\text{A–S}}$ (kya) (fixed) | | | 2000 | — |
| $N_{\text{S}}$ (fixed) | | | 20000 | — |
| $\gamma_{\text{S}\to\text{Den}}$ | | | 0.0135 | 0.0027 – 0.023 |

Table S5: A summary of parameter MLE and 95% bootstrap CI for models with (S→Denisova) and without (Null) Superarchaic-to-Denisova introgression (Bhérer dataset).

| | Migration model $LL=-88$ | | Admixture model $LL=-79$ | |
|---|---|---|---|---|
| Parameter | MLE | 95% CI | MLE | 95% CI |
| $T_{\text{ND–AMH}}$ (ka) | 812 | 751 – 862 | 820 | 761 – 870 |
| $T_{\text{AMH}\to\text{Vin}}$ (ka) (fixed) | 250 | — | 250 | — |
| $T_{\text{Yor–OOA}}$ (ka) | 53.6 | 49.8 – 56.7 | 53.8 | 51.1 – 57.8 |
| $T_{\text{OOA–Stu}}$ (ka) | 52.3 | 48 – 54 | 52.7 | 48.8 – 56.1 |
| $T_{\text{Vin}\to\text{OOA}}$ (ka) (fixed) | 48 | — | 48 | — |
| $T_{\text{OOA–Ust}}$ (ka) (fixed) | 47 | — | 47 | — |
| $N_{\text{A}}$ | 16800 | 16100 – 17800 | 16800 | 16100 – 17800 |
| $N_{\text{Vin}}$ | 2350 | 2130 – 2540 | 2310 | 2120 – 2530 |
| $N_{\text{AMH}}$ | 29000 | 26600 – 31800 | 29300 | 26900 – 32400 |
| $N_{\text{Yor}}$ | 21300 | 11300 – 49900 | 19700 | 11300 – 38500 |
| $N_{\text{OOA}}$ | 94.7 | 40.1 – 362 | 77.6 | 46.1 – 353 |
| $N_{\text{Los}}$ | 957 | 817 – 1210 | 1490 | 1270 – 1730 |
| $N_{\text{Ust}}$ | 20000 | 519 – 10000 | 2120 | 545 – 20000 |
| $N_{\text{Stu}}$ | 4810 | 3350 – 6470 | 5150 | 3020 – 9720 |
| $m_{\text{Yor–Stu}}(\times 10^{-5})$ | 0.001 | 0.001* – 6.46 | 0.0406 | 0.001* – 5.72 |
| $m_{\text{Yor–Los}}(\times 10^{-5})$ | 5.67 | 1.46 – 8.58 | 6.25 | 2.9 – 9.72 |
| $\gamma_{\text{AMH}\to\text{AN}}$ | 0.133 | 0.0998 – 0.18 | 0.141 | 0.102 – 0.187 |
| $\gamma_{\text{Vin}\to\text{OOA}}$ | 0.022 | 0.0126 – 0.0315 | 0.0201 | 0.0127 – 0.0288 |
| $m_{\text{Los–Stu}}(\times 10^{-5})$ | 36.9 | 28.4 – 44.4 | | |
| $T_{\text{Los}\to\text{Stu}}$ (ka) | | | 22.5 | 8.75 – 33.6 |
| $\gamma_{\text{Los}\to\text{Stu}}$ | | | 0.435 | 0.297 – 0.693 |

Table S6: A summary of parameter MLE and 95% bootstrap CI for Loschbour-Stuttgart migration and admixture models (Bhérer dataset).

| | Bhérer map | | Zhou JHS map | |
|---|---|---|---|---|
| | $LL = -548$ | | | |
| Parameter | MLE | 95% CI | MLE | 95% CI |
| $T_{\text{A–S}}$ (ka) (fixed) | 2000 | — | 2000 | — |
| $T_{\text{ND–AMH}}$ (ka) | 798 | 748 – 827 | 748 | 721 – 772 |
| $T_{\text{N–Den}}$ (ka) | 688 | 639 – 734 | 718 | 684 – 735 |
| $T_{\text{AMH}\to\text{N}}$ (ka) (fixed) | 250 | — | 250 | — |
| $T_{\text{WN–Alt}}$ (ka) | 123 | 117 – 137 | 118 | 117 – 125 |
| $T_{\text{AMH}\to\text{WN}}$ (ka) (fixed) | 110 | — | 110 | — |
| $T_{\text{Cha–Vin}}$ (ka) | 60.5 | 57.3 – 67.7 | 61.1 | 57.9 – 64.4 |
| $T_{\text{Yor–OOA}}$ (ka) | 56.9 | 53.6 – 60 | 59.3 | 53.4 – 60.1 |
| $T_{\text{OOA–Stu}}$ (ka) | 54.7 | 49.7 – 57.6 | 56.8 | 50.8 – 58.2 |
| $T_{\text{Vin}\to\text{OOA}}$ (ka) (fixed) | 48 | — | 48 | — |
| $T_{\text{OOA–Ust}}$ (ka) (fixed) | 47 | — | 47 | — |
| $T_{\text{Los}\to\text{Stu}}$ (ka) | 29.4 | 13.2 – 35.9 | 28.4 | 12.5 – 35.3 |
| $N_{\text{A}}$ | 16500 | 15900 – 17300 | 17200 | 16700 – 17700 |
| $N_{\text{S}}$ (fixed) | 20000 | — | 20000 | — |
| $N_{\text{ND}}$ | 5060 | 3540 – 5760 | 1940 | 1940 – 2560 |
| $N_{\text{N}}$ | 2800 | 2530 – 3090 | 2640 | 2420 – 2920 |
| $N_{\text{AMH}}$ | 30100 | 27700 – 31800 | 28700 | 26300 – 30000 |
| $N_{\text{OOA}}$ | 176 | 99.3 – 331 | 205 | 68.6 – 222 |
| $N_{\text{Alt}}$ | 402 | 94.6 – 1480 | 150 | 67.1 – 640 |
| $N_{\text{Cha}}$ | 237 | 83.2 – 602 | 292 | 123 – 521 |
| $N_{\text{Vin}}$ | 829 | 592 – 1220 | 872 | 659 – 1110 |
| $N_{\text{Den}}$ | 3270 | 2870 – 3730 | 3120 | 2870 – 3590 |
| $N_{\text{Los}}$ | 1530 | 1340 – 1840 | 1530 | 1360 – 1860 |
| $N_{\text{Ust}}$ | 3110 | 584 – 100000* | 100000 | 701 – 100000* |
| $N_{\text{Stu}}$ | 7060 | 4160 – 13900 | 6390 | 3780 – 13500 |
| $N_{\text{Yor}}$ | 22900 | 15600 – 59600 | 24800 | 17800 – 85700 |
| $m_{\text{N–Den}}$ ($\times 10^{-5}$) | 0.393 | 0.0866 – 0.685 | 0.618 | 0.306 – 0.733 |
| $m_{\text{Yor–Los}}$ ($\times 10^{-5}$) | 6.24 | 2.22 – 8.98 | 6.81 | 2.02 – 9.38 |
| $\gamma_{\text{S}\to\text{Den}}$ | 0.0149 | 0.00748 – 0.0265 | 0.0131 | 0.00286 – 0.0235 |
| $\gamma_{\text{AMH}\to\text{AN}}$ | 0.054 | 0.0282 – 0.0702 | 0.0583 | 0.039 – 0.0814 |
| $\gamma_{\text{AMH}\to\text{WN}}$ | 0.0115 | 0.00651 – 0.0166 | 0.0112 | 0.00684 – 0.017 |
| $\gamma_{\text{Vin}\to\text{OOA}}$ | 0.0202 | 0.012 – 0.0274 | 0.0199 | 0.0115 – 0.0265 |
| $\gamma_{\text{Los}\to\text{Stu}}$ | 0.575 | 0.333 – 0.752 | 0.616 | 0.362 – 0.794 |

Table S7: A summary of parameter MLE and 95% bootstrap CI for the focal model (Bhérer and Zhou datasets). The log-likelihood of the focal model fitted to the Bhérer dataset is provided for comparisons with results in Tables S8, S9.

| Parameter | $\mu = 1.1 \times 10^{-8}$ $LL = -609$ MLE | 95% CI | $\mu = 1.5 \times 10^{-8}$ $LL = -630$ MLE | 95% CI |
|---|---|---|---|---|
| $T_{\text{A–S}}$ (ka) (fixed) | 2000 | — | 2000 | — |
| $T_{\text{ND–AMH}}$ (ka) | 921 | 879 – 946 | 700 | 654 – 731 |
| $T_{\text{N–Den}}$ (ka) | 856 | 819 – 882 | 615 | 548 – 650 |
| $T_{\text{AMH}\to\text{AN}}$ (ka) (fixed) | 250 | — | 250 | — |
| $T_{\text{WN–Alt}}$ (ka) | 136 | 126 – 152 | 116 | 115 – 120 |
| $T_{\text{AMH}\to\text{WN}}$ (ka) (fixed) | 110 | — | 110 | — |
| $T_{\text{Cha–Vin}}$ (ka) | 68.4 | 60.6 – 95.9 | 59.5 | 57.3 – 63.6 |
| $T_{\text{Yor–OOA}}$ (ka) | 57.7 | 53.1 – 59.4 | 57.2 | 52.5 – 58 |
| $T_{\text{OOA–Stu}}$ (ka) | 56.2 | 51.6 – 58 | 53.6 | 48.6 – 55.6 |
| $T_{\text{Vin}\to\text{OOA}}$ (ka) (fixed) | 48 | — | 48 | — |
| $T_{\text{OOA–Ust}}$ (ka) (fixed) | 47 | — | 47 | — |
| $T_{\text{Los}\to\text{Stu}}$ (ka) | 30.2 | 15 – 36.4 | 28.2 | 10.1 – 35.6 |
| $N_{\text{A}}$ | 19600 | 19100 – 20400 | 14200 | 13700 – 15000 |
| $N_{\text{S}}$ (fixed) | 20000 | — | 20000 | — |
| $N_{\text{ND}}$ | 3280 | 3030 – 3580 | 5150 | 3690 – 6960 |
| $N_{\text{N}}$ | 3460 | 2920 – 3680 | 2280 | 2080 – 2540 |
| $N_{\text{AMH}}$ | 36 800 | 33 900 – 38 800 | 25 500 | 23 400 – 26 700 |
| $N_{\text{OOA}}$ | 113 | 83.1 – 124 | 285 | 114 – 328 |
| $N_{\text{Alt}}$ | 935 | 453 – 1700 | 57.3 | 21 – 335 |
| $N_{\text{Cha}}$ | 647 | 234 – 2050 | 184 | 79.7 – 398 |
| $N_{\text{Vin}}$ | 1290 | 747 – 2950 | 778 | 590 – 1050 |
| $N_{\text{Den}}$ | 3780 | 3290 – 4080 | 2610 | 2350 – 3070 |
| $N_{\text{Los}}$ | 1510 | 1340 – 1810 | 1550 | 1370 – 1880 |
| $N_{\text{Ust}}$ | 3530 | 565 – 100000* | 3670 | 562 – 100000* |
| $N_{\text{Stu}}$ | 8190 | 4860 – 17700 | 6220 | 3600 – 12200 |
| $N_{\text{Yor}}$ | 20500 | 15000 – 49100 | 25800 | 17700 – 88500 |
| $m_{\text{N–Den}}$ ($\times 10^{-5}$) | 0.398 | 0.277 – 0.62 | 0.724 | 0.279 – 0.985 |
| $m_{\text{Yor–Los}}$ ($\times 10^{-5}$) | 7.14 | 2.98 – 9.47 | 5.56 | 0.935 – 8.03 |
| $\gamma_{\text{S}\to\text{Den}}$ | 0.0312 | 0.0186 – 0.0459 | 0.0116 | 0.00404 – 0.0201 |
| $\gamma_{\text{AMH}\to\text{AN}}$ | 0.0365 | 0.0162 – 0.0563 | 0.0501 | 0.0219 – 0.0761 |
| $\gamma_{\text{AMH}\to\text{WN}}$ | 0.00528 | 0.000686 – 0.0102 | 0.0179 | 0.0128 – 0.0231 |
| $\gamma_{\text{Vin}\to\text{OOA}}$ | 0.0222 | 0.0136 – 0.0294 | 0.019 | 0.0107 – 0.0255 |
| $\gamma_{\text{Los}\to\text{Stu}}$ | 0.602 | 0.37 – 0.78 | 0.546 | 0.305 – 0.747 |

Table S8: A summary of parameter MLE and 95% bootstrap CI of the full model, fitted with a low ($1.1 \times 10^{-8}$ generation$^{-1}$ bp$^{-1}$) and high ($1.5 \times 10^{-8}$ generation$^{-1}$ bp$^{-1}$) fixed mutation rate parameter (Bhérer dataset).

| | With S→Denisova | | Without S→Denisova | |
|---|---|---|---|---|
| | $LL = -548$ | | $LL = -576$ | |
| Parameter | MLE | 95% CI | MLE | 95% CI |
| $T_{\text{A–S}}$ (ka) (fixed) | 2000 | — | 2000 | — |
| $T_{\text{ND–AMH}}$ (ka) | 798 | 748 – 827 | 792 | 752 – 847 |
| $T_{\text{N–Den}}$ (ka) | 688 | 639 – 734 | 678 | 640 – 751 |
| $T_{\text{AMH}\to\text{AN}}$ (ka) (fixed) | 250 | — | 250 | — |
| $T_{\text{WN–Alt}}$ (ka) | 123 | 117 – 137 | 121 | 116 – 132 |
| $T_{\text{AMH}\to\text{WN}}$ (ka) (fixed) | 110 | — | 110 | — |
| $T_{\text{Cha–Vin}}$ (ka) | 60.5 | 57.3 – 67.7 | 60.7 | 56.7 – 70.5 |
| $T_{\text{Yor–OOA}}$ (ka) | 56.9 | 53.6 – 60 | 58 | 52.5 – 60.6 |
| $T_{\text{OOA–Stu}}$ (ka) | 54.7 | 49.7 – 57.6 | 55.7 | 49.6 – 58.7 |
| $T_{\text{Vin}\to\text{OOA}}$ (ka) (fixed) | 48 | — | 48 | — |
| $T_{\text{OOA–Ust}}$ (ka) (fixed) | 47 | — | 47 | — |
| $T_{\text{Los}\to\text{Stu}}$ (ka) | 29.4 | 13.2 – 35.9 | 31 | 11 – 36 |
| $N_{\text{A}}$ | 16500 | 15900 – 17300 | 16900 | 16000 – 17500 |
| $N_{\text{S}}$ (fixed) | 20000 | — | 20000 | — |
| $N_{\text{ND}}$ | 5060 | 3540 – 5760 | 4240 | 2750 – 5360 |
| $N_{\text{N}}$ | 2800 | 2530 – 3090 | 2670 | 2240 – 2870 |
| $N_{\text{AMH}}$ | 30100 | 27700 – 31800 | 29000 | 27000 – 31100 |
| $N_{\text{OOA}}$ | 176 | 99.3 – 331 | 191 | 68.2 – 309 |
| $N_{\text{Alt}}$ | 402 | 94.6 – 1480 | 338 | 50 – 1180 |
| $N_{\text{Cha}}$ | 237 | 83.2 – 602 | 245 | 66.2 – 821 |
| $N_{\text{Vin}}$ | 829 | 592 – 1220 | 845 | 573 – 1480 |
| $N_{\text{Den}}$ | 3270 | 2870 – 3730 | 3680 | 3130 – 3940 |
| $N_{\text{Los}}$ | 1530 | 1340 – 1840 | 1530 | 1330 – 1840 |
| $N_{\text{Ust}}$ | 3110 | 584 – 100000* | 2790 | 544 – 100000* |
| $N_{\text{Stu}}$ | 7060 | 4160 – 13900 | 7470 | 4110 – 14600 |
| $N_{\text{Yor}}$ | 22900 | 15600 – 59600 | 22200 | 15600 – 56000 |
| $m_{\text{N–Den}}$ ($\times 10^{-5}$) | 0.393 | 0.0866 – 0.685 | 0.132 | 0.0208 – 0.509 |
| $m_{\text{Yor–Los}}$ ($\times 10^{-5}$) | 6.24 | 2.22 – 8.98 | 6.21 | 2.12 – 9.15 |
| $\gamma_{\text{S}\to\text{Den}}$ | 0.0149 | 0.00748 – 0.0265 | | |
| $\gamma_{\text{AMH}\to\text{AN}}$ | 0.054 | 0.0282 – 0.0702 | 0.086 | 0.0639 – 0.113 |
| $\gamma_{\text{AMH}\to\text{WN}}$ | 0.0115 | 0.00651 – 0.0166 | 0.0115 | 0.0066 – 0.0171 |
| $\gamma_{\text{Vin}\to\text{OOA}}$ | 0.0202 | 0.012 – 0.0274 | 0.0208 | 0.0122 – 0.0276 |
| $\gamma_{\text{Los}\to\text{Stu}}$ | 0.575 | 0.333 – 0.752 | 0.623 | 0.311 – 0.773 |

Table S9: A summary of parameter MLE and 95% bootstrap CI for the full model, with and without the S→Denisova introgression event (Bhérer dataset).

### 7 Supplementary Figures

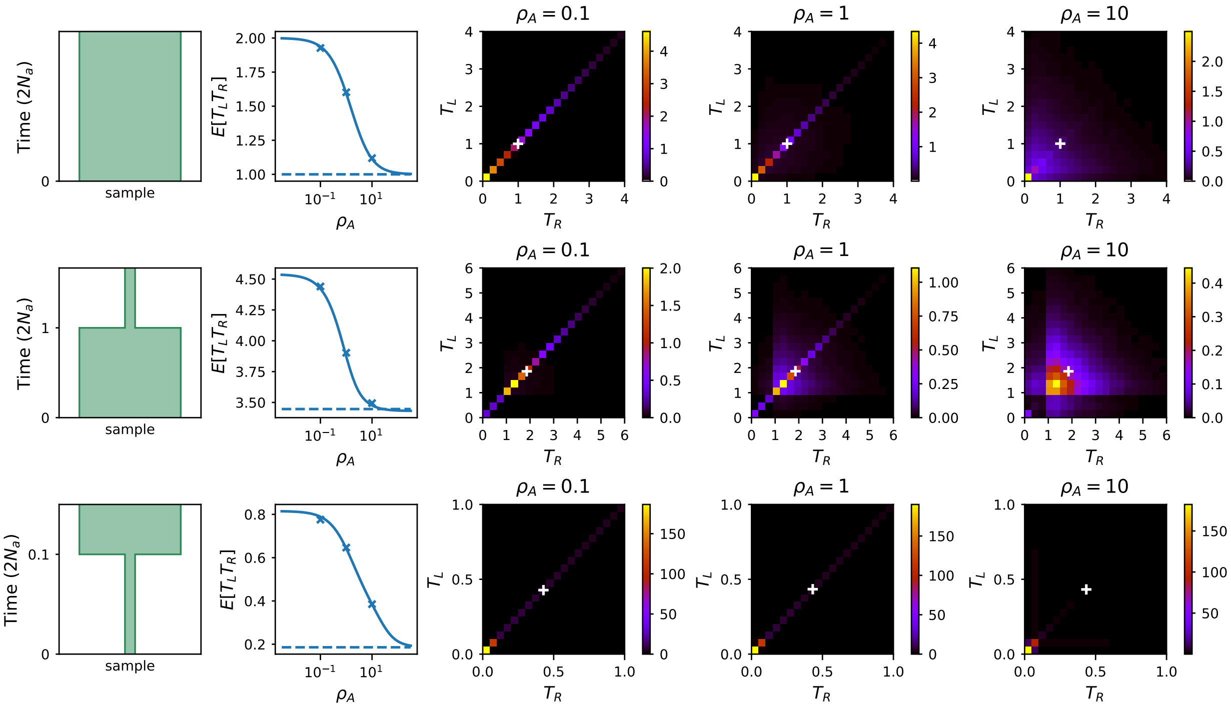

Figure S1: Joint distributions of coalescence times $T_L$, $T_R$ obtained by Monte Carlo simulation for three one-population models. In each row, the first panel from the left displays a schematic model plot. Everywhere, time is scaled by $2N_A$ ($N_A$ is the ancestral effective population size). The second panel displays the decay of $\mathbb{E}[T_LT_R]$ across $\rho_A = 4N_Ar$. $\mathbb{E}[T_LT_R]$ is directly proportional to $\mathbb{E}[H_2]$. Average $\overline{T_LT_R}$ at three focal $\rho_A$ are marked by X, and a horizontal line shows the value of $\mathbb{E}[T_L]^2 = \mathbb{E}[T_R]^2$. The last three panels display joint distributions of $T_L$, $T_R$ at focal $\rho_A$. White crosses mark average marginal times to coalescence $\bar{T}_L$, $\bar{T}_R$.

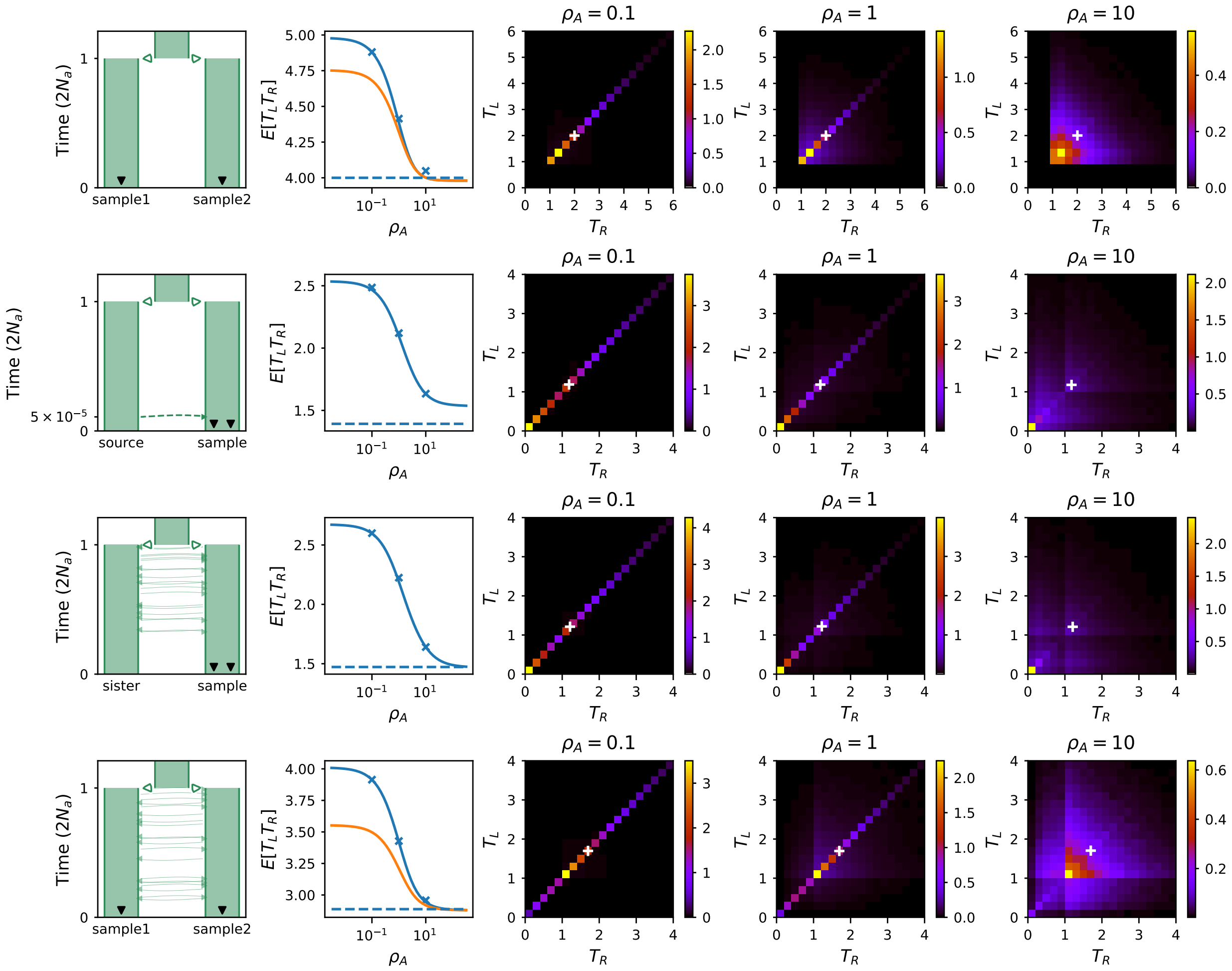

Figure S2: Joint distributions of coalescence times $T_L$, $T_R$ obtained by Monte Carlo simulation for four multi-population models. The format is as in Figure S1. In the first panel of each row, black triangles mark the sources of sampled haplotypes. Phased $\mathbb{E}[T_L T_R]$ is shown in blue and unphased $\mathbb{E}[T_L T_R]$ in orange where the distinction is meaningful; unphased $\mathbb{E}[T_L T_R]$ is proportional to $\mathbb{E}[H_{2\ \text{unphased}}(\text{sample1},\ \text{sample2})]$. In migration models, $M = 1$, while $\gamma = 0.1$ in the introgression model.

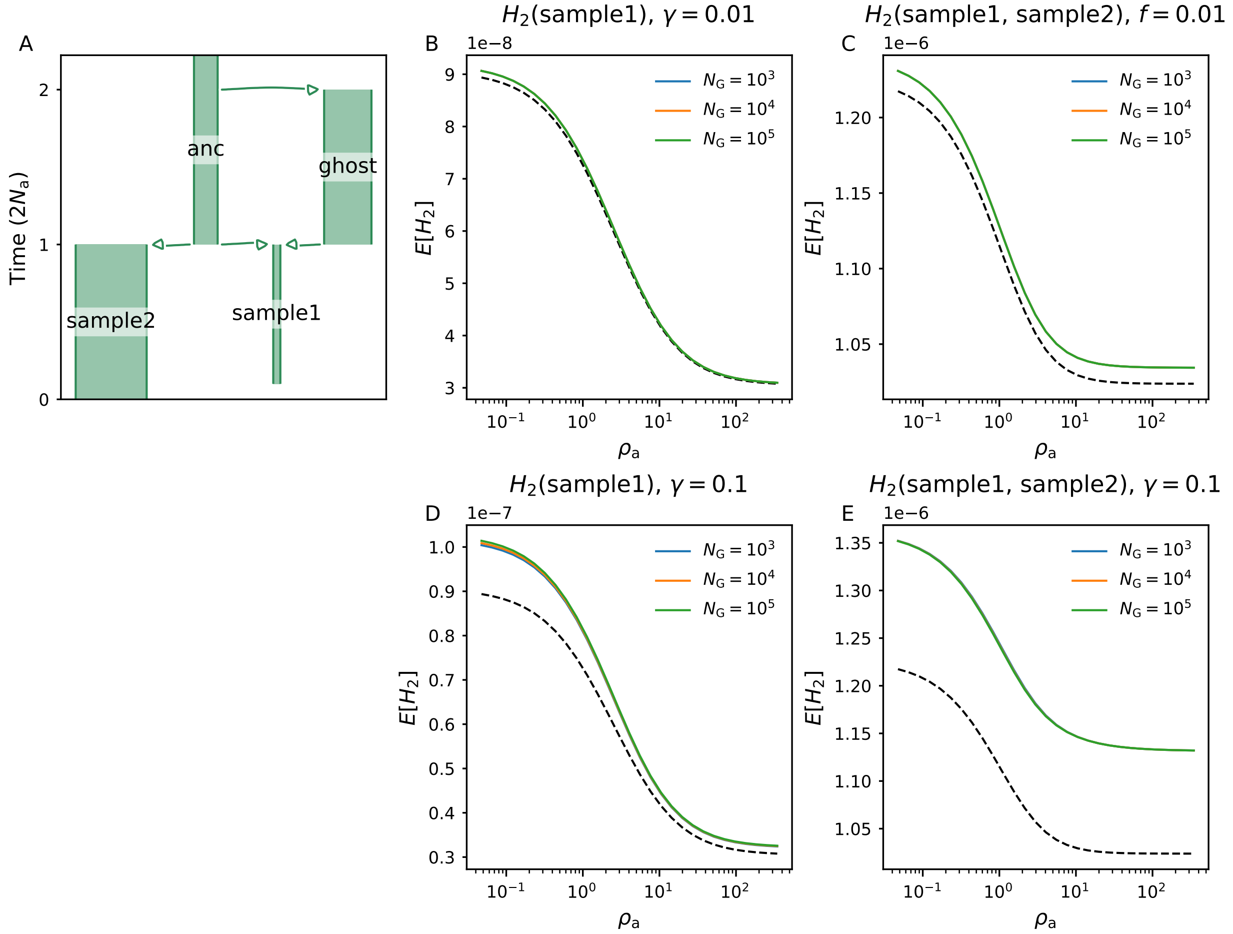

Figure S3: The effective size $N_G$ of an unsampled introgression source (ghost lineage) has little impact on one- or between-population $\mathbb{E}[H_2]$ in sampled demes, when the introgression proportion ($\gamma$) is held constant. Dotted curves show expectations in the absence of introgression ($\gamma = 0$).

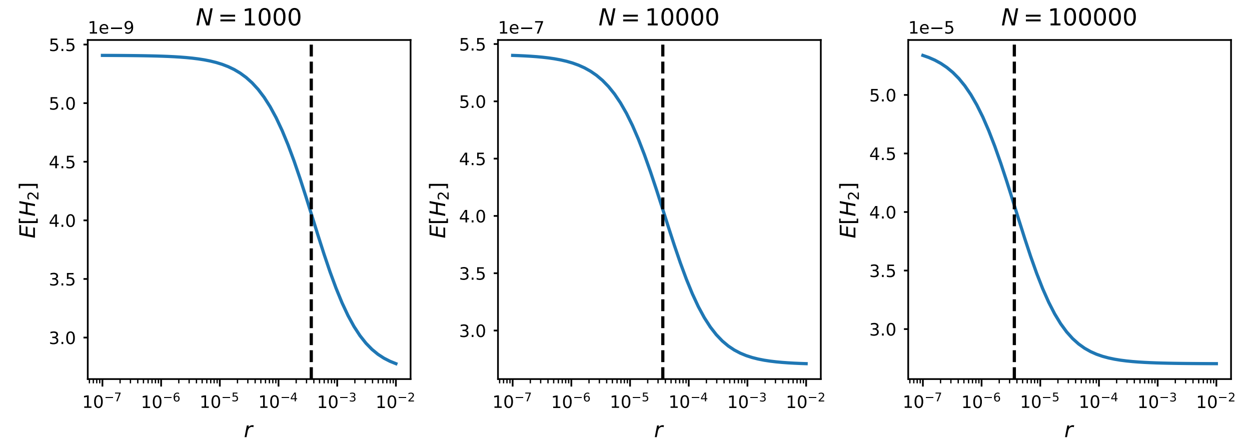

Figure S4: Equilibrium $N_e$ influences the domain of $r$ where $\mathbb{E}[H_2]$ decays to its asymptote ($\theta^2$). Dotted vertical lines mark the point where the $\mathbb{E}[H_2]$ curve decays to $3\theta^2/2$, which is at $\rho \approx 1.446$ or $r \approx 2.762/N_e$.

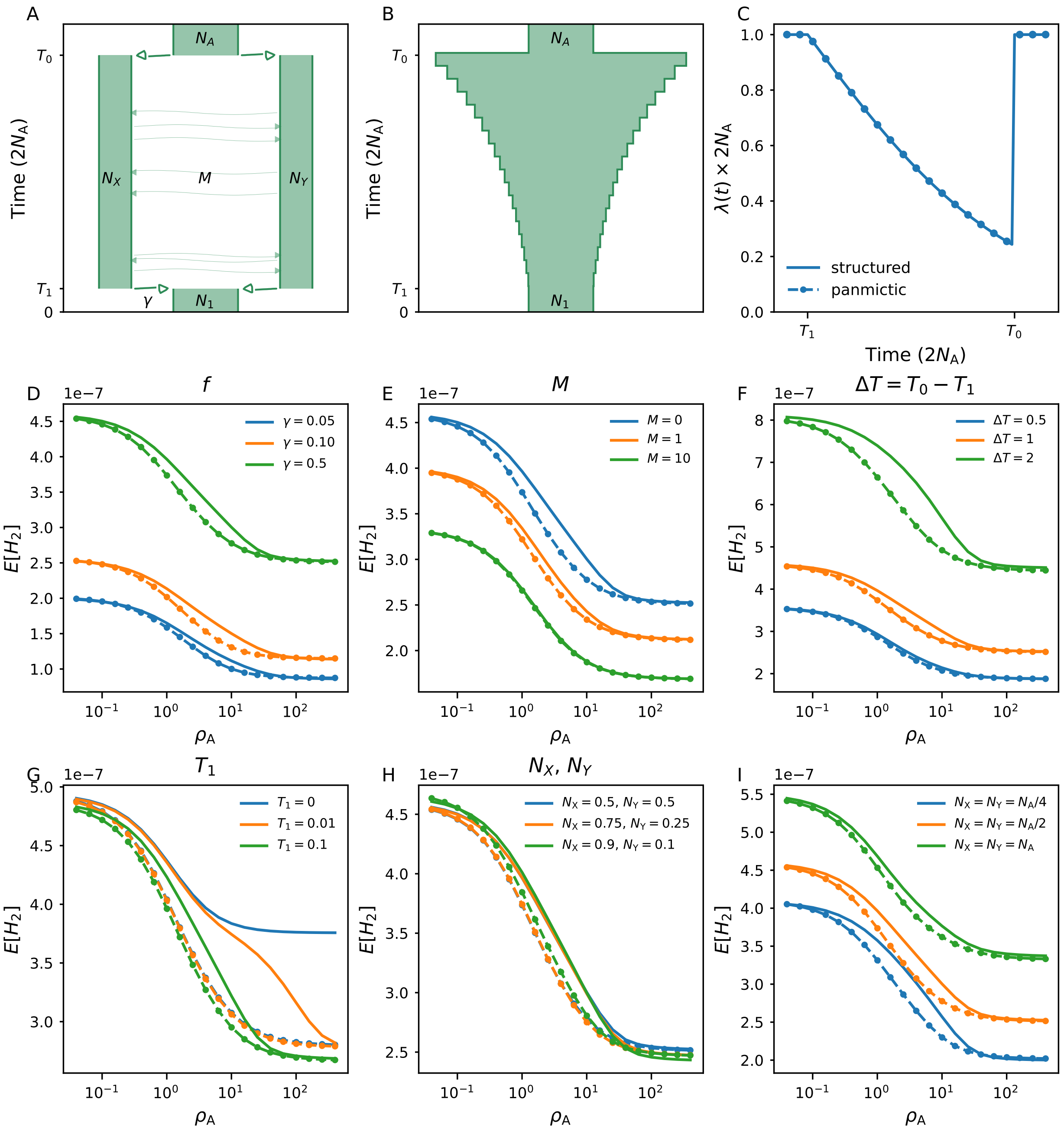

Figure S5: Various parameterizations of a simple model of ancestral structure result in different levels of $\mathbb{E}[H_2]$ discrepancy between structured models (panel A) and panmictic models (panel B) with equivalent coalescent rate profiles (panel C). Where not otherwise specified, $2N_X = 2N_Y = N_1 = N_A$, $T_1 = 0.1$, $T_0 - T_1 = 1$, $M = 0$ and $\gamma = 0.5$, where times are given in units of $2N_A$ and $N_A$ is the ancestral/reference effective population size.

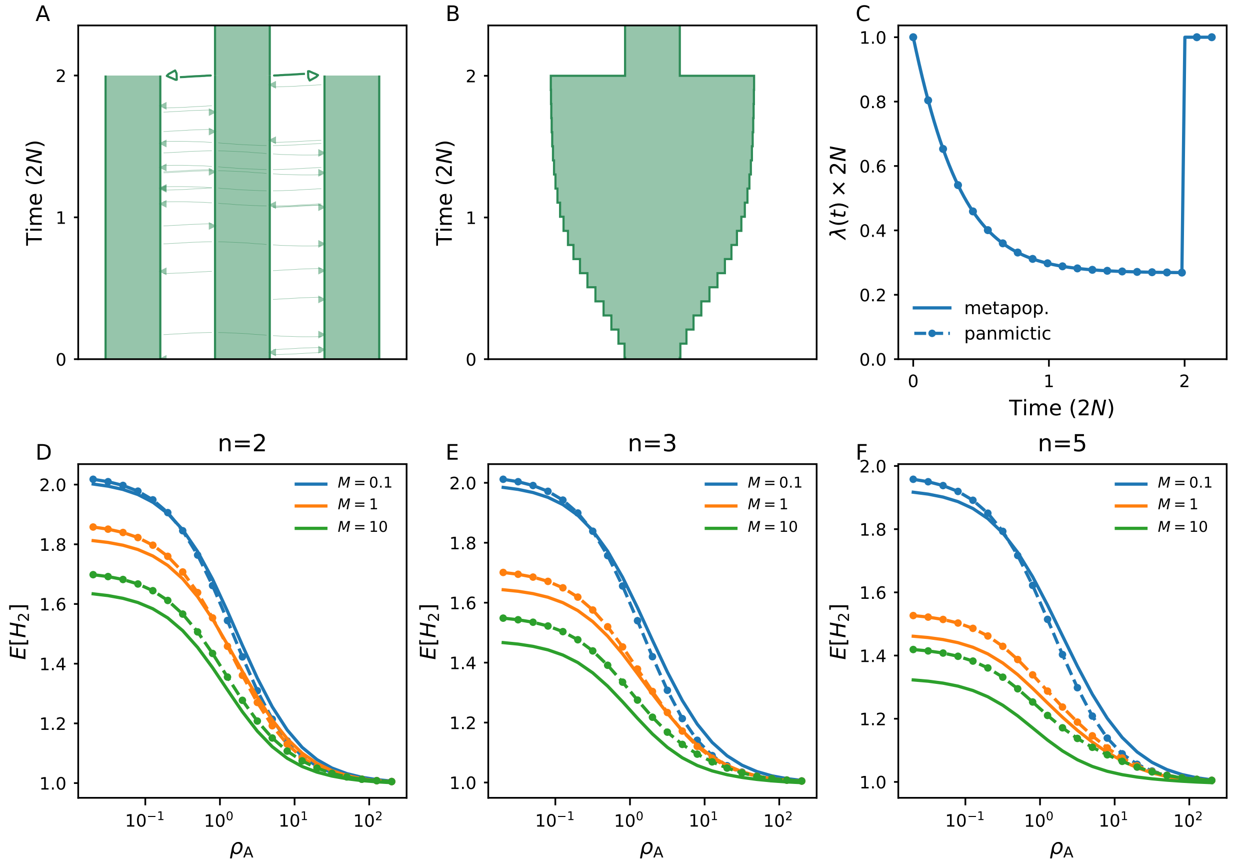

Figure S6: Metapopulation models produce $\mathbb{E}[H_2]$ curves distinct from panmictic models with identical coalescent rate profiles $\lambda(t)$ (B). This non-stationary metapopulation model family (A) is parameterized by $(N_e, n, M)$, where $N_e$ is the effective population size of each deme, $n$ is the number of demes, and $M = 4N_e m$ is the symmetric migration rate between each pair of demes. Throughout, $N_e = 5000$. (C) $\lambda(t)$ profiles, scaled by $N$, for the model with $n = 3$, $M = 1$. (D-F) Metapopulation and panmictic model expectations (solid and dotted curves) for models with several $n$ and $M$.

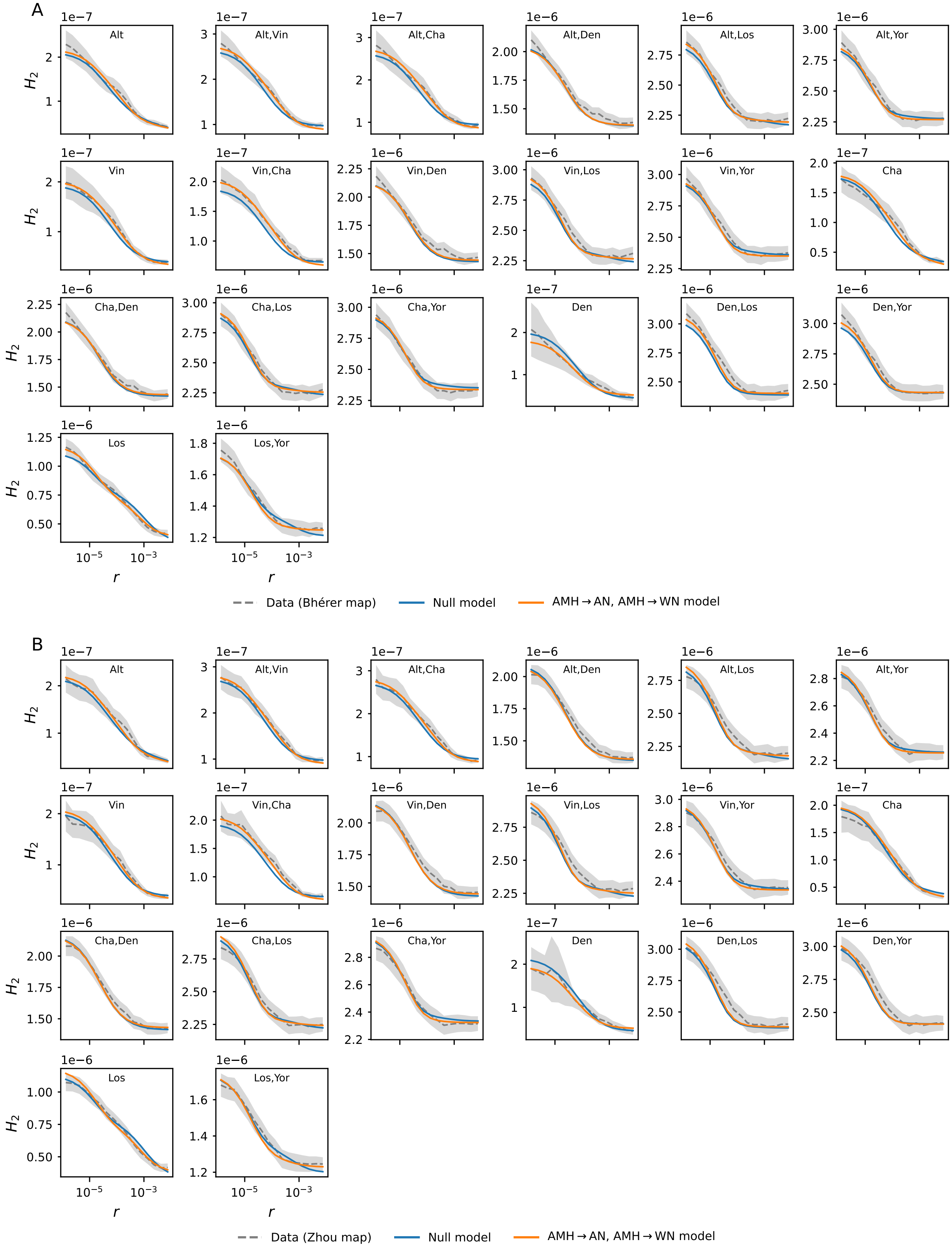

Figure S7: Expected $H_2$ under models with and without AMH-to-Neanderthal introgressions, compared to observations from Bhérer (panel A) and Zhou (panel B) datasets (shading shows 95% bootstrap CI).

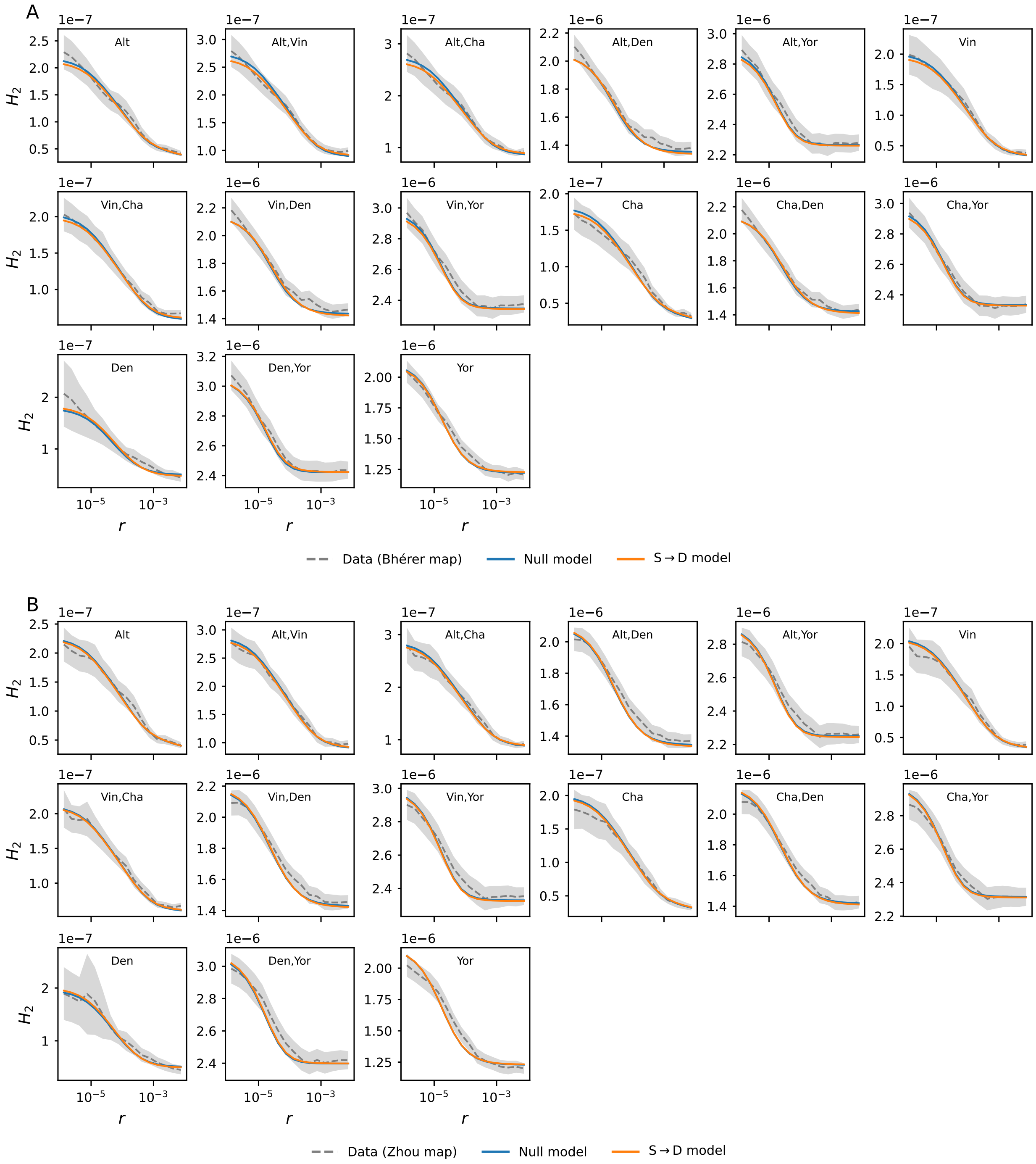

Figure S8: Expected $H_2$ under models with and without Superarchaic-to-Denisovan introgression, compared to observations from Bhérer (panel A) and Zhou (panel B) datasets (shading shows 95% bootstrap CI).

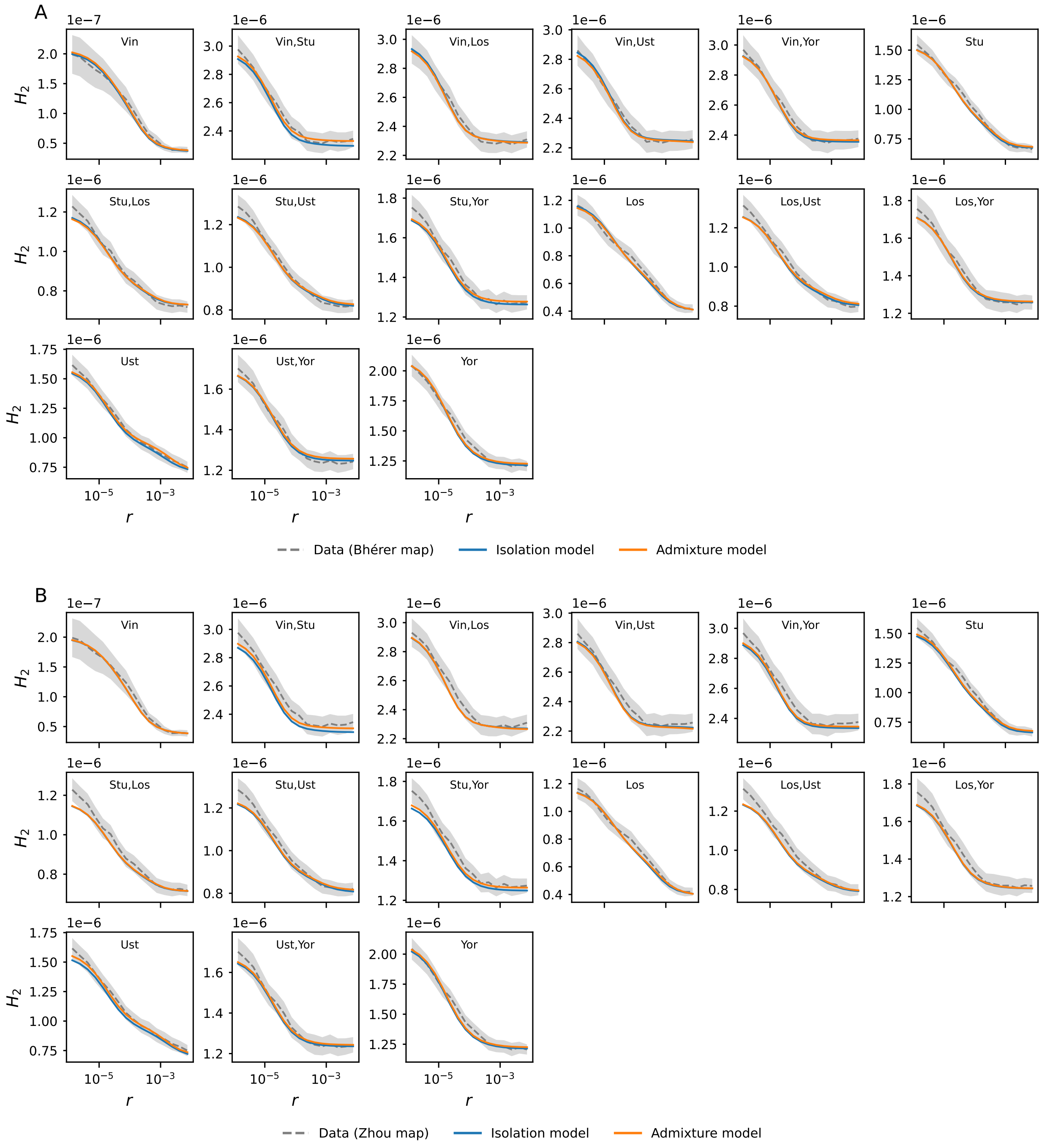

Figure S9: Expected $H_2$ under two models of gene flow between Stuttgart and Loschbour lineages, compared to observations from Bhérer (panel A) and Zhou (panel B) datasets (shading shows 95% bootstrap CI).

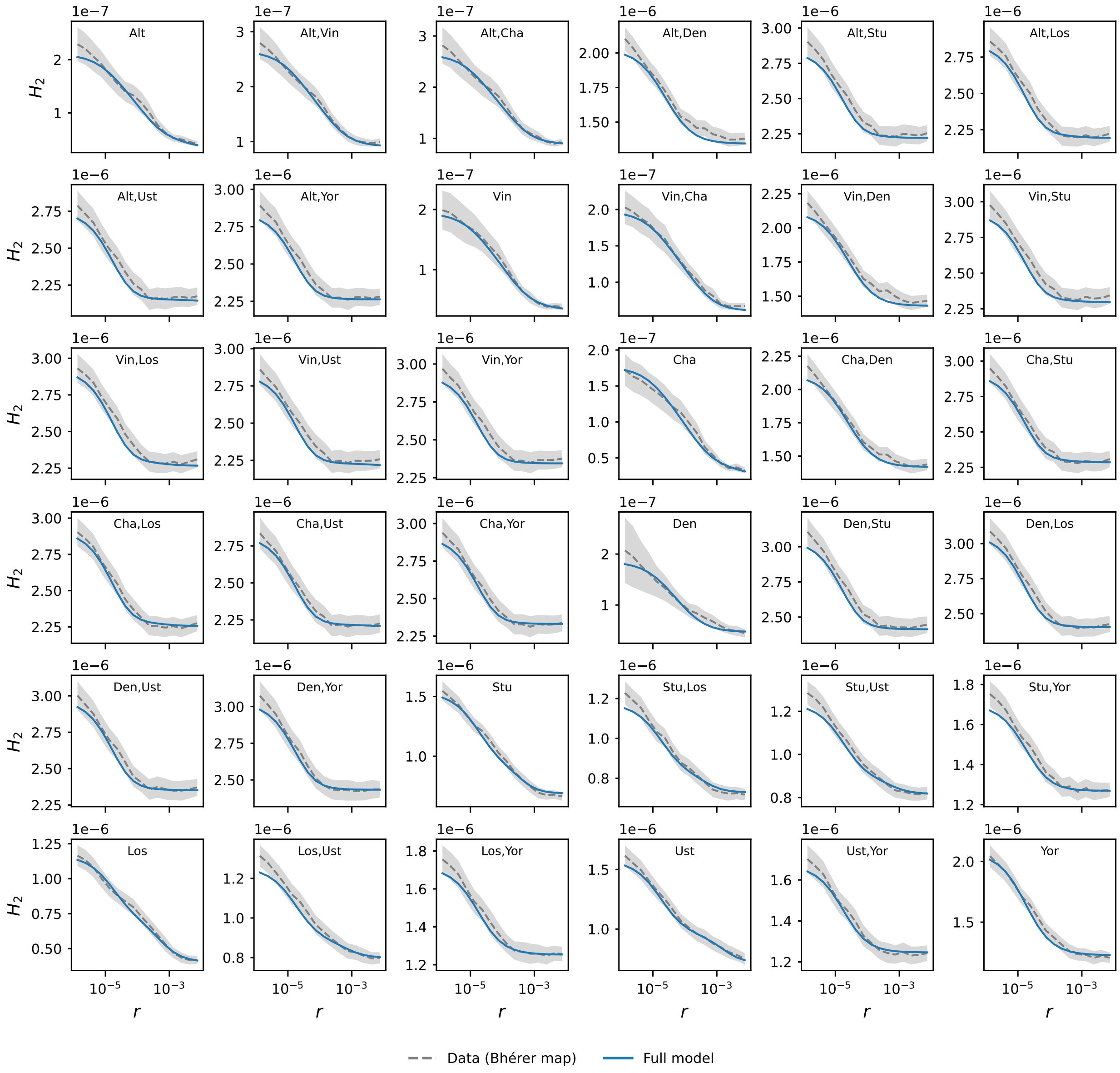

Figure S10: Expected $H_2$ under the full model compared to observations from the Bhérer dataset (shading shows 95% bootstrap CI).

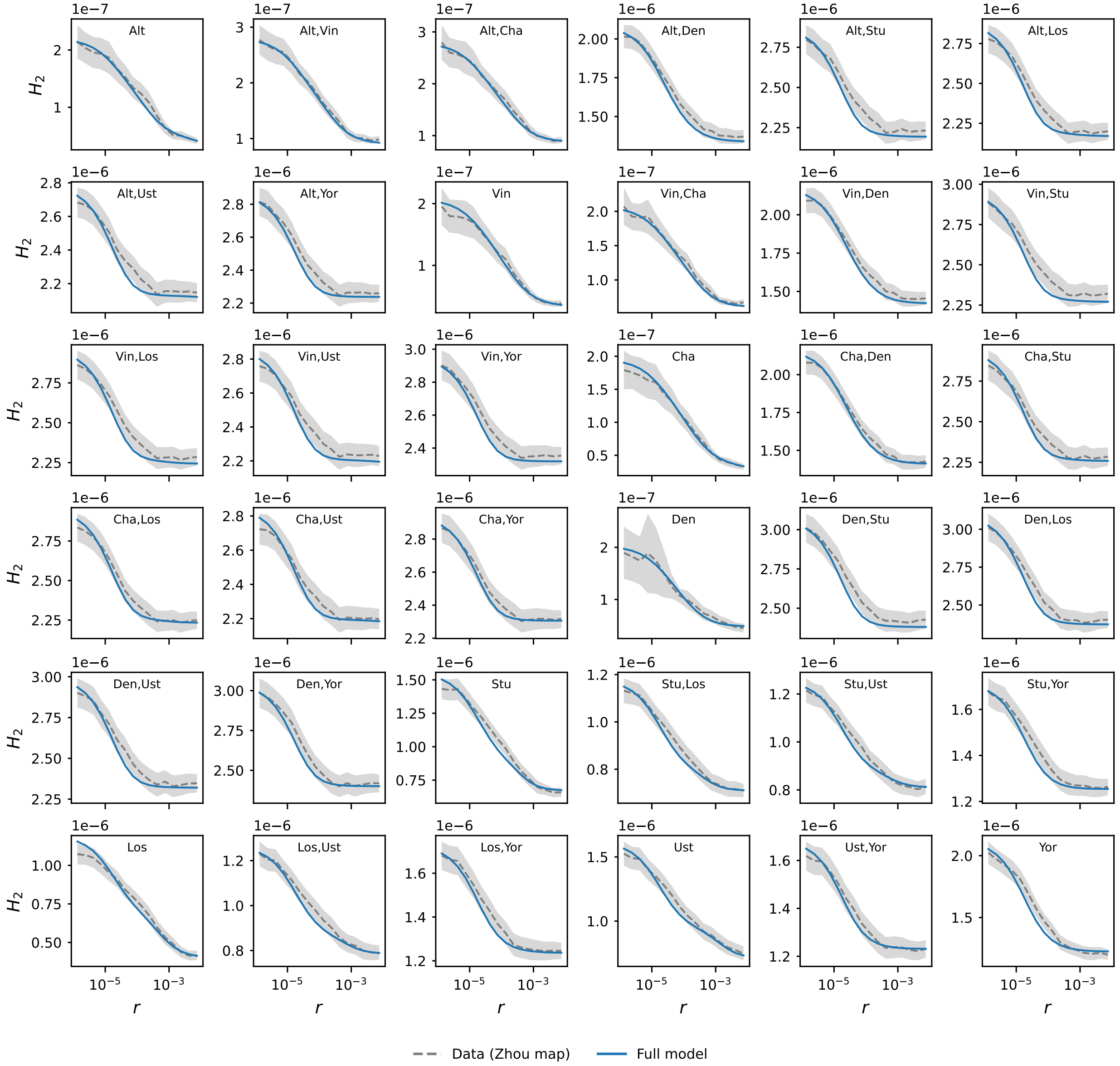

Figure S11: Expected $H_2$ under the full model compared to observations from the Zhou dataset (shading shows 95% bootstrap CI).

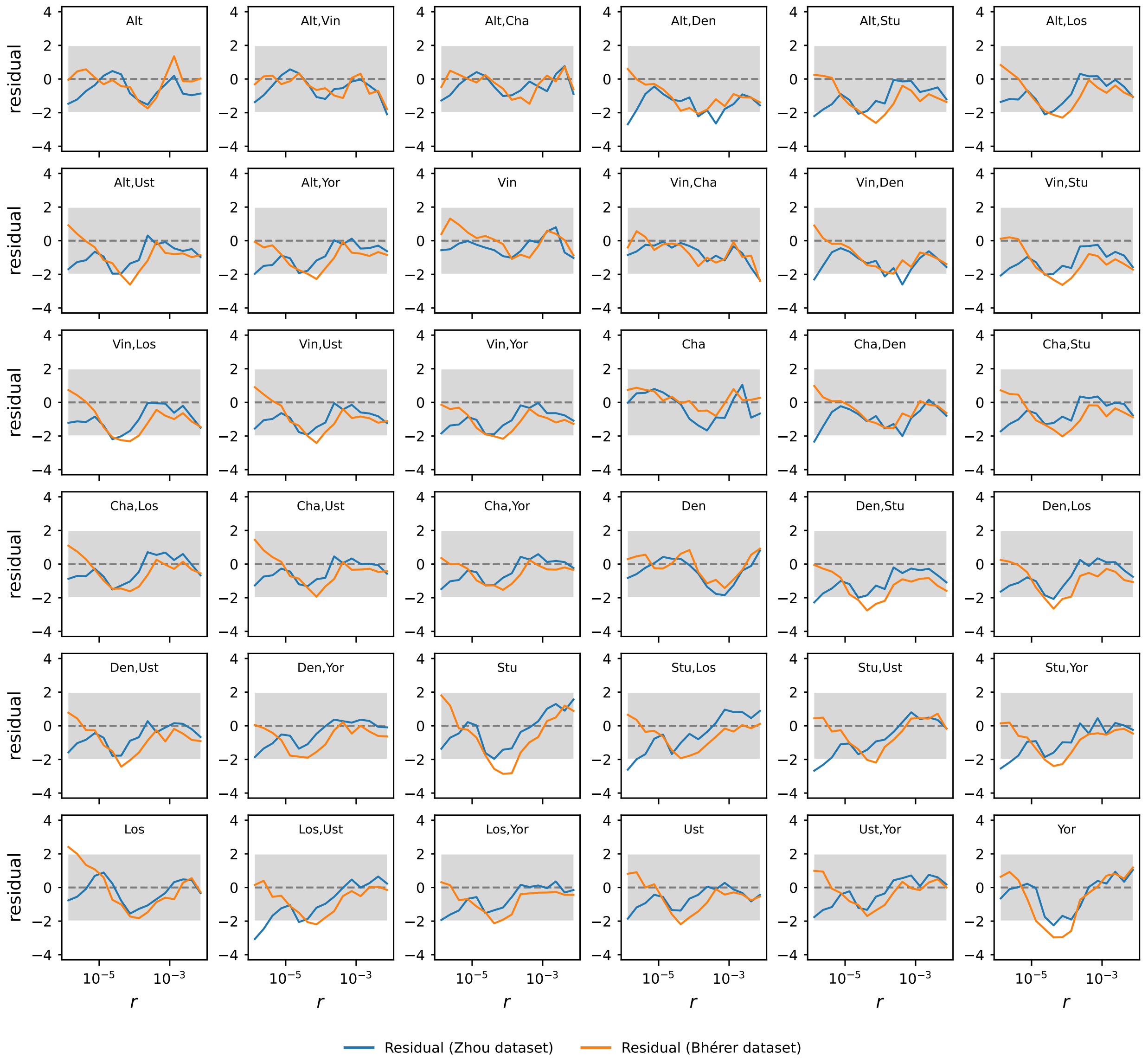

Figure S12: Scaled residuals of the full model for Bhérer and Zhou datasets. Residuals are calculated as $(\text{Model}_{i,j} - \text{Data}_{i,j})/\sqrt{V_{i,j}}$, where $V_{i,j}$ is the bootstrap variance observed for statistic $i$ in bin $j$. The shaded area corresponds to $\pm 1.96$ standard deviations.

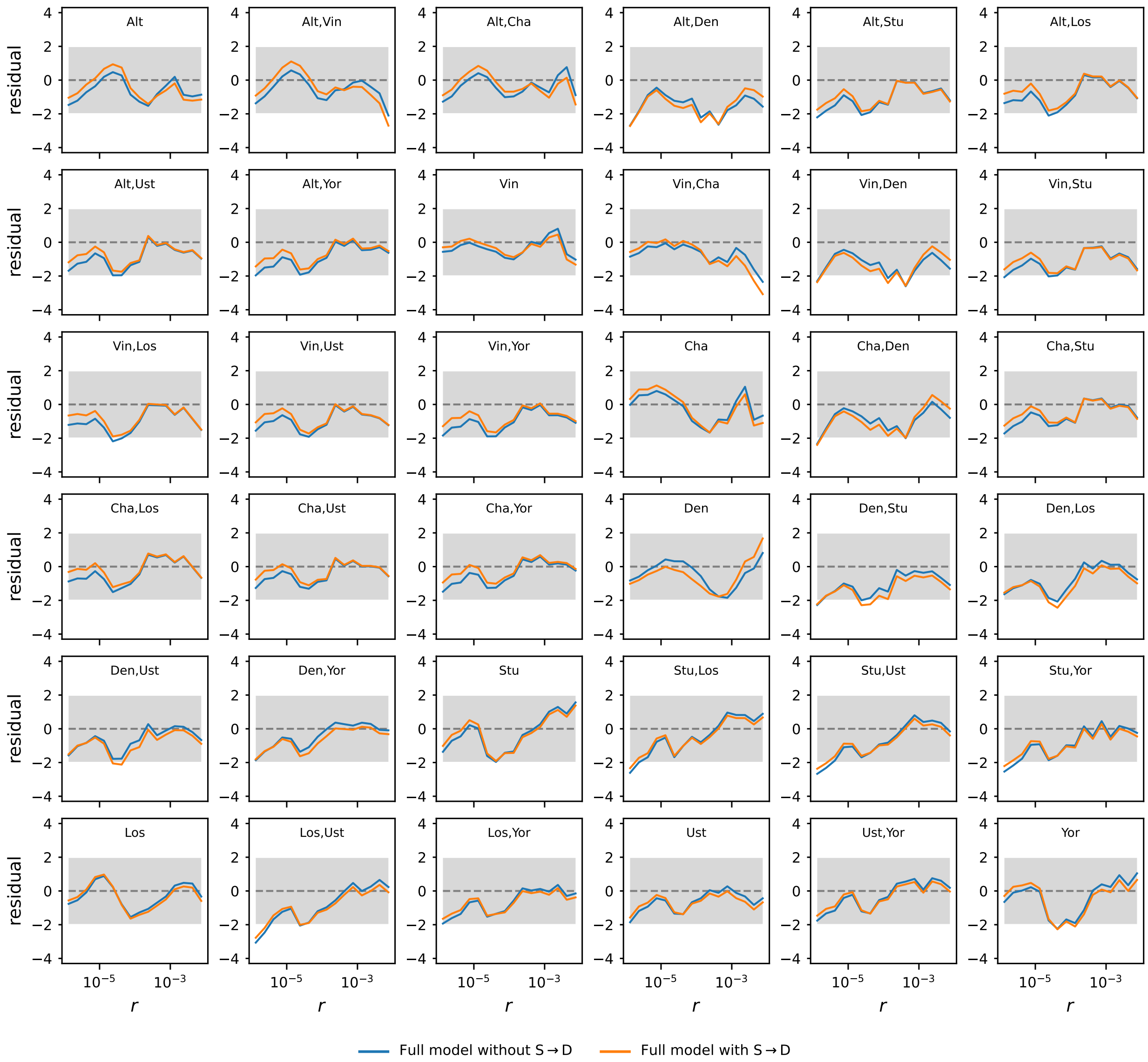

Figure S13: Scaled residuals of the full model fitted with and without Superarchaic-to-Denisovan introgression (Bhérer dataset; shaded area corresponds to ±1.96 standard deviations). Residuals are calculated as $(\text{Model}_{i,j} - \text{Data}_{i,j})/\sqrt{V_{i,j}}$, where $V_{i,j}$ is the bootstrap variance observed for statistic $i$ in bin $j$.

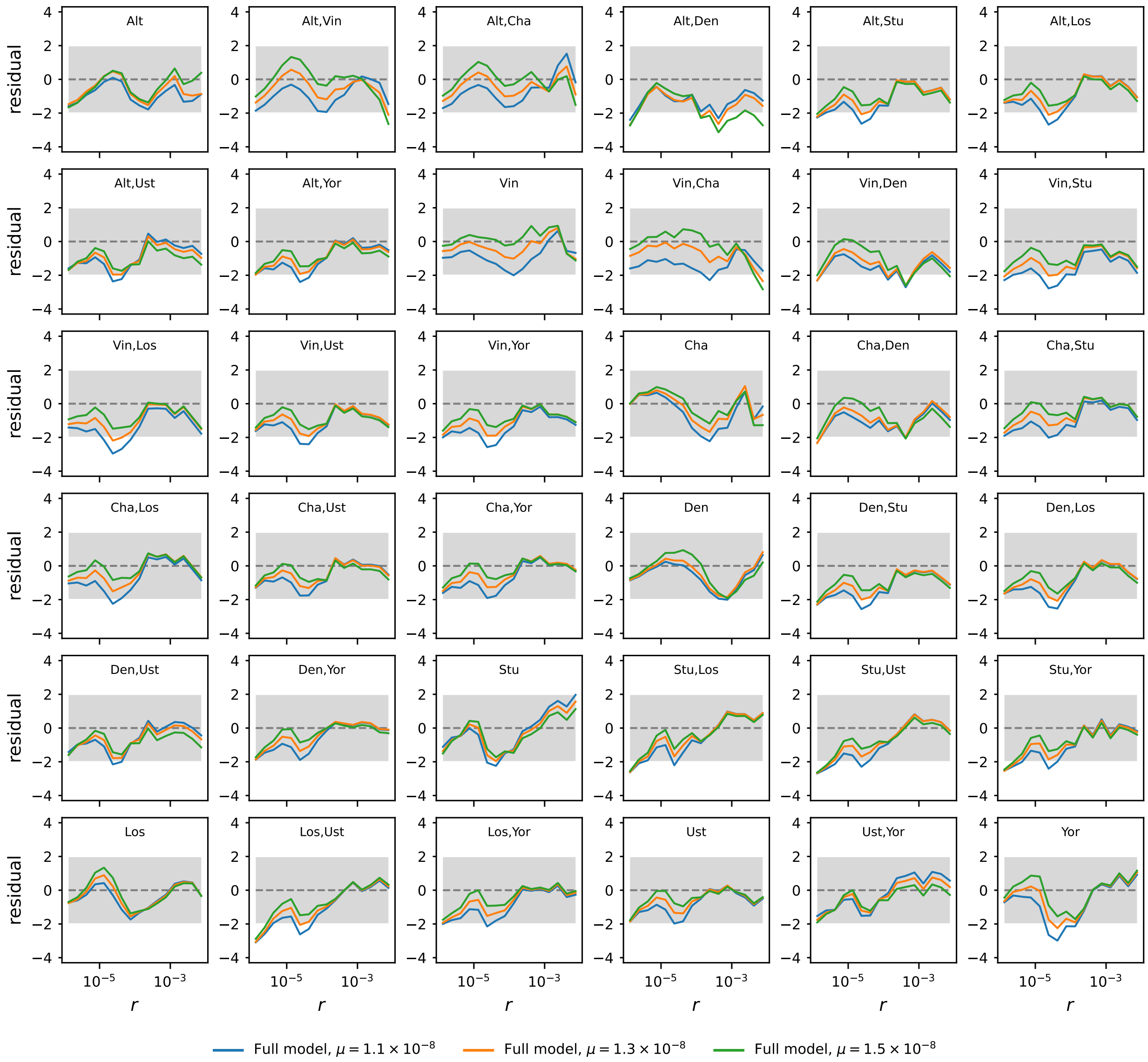

Figure S14: Residuals of the full model fitted with three different point mutation rate estimates ($1.1 \times 10^{-8}$, $1.3 \times 10^{-8}$, $1.5 \times 10^{-8}$ bp$^{-1}$ generation$^{-1}$). Residuals are calculated as $(\text{Model}_{i,j} - \text{Data}_{i,j})/\sqrt{V_{i,j}}$, where $V_{i,j}$ is the bootstrap variance observed for statistic $i$ in bin $j$. All models were fitted to the Bhérer dataset (shaded area corresponds to ±1.96 standard deviations).

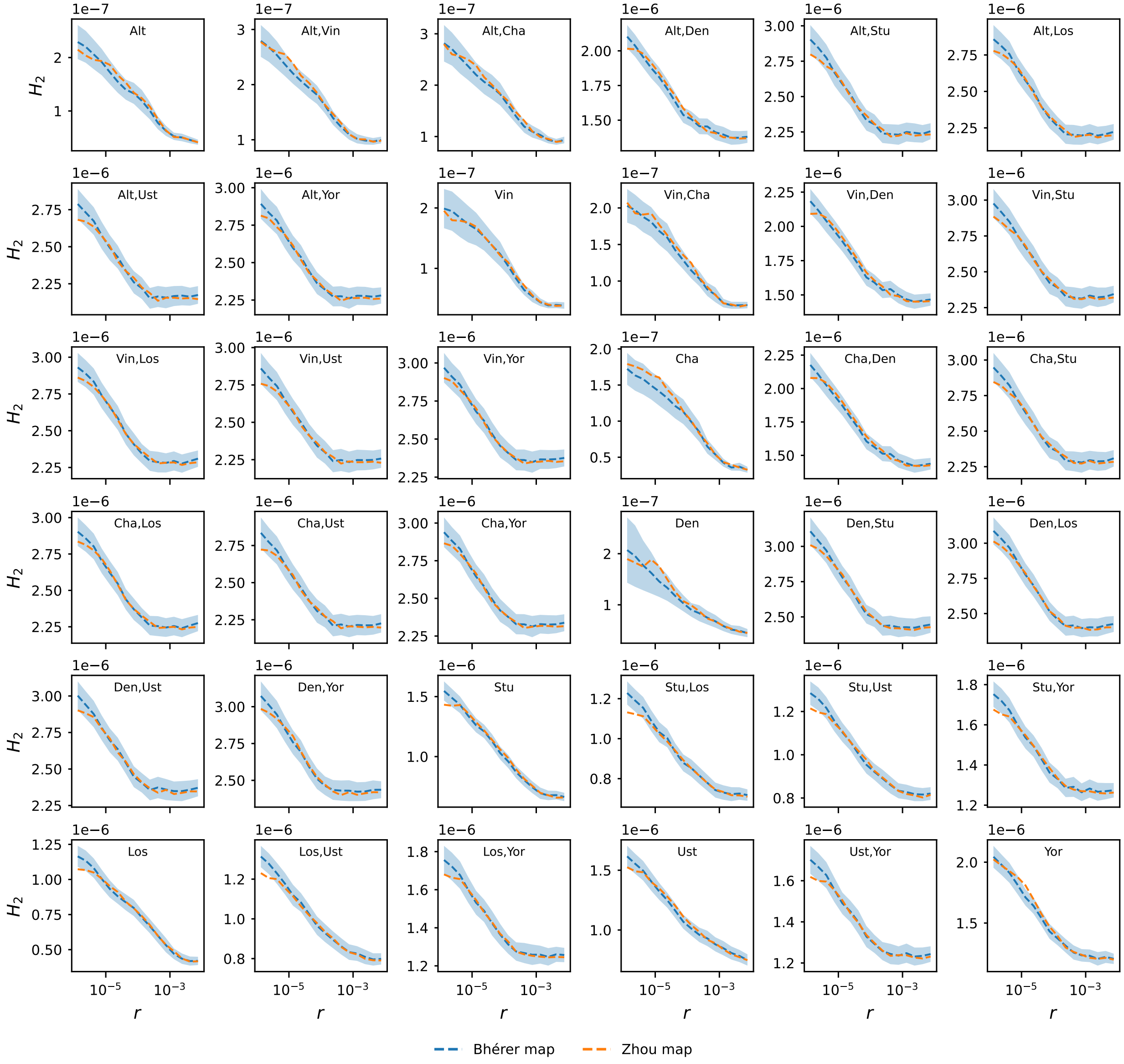

Figure S15: A direct comparison between $H_2$ datasets estimated using Bhérer and Zhou recombination maps. 95% bootstrap CI are shown for estimates made with the Bhérer map.

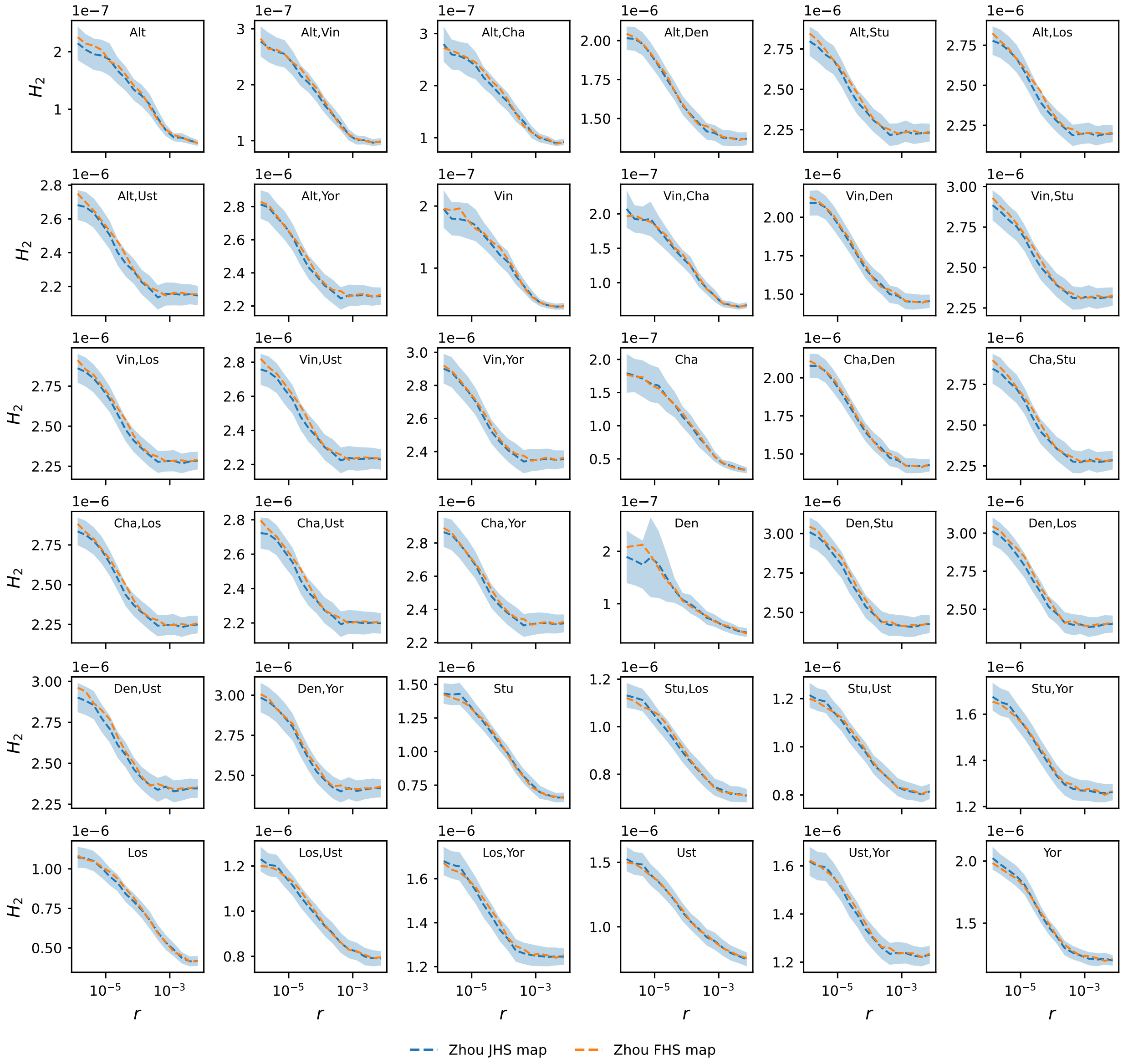

Figure S16: A comparison between $H_2$ datasets estimated with Zhou JHS (Jackson Heart Study; referred to as the *Zhou map* elsewhere in this work) and Zhou FHS (Framingham Heart Study, not used elsewhere in this work) recombination maps. 95% bootstrap CI are shown for Zhou JHS estimates.

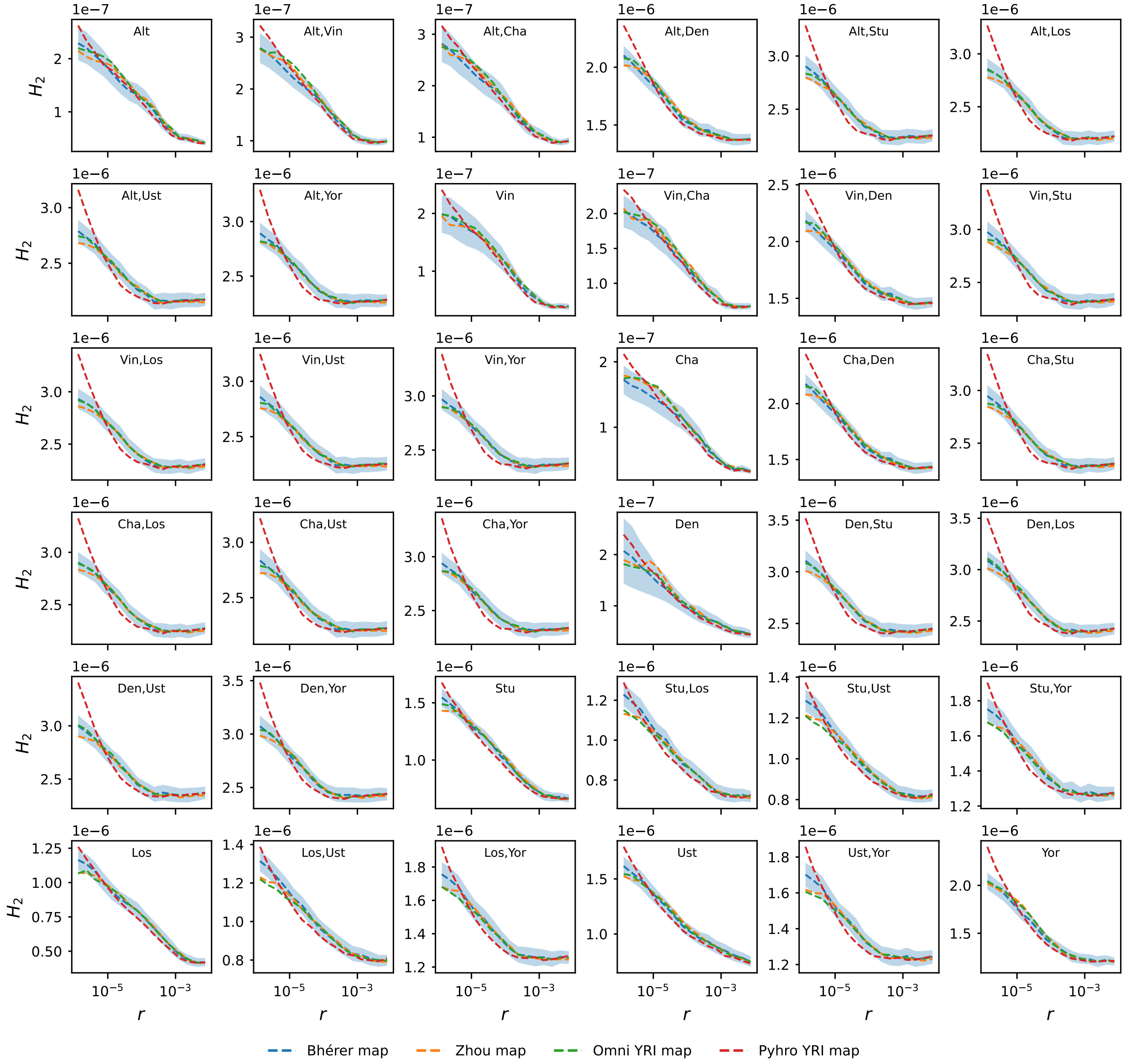

Figure S17: A comparison between $H_2$ datasets estimated using population-specific (YRI; Yoruba from Ibadan, Nigeria) recombination maps inferred with LD patterns (pyrho and Omni) and the Bhérer and Zhou datasets used in this study. 95% bootstrap CI are shown for the Bhérer dataset.

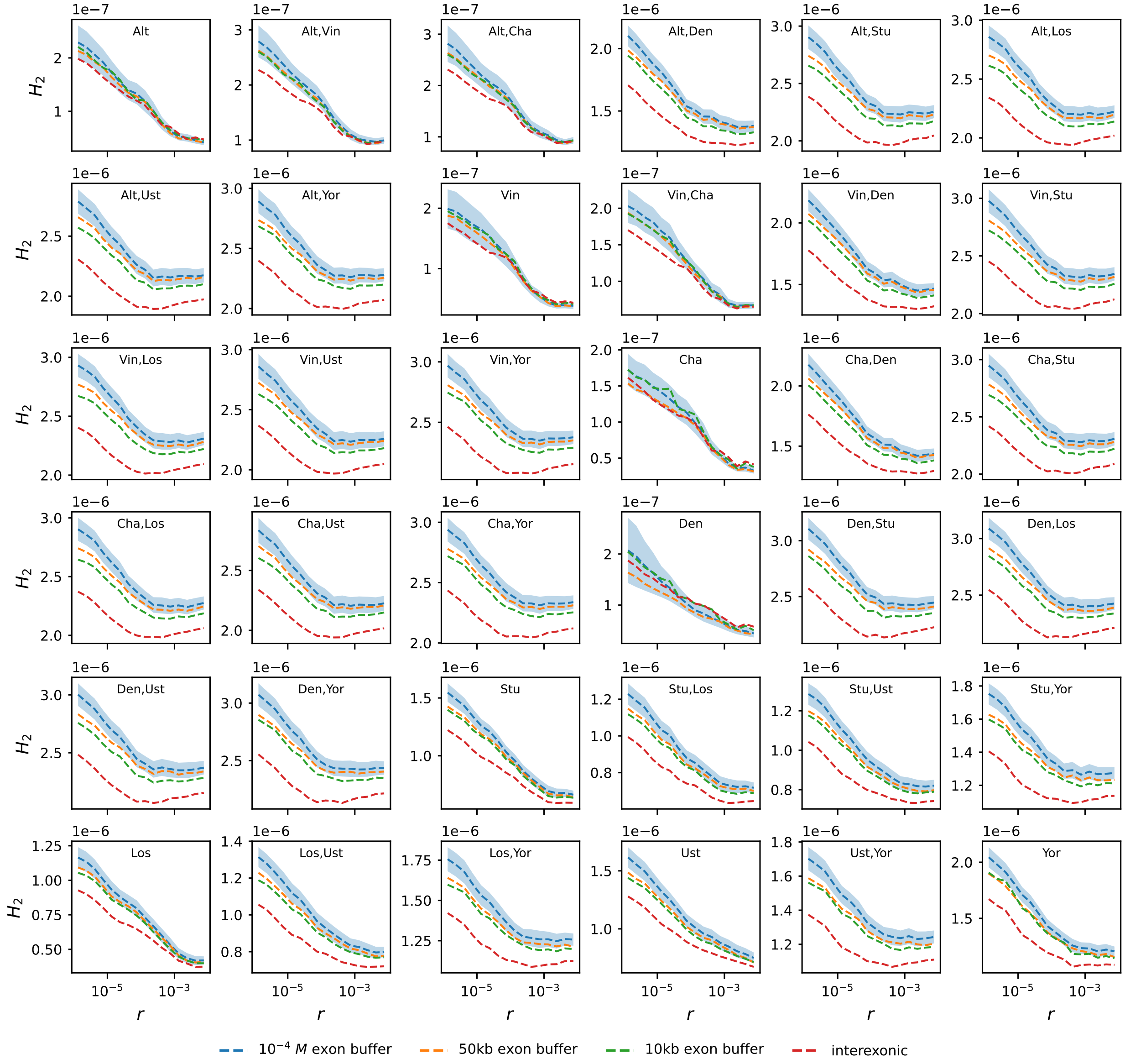

Figure S18: A comparison of $H_2$ estimated with four different genetic masks, defined by the length of the minimum distance (in physical or map units) imposed about exonic sites. 95% bootstrap CI are shown for $10^{-4}$ $M$ exon buffer estimates, which we use in all analyses.

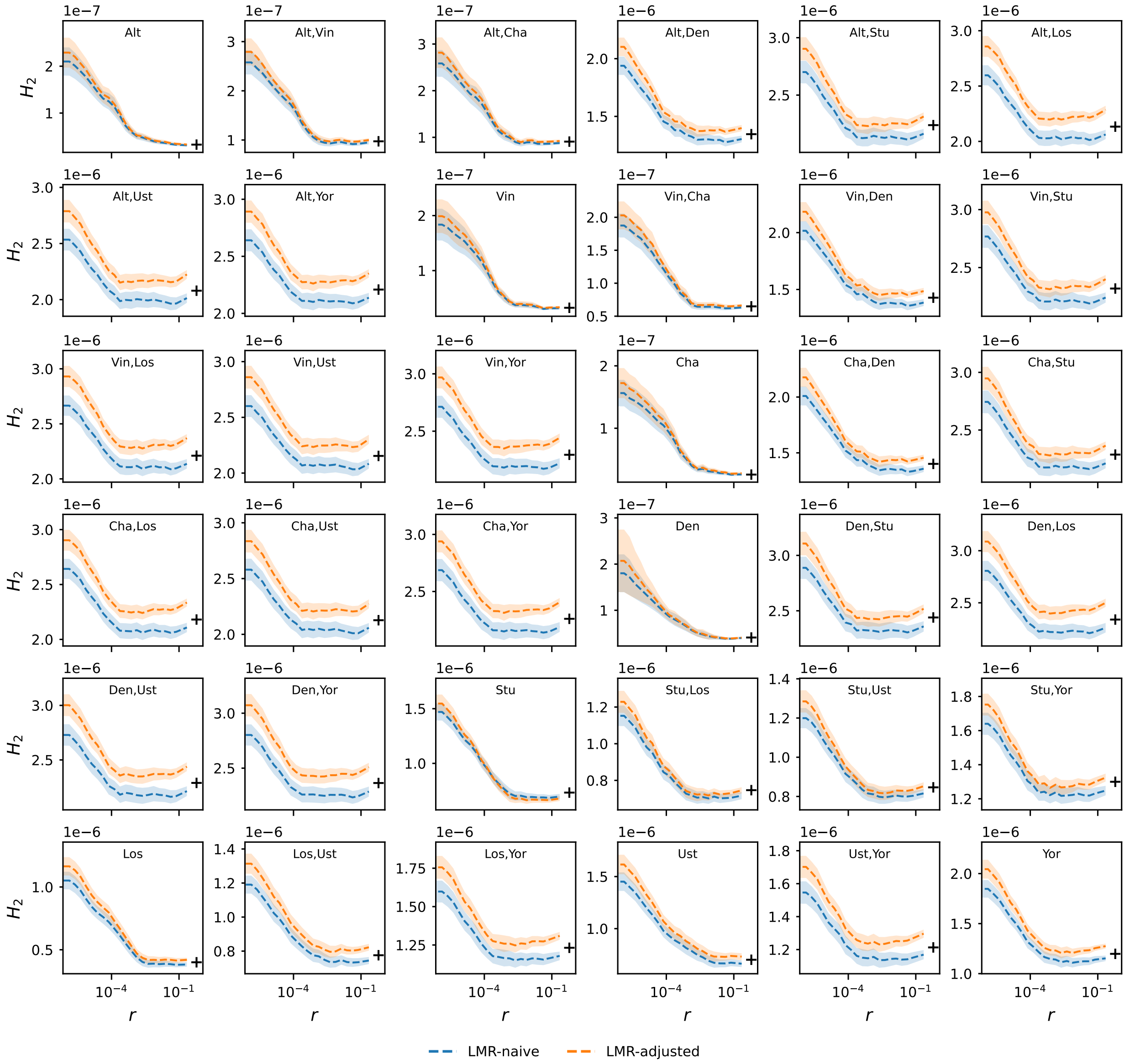

Figure S19: $H_2$ across the complete domain of recombination distances $[0, 0.5)$, estimated using local mutation rate (LMR)-naive and adjusted statistics (Bhérer dataset). Black crosses mark genome-wide the square of the genome-wide heterozygosity $H^2$, which is expected to equal $H_2$ at long distances.

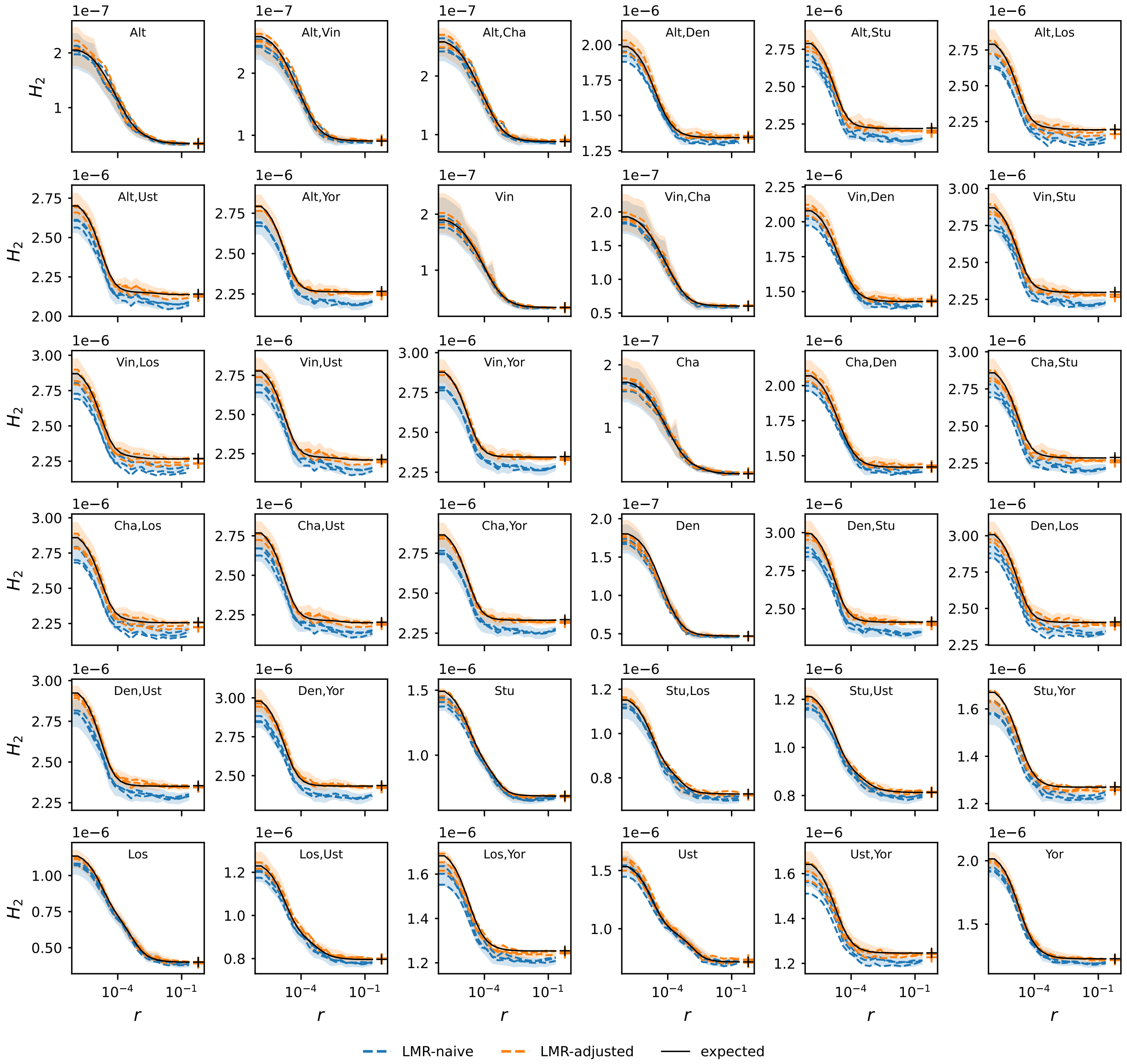

Figure S20: A comparison of three replicate simulated datasets, produced under a model with local mutation rate (LMR) variation. For each replicate, we show the LMR-naive and LMR-adjusted statistics. 95% bootstrap CI are shown for the first replicate. Crosses at panel right show the observed or expected value of the square of genome-wide heterozygosity, $H^2$.

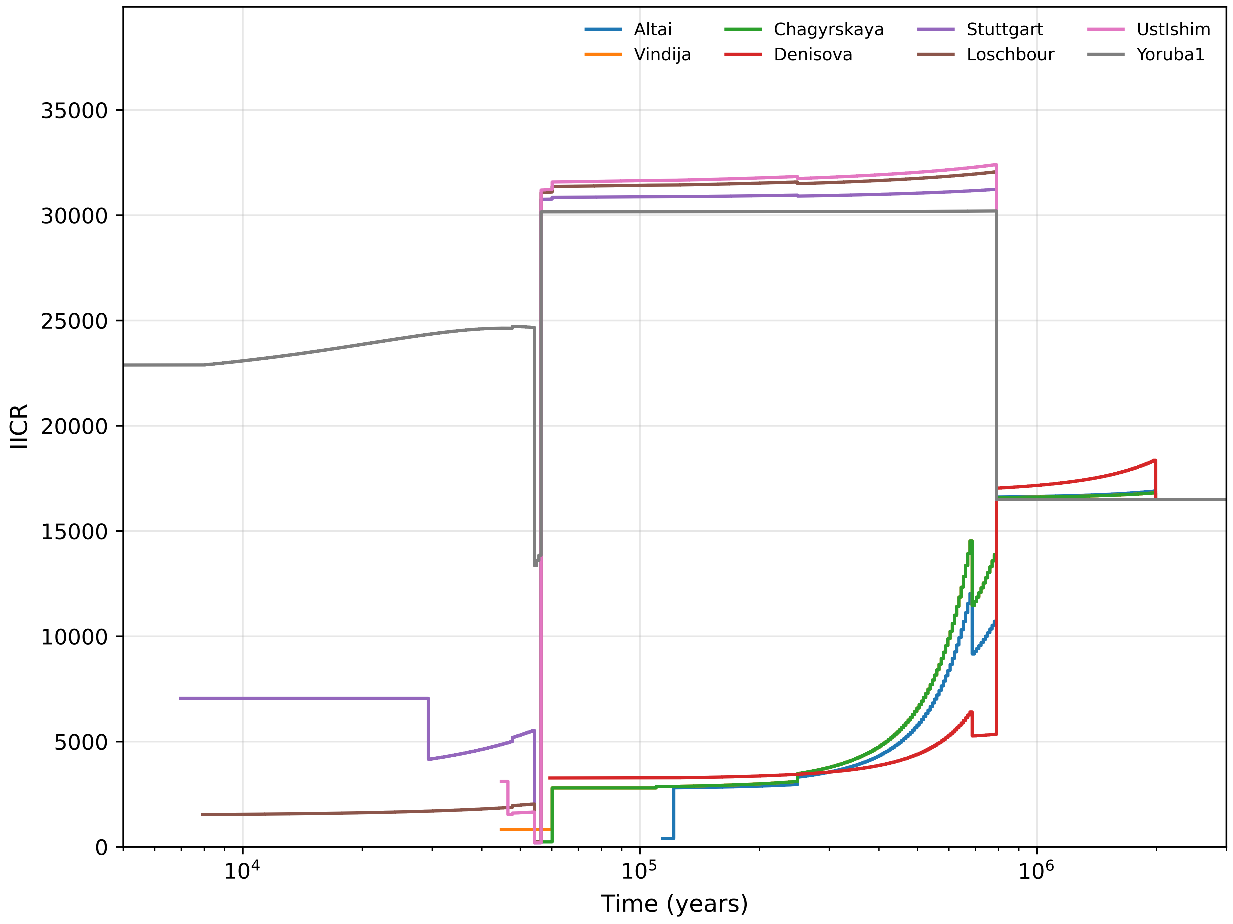

Figure S21: Inverse instantaneous coalescence rates (IICR) predicted for the full model (Bhérer dataset).

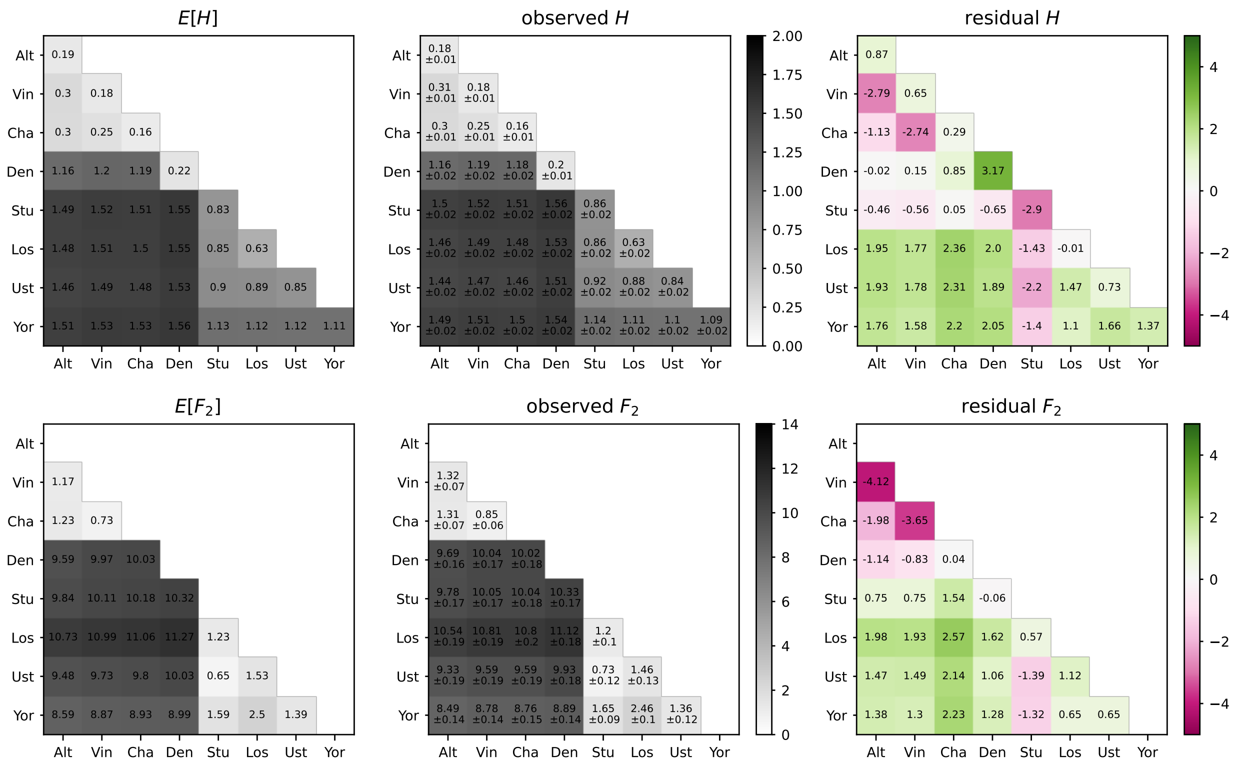

Figure S22: A comparison between predicted and empirical $H$ and $F_2$ statistics for the full model (Bhérer dataset). All $H$ statistics are scaled by $10^3$ and all $F_2$ statistics are scaled by $10^4$. Colormaps are shared between expected/observed panels for each statistic. Residuals are calculated as $(\text{Model}_{i,j} - \text{Data}_{i,j})/\sqrt{V_{i,j}}$, where $V_{i,j}$ is the bootstrap variance observed for statistic $i$ in bin $j$.

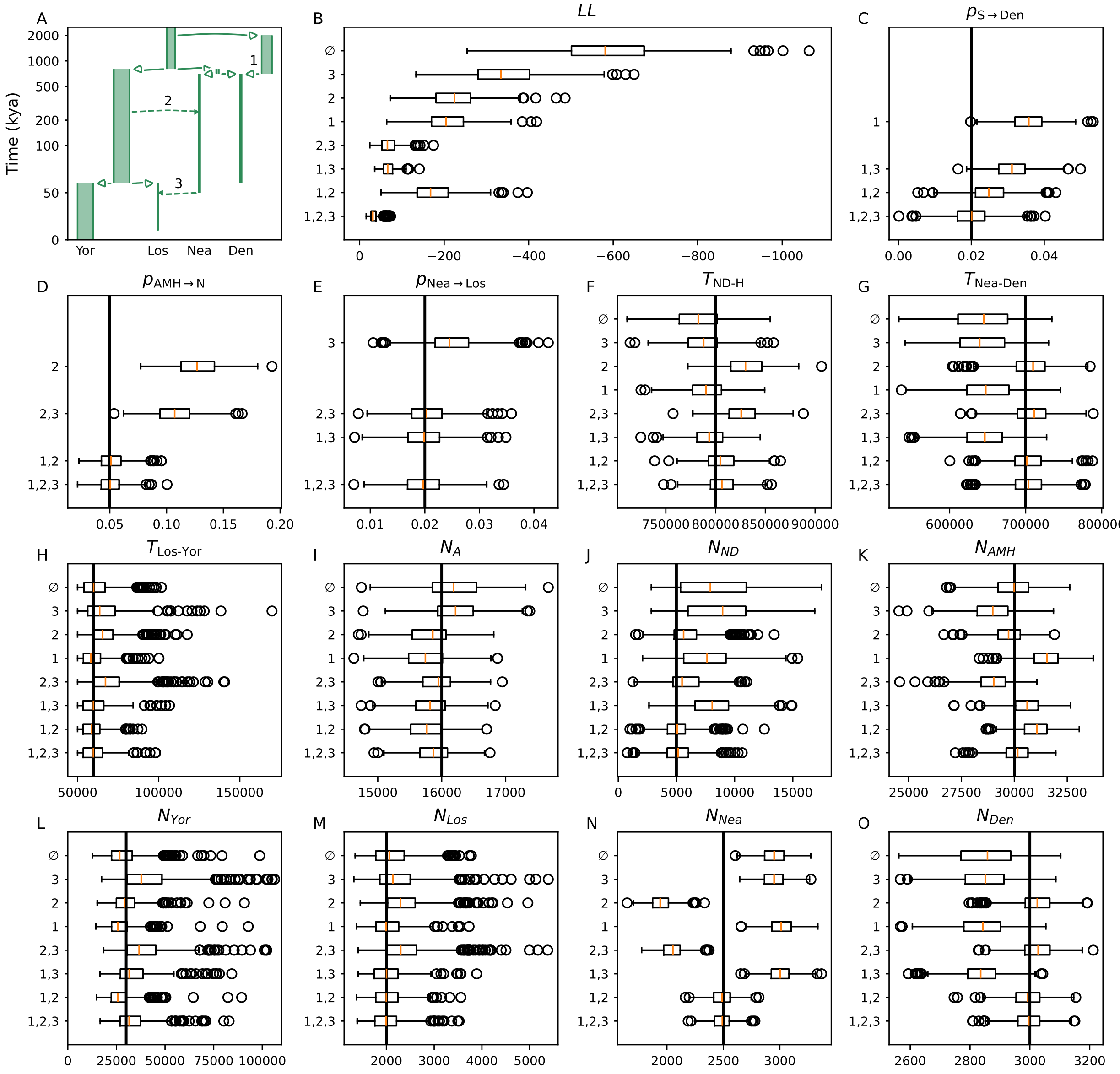

Figure S23: Parameter estimates are biased under misspecified models (500 replicate simulations). (A) The underlying model has three instantaneous gene flow events, labeled 1, 2, and 3, with respective proportions 2%, 5%, 2%. (B) Summary of log-likelihoods under misspecified and true models. Models are named by the introgression pulses that they incorporate (e.g., model 1,2,3 corresponds to the ground truth with all three pulses, while model 3 incorporates only the Neanderthal-to-Loschbour introgression and model ∅ has no gene flow). (C-O) Distributions of MLE parameters under each model. True parameter values are marked by vertical lines.
